# Slab-PTM: A Coarse-Grained Force-Field Parameter Patch for Modeling Post-Translational Modification Effects on Biomolecular Condensates

**DOI:** 10.1101/2025.11.24.690320

**Authors:** Junxi Mu, Luhua Lai

## Abstract

Intrinsically disordered proteins (IDPs) play crucial roles in biomolecular condensate formation. While molecular dynamics simulations employing coarse-grained models have emerged as useful tools for studying IDP phase behavior, current force-field parameterizations remain limited in their ability to simulate post-translational modifications (PTMs), which are the critical regulatory elements of IDP involved condensation with profound biological implications.

To address this gap, we developed interaction parameters for five common PTM types: phosphorylated serine (pSer), threonine (pThr), and tyrosine (pTyr); acetylated lysine (AcLys); and asymmetric dimethylarginine (aDMA). Using all-atom umbrella sampling simulations, we computed residue-specific potentials of mean force (PMFs) between modified and canonical amino acids. These PMF-derived parameters were systematically integrated into the established slab-geometry coarse-grained models (CALVADOS and Mpipi) via an additive module termed as Slab-PTM. Benchmark simulations demonstrate that Slab-PTM accurately captures the effects of PTMs on IDP phase behavior while remaining fully compatible with existing LLPS simulation frameworks. In addition, Slab-PTM enables the identification of molecular grammar elements through which PTMs modulate IDP-driven phase separation.

## Introduction

Biomolecular condensates formed through protein liquid-liquid phase separation (LLPS) are essential for regulating various biological processes^1^, such as cellular development^2^, environmental stress response^3^, transcriptional control^4^, DNA damage repair^5^, RNA processing and transport^6^, signal transduction^7^, innate immune functions^8^, and neuronal communication^9^. These condensates, which typically lack enclosing membranes, often form due to multivalent interactions among intrinsically disordered proteins (IDPs)^10^. It has been shown that the specific sequence characteristics of IDPs substantially affect the physicochemical properties and behaviors of these condensates^11,12^.

Recent studies have further highlighted the importance of post-translational modifications (PTMs) in modulating the phase behavior of IDP condensates, thereby regulating various biological functions^13,14^. For instance, Ding et al. revealed that site-specific phosphorylation at Ser61 disrupts intra- and intermolecular interactions, providing crucial insights into preventing the pathological aggregation of FUS proteins^15^. Similarly, Nott et al. demonstrated that arginine methylation destabilizes Ddx4 droplets, significantly affecting LLPS both in vitro and in cellular contexts^16^. Additionally, Saito et al. reported that HDAC6-mediated deacetylation of the N-terminal IDR1 domain of DDX3X enhances its LLPS, facilitating stress granule maturation^17^.

Molecular dynamics (MD) simulations have been widely applied in elucidating the molecular interactions and kinetic mechanisms underlying LLPS^18^. While all-atom simulations provide detailed insights, they remain computationally intensive for IDP condensate systems^19,20^. Consequently, various coarse-grained force fields have been developed to effectively simulate these systems^21^. For instance, Pappu et al. introduced lattice-based methods^21^, whereas Mittal et al. developed the widely adopted hydrophobicity-scale (HPS) model implemented within a slab geometry simulation framework^22^. Subsequent models, such as the CALVADOS force field by Lindorff et al.^23-25^ and the Mpipi force field^26^, have similarly employed slab geometries to efficiently simulate IDP LLPS phenomena.

Nevertheless, current coarse-grained models generally lack comprehensive parameterization for residues undergoing post-translational modifications. Although certain studies have partially addressed this gap by providing parameters for specific PTM residues^27,28^, these parameters remain limited and are typically not transferable across different force fields. To address this limitation, we developed a generally applicable parameter set, Slab-PTM, which can be integrated into prominent HPS-based coarse-grained models, such as CALVADOS2^23^ and Mpipi^26^. The Slab-PTM parameters include five critical PTM residues: phosphorylated serine (pSer), threonine (pThr), tyrosine (pTyr), acetylated lysine (AcK), and methylated arginine (meArg). These residues collectively encompass the majority of PTM scenarios relevant to IDP LLPS, and the Slab-PTM parameters demonstrate excellent agreement with experimental observations.

To validate Slab-PTM, we benchmarked a series of experimentally characterized IDP systems across five major PTM types. Our simulations accurately recapitulate both PTM-induced suppression and enhancement of phase separation, in agreement with experimental observations. These include phosphorylation-modulated systems such as hnRNP A2, TDP-43, FUS, and NS2; acetylation-regulated LLPS in DDX3X; and arginine methylation effects in hnRNP A2 and DDX4. Across all cases, Slab-PTM captures the residue-level interaction changes underlying altered phase behavior and proves transferable across CALVADOS2 and Mpipi force fields. Together, these results establish Slab-PTM as a robust and generalizable framework for modeling PTM-regulated IDP involved phase separation and decoding the molecular grammar of biomolecular condensation.

## Methods and Materials

### Parameterization

Currently, two main series of slab-geometry-based LLPS force fields exist: the Mpipi series^26,29^ and the HPS series^22,23,25,30-32^. In these coarse-grained force fields, each amino acid residue or nucleic acid is represented by a single bead. This bead possesses a corresponding mass, charge (q parameter), molecular diameter (σ), and an energy scale parameter (€ in the Mpipi series and λ in the HPS series). The potential energy of such force fields can be described as follows:

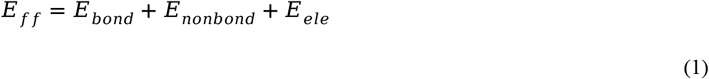

Where *E*_*bond*_ is the harmonic bond potential between two neighbor residues within one protein chain, as follows:

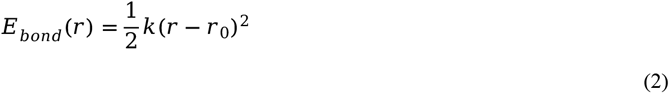

Where *k* is the force constant with value of 8.033 J mol^-1^pm^-2^. *r* is the distance between two residues and *r*_0_ is the equilibrium distance of 380 pm.

Both Mpipi and HPS series of force fields use the Debye-Hückel (DH) potential to describe the electrostatic interaction, as follows:

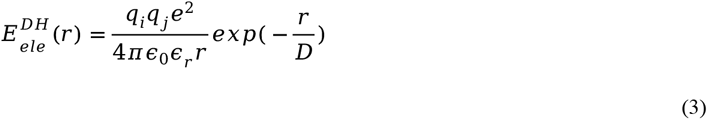

Where q is the average amino acid charge number, e is the elementary charge, €_0_ is the dielectric constant, and r is the distance between two residues.

Here €_r_ denotes the temperature-dependent dielectric constant, which is calculated using the following empirical relationship^33^:

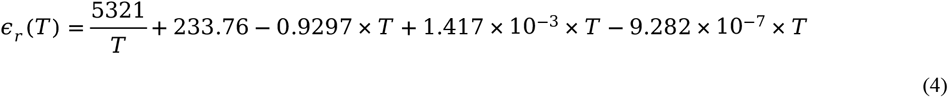

Here *D* is the Debye length calculated as follows:

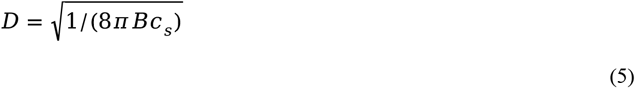

Where c_*s*_ is the ionic strength, and B is the Bjerrum length, as follows:

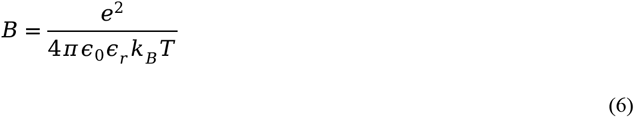

Where €_r_ represents the temperature-dependent dielectric constant and €_0_ represents the dielectric constant of free space.

And k_B_ denotes the Boltzmann constant.

As for *E*_*nonbond*_, the Mpipi force field use the Wang*-*Frenkel (WF) potential^34^ while HPS force fields use the Ashbaugh-Hatch (AH) potential^35^ to describe, both are the variants of the traditional Lennard-Jones (LJ) potential.

The AH potential^35^ is defined as follows:

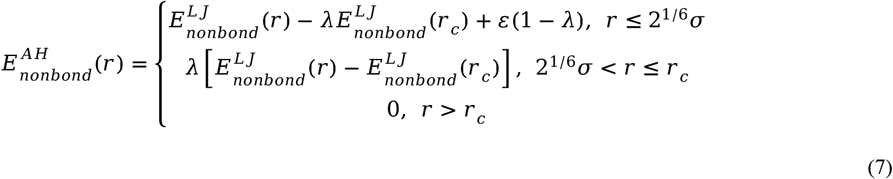

Where ε is the constant parameter of 836.8 J mol-1, r_c_ is 2 nm. *E*^*L]*^ is the LJ potential as follows:

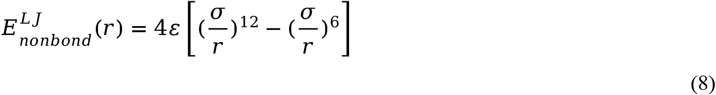

While the WF potential^34^ could be seen as follows:

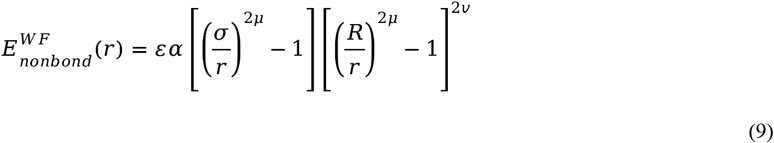

Where *ε* is the adjustable parameter, unlike that of AH potential^35^. *r* is the distance between two residues. Here *R =* 3 *σ* and *v =*1. The *α* could be calculated as follows:

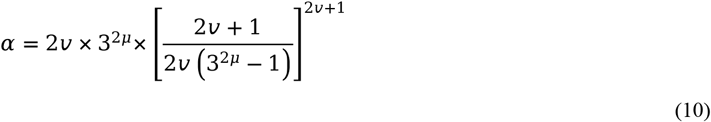

Here *µ* is another adjustable parameter, which typically has a value of 2 for most residue pairs, following the parameters used in Mpipi model. Detailed parameter values are available at https://github.com/decodermu/Slab-PTM.

#### Non-bond fixable energy form

For most HPS force fields, the non-bond interaction parameters (a and 1) for residue pair interactions are typically derived from residue-specific properties using combination rules. Specifically:

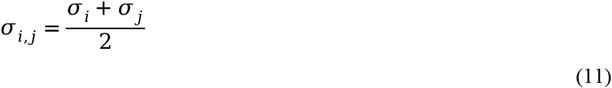

Here the *σ*_*i, j*_ is the parameter used to calculate the non-bond interaction between residue *i* and *j* during simulation. *σ*_*i*_ and *σ*_*j*_ are the molecular diameter parameter of residue *i* and *j*.

The λ parameter was also calculated during simulation, as follows:

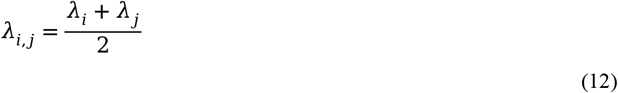

Here the λ_*i, j*_ is the parameter used to calculate the non-bond interaction between residue *i* and *j* during simulation. λ _*i*_ and λ _*j*_ are the energy scale parameter of residue *i* and *j*.

However, this implicit pairwise parameterization scheme inherently couples the interactions of distinct pairs. Specifically, defining the parameters for pairs i,j and j, k automatically constrains the parameters for pair i, k. This emergent behavior limits the ability to independently control and fine-tune specific pairwise interactions.

To overcome this limitation and enable direct, independent control over pairwise interactions, we adopt a direct pair-level parameterization strategy in this work. Instead of deriving *σ*_*i, j*_ and λ_*i, j*_ from residue-level values, we explicitly parametrize the pair-specific parameters *σ*_*i, j*_ and λ_*i, j*_ themselves.

As for Mpipi force field, the W-F potential is already non-bond fixable, here we kept its energy form.

#### Non-Bonded Interactions

To ensure post-modification parameters could be applied across as many existing force fields as possible, we opted for a relatively in-situ parameter patching method rather than developing an entirely new force field. Specifically, we primarily scaled the non-bonded interactions of the post-modification residues, using their native counterparts as references. For example, phosphorylated residues (pSer, pThr, pTyr) were referenced against their unphosphorylated forms (Ser, Thr, Tyr, respectively). Similarly, methylated arginine used native arginine as a reference, while acetylated lysine was referenced against glutamine (Q). The detailed relationships are illustrated in Figure 1 and further elaborated in Table S1.

**Figure 1.**
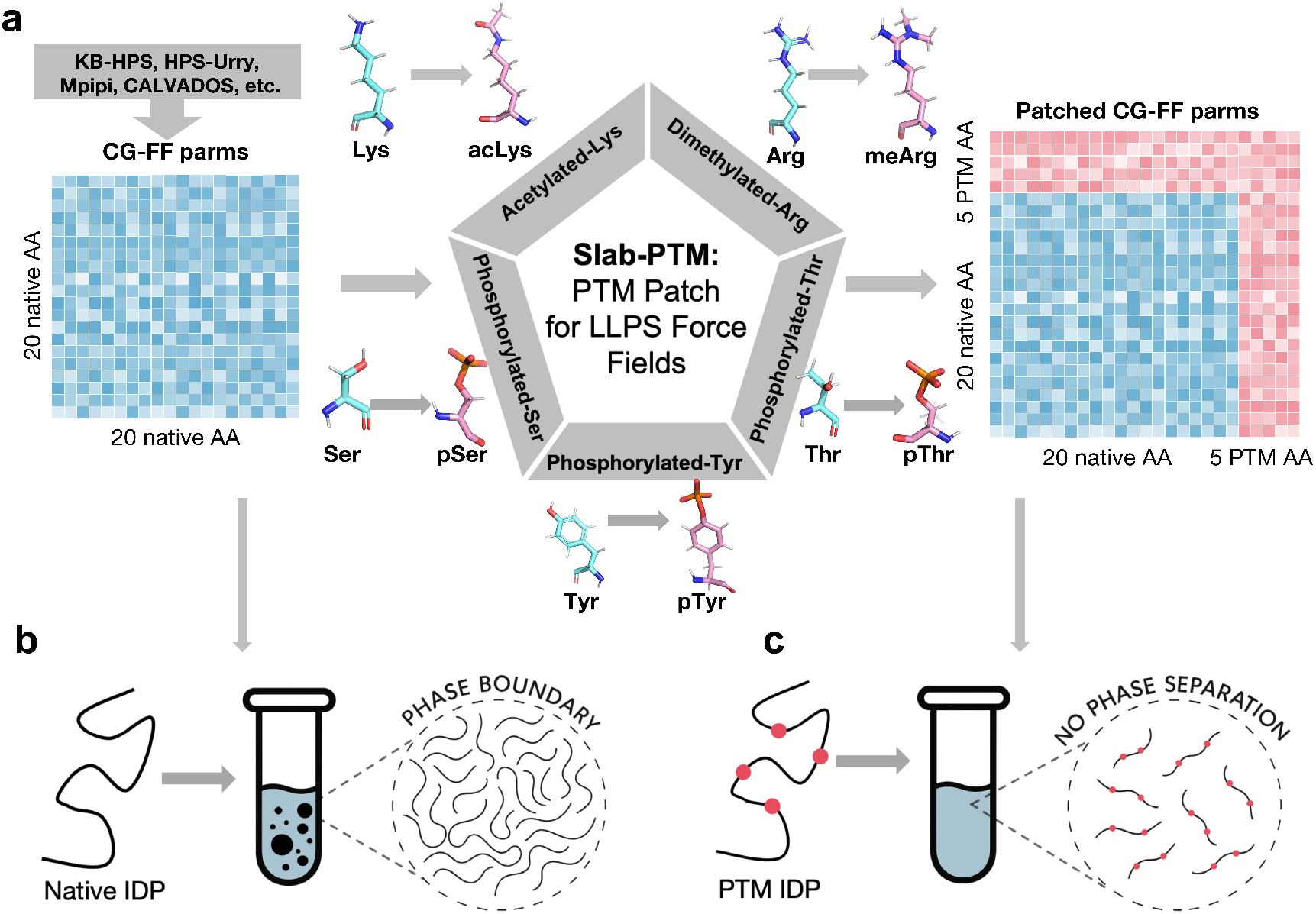
The overall working flow of Slab-PTM. **(a)** Building on the previous coarse-grained force field for phase separation, Slab-PTM provides the interaction parameters between the five PTM residues and the remaining 20 standard amino acids, as well as the interaction parameters among the five PTM residues themselves. **(b-c)** The Slab-PTM-patched force field enables the simulation of phase behavior alterations in IDPs resulting from post-translational modifications.

To address the differing potential energy function forms of the WF and AH potentials, we developed two distinct parameter scaling strategies.

For the WF potential, we modified the σ and *ε* parameters, which describe the position and depth of the potential energy well, respectively. The detailed parameter modifications are as follows:

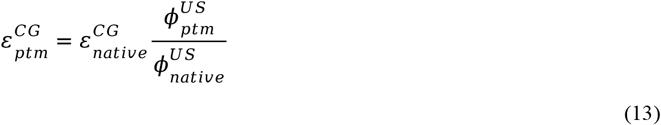

Where 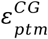 is the modified *σ* parameter of PTM residue in the coarse-grained force field, and 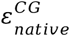 is that of the reference native residue. 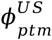 represents the energy trap depth of the PTM residue pair pmf curve sampled from umbrella sampling, while 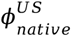 is that of the reference residue pair.

For the *σ* parameter of WF potential^34^, the modification could be seen as follows:

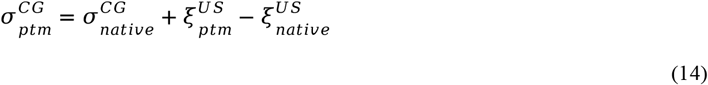

Where 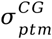 is the modified *σ* parameter of PTM residue in the coarse-grained force field, 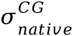 is that of the reference native residue.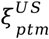 represents the energy trap position of the PTM residue pair pmf curve sampled from umbrella sampling, while 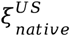 is that of the reference residue pair.

The AH potential^35^, unlike the WF potential, employs a constant *ε* value and instead utilizes the λ parameter to control the non-bonded interaction strength. The modification approach for σ is similar to that of the WF potential, as presented in Eq. 2. However, λ has a range between 0 and 1 and is linearly scaled, in contrast to the multiplicative scaling applied to the *ε* parameter. Consequently, we employed the following formula to scale the *1* parameter:

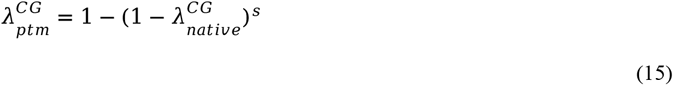

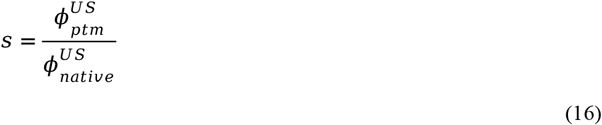

Where 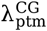 is the modified λ parameter of PTM residue in the coarse-grained force field, and 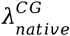 is that of the reference native residue. 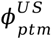 represents the energy trap depth of the PTM residue pair pmf curve sampled from umbrella sampling, while 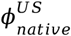 is that of the reference residue pair. See in Figure S1, this scaling strategy achieves the following rules:

1. The modified parameter has a range between 0 and 1.
2. When 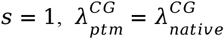.
3. The modified parameter increases monotonically with *s* (since *s* is always positive):

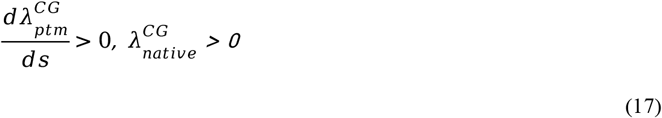
4. When the parameter is significantly less than 1, it can be approximated as being scaled multiplicatively by *s*:

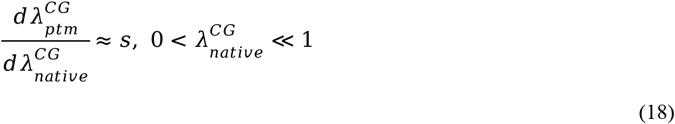
5. When the parameter is close to 1, the partial derivative of the function with respect to 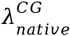 approaches 0:

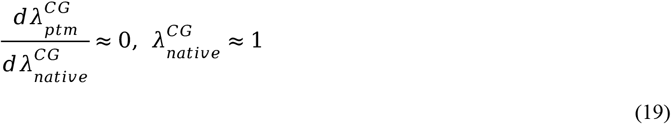

For the σ parameter, the scaling strategy employed is identical to that used for the WF potential.

#### Electrostatic interaction

The charge of residues may be changed completely by post-translational modification. For phosphorylated residues, the average charge was estimated using the Henderson-Hasselbalch equation. The pKa values employed in this calculation are presented in Table S2. For these phosphorylated residues, the average charge can be calculated as:

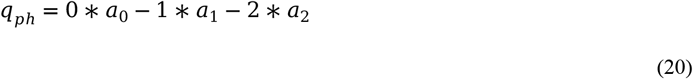

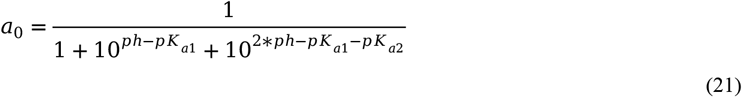

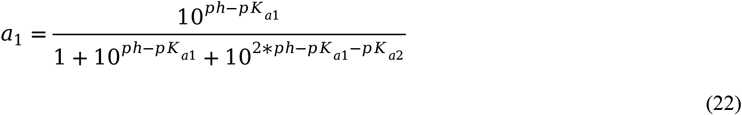

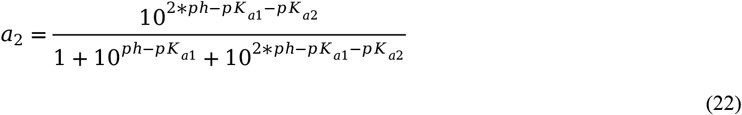

Where *q* _*ph*_ represents the average charge at the corresponding pH condition *a*_0,_ *a*_1_ and *a*_2_ denote the percentages of different protonation states.

Meanwhile, the average charge of methylated arginine is set to +1, while acetylated lysine is considered neutral.

### Umbrella Sampling

To simulate the interaction between two residues in IDPs, we employed amino acid dimers. Each residue’s N- and C-terminal ends were capped with acetyl and N-methyl groups, respectively. The amino acid pairs were oriented with their side chains facing each other. The dimer system was solvated within a water box with an edge thickness of 1.5 nm. Sodium (Na^+^) and chloride (Cl^-^) ions were added to achieve a salt concentration of approximately 150 mM and to ensure charge neutrality. The amino acid dimers were modeled by the CHARMM-36 force field^36^ and its cation-pi refitted patch^37^. The solvent molecules were modeled by TIP3P^38^.

For each system, initial energy minimization was performed using the steepest descent algorithm, with a force tolerance of 1000 J mol^−1^pm^−1^. Bond lengths involving hydrogen atoms were constrained using the LINCS algorithm^39^. Van der Waals interactions were smoothly switched off between 1.0 nm and 1.2 nm using a force-switch modifier, while electrostatic interactions were calculated using the Particle Mesh Ewald (PME) method^40^ with a direct-space cutoff of 1.2 nm. Position restraints were applied as defined in the system topology, with a strength of 100 J mol^−1^ pm^−2^.

For production runs, positional restraints of 1 J mol^−1^ pm^−2^ were applied to constrain heavy atoms in directions perpendicular to the pulling direction. The center-of-mass (COM) distance between interacting pairs was restrained with a harmonic umbrella potential, using a pulling force constant of 6 J mol^−1^ pm^−2^.

For each system, we conducted simulations across 33 windows, ranging from 0.35 nm to 2 nm, with a step width of 0.05 nm. For each window, three parallel trajectories were run for 10 ns. Umbrella sampling data were analyzed using the Weighted Histogram Analysis Method (WHAM)^41,42^. The first 2 ns of each simulation was allocated for equilibration and excluded from the WHAM analysis. All atomistic simulations and analyses were carried out using the GROMACS simulation package.

### Benchmark

The simulation codes and parameters are available at https://github.com/decodermu/Slab-PTM. We performed LLPS simulations for experimentally validated IDP systems. By comparing the simulated condensate properties of PTM IDPs with those of native IDPs, we aim to benchmark the performance of Slab-PTM for PTM LLPS systems. The simulated systems and their corresponding experimental references are listed in Table S1.

For benchmarking purposes, we selected two force fields: Mpipi^26^ and CALVADOS2^23^. These represent widely used models from the Mpipi series and HPS series, respectively. We have compiled a collection of IDPs reported in the literature for which experimental phase separation data are available for both their native and post-modified states. Detailed IDP systems and reference information can be found in Table S3.

The IDP chains were aligned along the z-axis, with their middle beads randomly placed in the xy-plane at positions separated by more than 0.7 nm. For each system, simulations were performed across a temperature gradient. The detailed temperature profiles for each system are provided in Table S3. Each trajectory ran for 1000 ns, with the initial 400 ns allocated for equilibration and excluded from subsequent analysis.

During density analysis, the slab was centered in the simulation box at each frame, consistent with previous studies^23,25^. The protein densities of both the dilute and dense phases were estimated by fitting the semi-profiles in the z*>* 0 and z*<* 0 regions to the following equation:

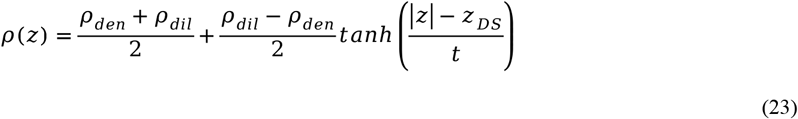

Where *ρ* (*z*) is the equilibrium density profile, calculated by averaging over the trajectory of the system at equilibrium. *ρ* _*den*_and *ρ* _*dil*_are the protein densities of the dense phase and dilute phase, respectively. z means the position along the z-axis of the simulation box. z_*DS*_ means the position of the dividing surface, and *t* is the thickness of the interfacial region.

The critical temperature could be estimated by the *ρ* _*den*_ and *ρ* _*dil*_ values at different temperatures, by fitting the following:

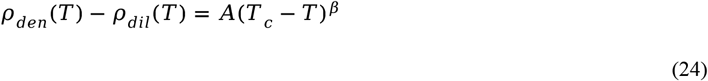

Where *T*_*c*_ is the fitted critical temperature and *A* is another fitted parameter, while *β* is the critical exponent which is set to 0.325 (universality class of 3D Ising model).

Then the critical density could be estimated by fitting the following:

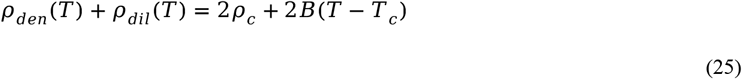

Where *ρ* _*c*_ is the fitted critical density and *B* is another fitted parameter.

To better capture and visualize the detailed influence of PTM residues, we quantified the changes in sequence-level interaction probabilities. For each residue *i*, we quantified the net effect of PTM by summing the absolute changes in interaction probabilities across all partners:

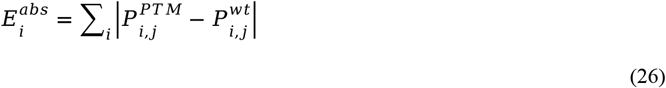

Where 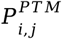 and 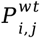 are the averaged interaction probability during the simulation between residue *i* and *j*.

To facilitate cross-residue comparison, 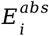 values were further standardized using Z-score normalization:

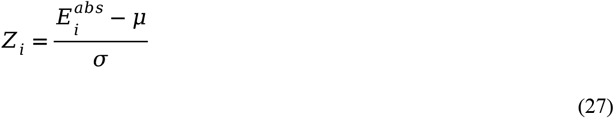

Where *µ* and σ denote the mean and standard deviation of 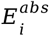 across all residues. When σ = 0, *Z*_*i*_ all values were set to zero to avoid division by zero.

## Results

### Development of Slab-PTM

As detailed in the Methods section, we developed Slab-PTM, a PTM parameter patch for coarse-grained force fields used in simulating IDP LLPS systems. As shown in Figure 1a, Slab-PTM can load parameters for five types of PTM residues onto a given base LLPS force field, enabling it to handle IDP LLPS systems containing PTM-modified residues. Furthermore, it allows the simulation of phase separation changes in IDPs upon PTM, as demonstrated in Figure 1b and 1c.

### Benchmark on Phosphorylation System

We benchmarked four types of LLPS-related IDPs previously studied for the effects of residue phosphorylation on phase behavior. The detailed simulation settings were given in Table S3. Slab-PTM accurately captured both the enhancing and weakening effects of phosphorylation on phase separation, regardless of whether it was patched into CALVADOS2 or the Mpipi model. Moreover, residue-level interaction probability maps, averaged across multi-chain simulations, revealed how PTMs alter the molecular grammar underlying IDP phase behavior. Contact probability heatmaps generated using both CALVADOS2 and Mpipi showed consistent PTM-induced interaction patterns, demonstrating the transferability of the Slab-PTM patch across force fields.

The low-complexity domain (LCD) of hnRNP A2 has previously been shown to undergo strong suppression of phase separation upon tyrosine phosphorylation (Figure 2a, b)^43^. Our simulations reproduce this effect: the non-phosphorylated A2 LCD forms a condensed phase in the slab box, while the phosphorylated form remains uniformly distributed, failing to condense (Figure 2c, g). Residue-level contact maps (Figure 2d, h and Figure 2e, i) show that phosphorylation disrupts key interaction hotspots, particularly in tyrosine-rich regions, introducing repulsion and weakening multivalent interactions necessary for LLPS. The interaction probability Z-score (Figure 2f, j) shows that phosphorylation at Y257, Y264, Y271, Y275, Y278, Y283, Y288, Y294, Y301, Y335, and Y341 play important roles in reducing multivalent interactions. Combined with the detailed IDR sequences and contact patterns (Figures S7 and S8), it can be observed that these key phosphorylation sites are all located within a π– π interaction-rich region. Therefore, it may be concluded that in the non-phosphorylated state, LLPS is driven primarily by π-π interactions between F and Y residues and by cation-n interactions involving a few R residues. Phosphorylation of Y residues reduce these interactions and adds electrostatic repulsion. Detailed protein-density plots are available for hnRNP A2 LCD: the native protein in Figure S21 (patched CALVADOS2) and Figure S22 (patched Mpipi), and the phosphorylated protein in Figure S25 (patched CALVADOS2) and Figure S26 (patched Mpipi).

**Figure 2.**
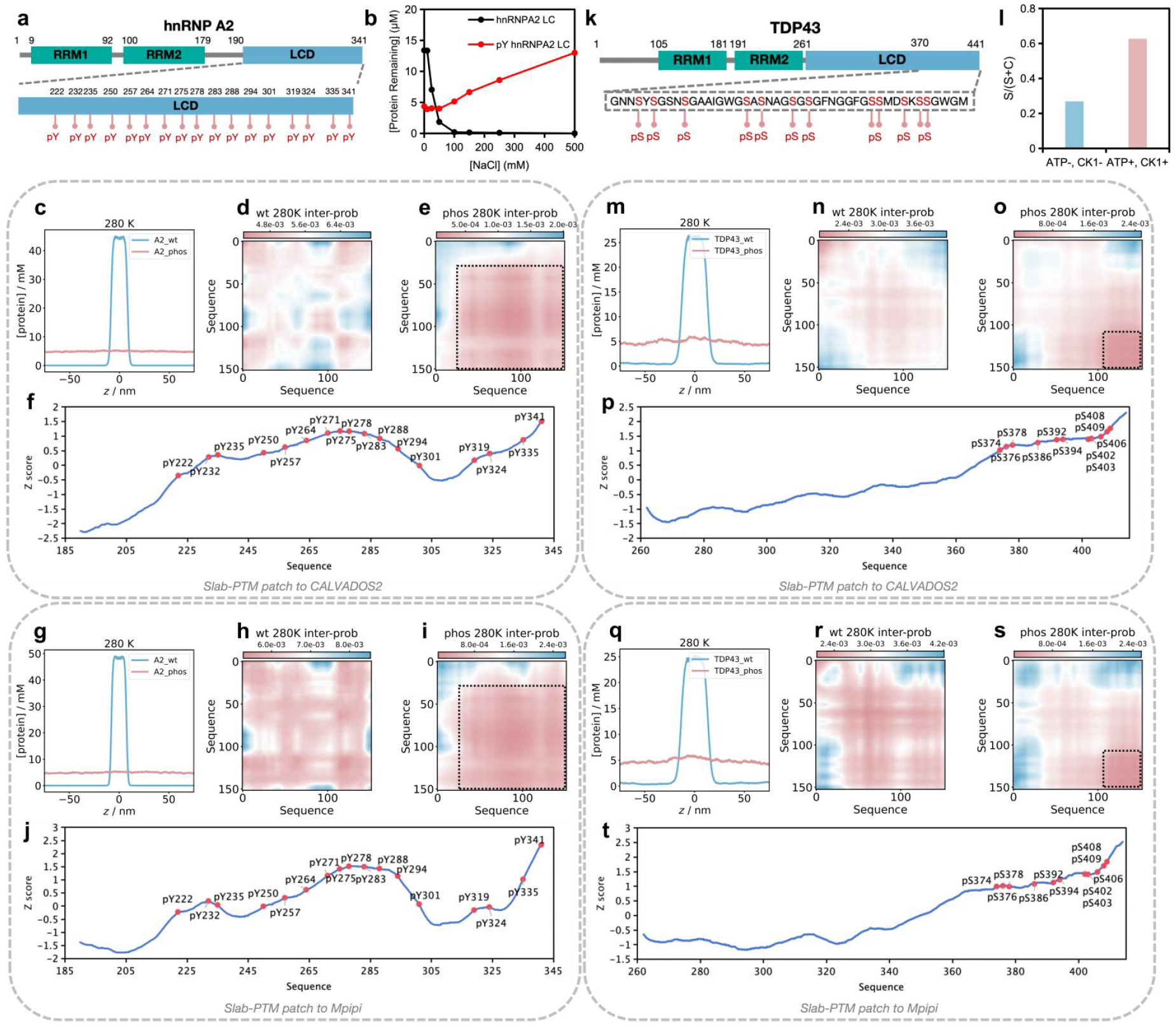
Benchmark on phosphorylated hnRNP A2 LCD and TDP-43 LCD. **(a)** Sequence of the low-complexity domain (LCD) of hnRNP A2 with phosphorylation sites indicated. **(b)** Experimental results comparing hnRNP A2 LCD and phosphorylated hnRNP A2 LCD^43^. **(c, g)** Protein density along the z-axis for hnRNP A2 LCD and its phosphorylated form, simulated using Slab-PTM patched CALVADOS2 (c) and Mpipi (g). Blue represents the native IDP and red represents the PTM-modified IDP. **(d, e, h, i)** Residue interaction probability maps for hnRNP A2 LCD (non-PTM: d, h) and phosphorylated hnRNP A2 LCD (e, i), simulated using Slab-PTM patched CALVADOS2 (d, e) and Mpipi. (h,ih). Blue indicates stronger interactions; red indicates weaker interactions. **(f, j)** Interaction probability shuffle Z-scores of amino acid residues, with PTM sites highlighted in red, simulated using Slab-PTM patched CALVADOS2 (f) and Mpipi (j). **(k)** Sequence of the low-complexity domain (LCD) of TDP43 with phosphorylation sites indicated. **(I)** Experimental results comparing TDP43 LCD and phosphorylated TDP43 LCD^44^. **(m, q)** Protein density along the z-axis for TDP43 LCD and its phosphorylated form, simulated using Slab-PTM patched CALVADOS2 (m) and Mpipi (q). Blue represents the native IDP and red represents the PTM-modified IDP. **(n, o, r, s)** Residue interaction probability maps for TDP43 LCD (non-PTM: n, r) and phosphorylated TDP43 LCD (o, s), simulated using Slab-PTM patched CALVADOS2 (n, o) and Mpipi. (r, s). Blue indicates stronger interactions; red indicates weaker interactions. **(p, t)** Interaction probability shuffle Z-scores of amino acid residues, with PTM sites highlighted in red, simulated using Slab-PTM patched CALVADOS2 (o) and Mpipi (t).

A similar effect is observed for the LCD of TDP-43, where phosphorylation at the C-terminal serine residues suppresses LLPS (Figure 2k, l)^44^. In simulations, the non-phosphorylated TDP-43 LCD forms a condensed phase, whereas its phosphorylated form does not (Figure 2m, q). Contact maps (Figure 2n, r and Figure 2o, s) again highlight reduced multivalent interactions due to phosphorylation-induced repulsion. As shown in Figures S9 and S10, unlike hnRNP A2, the primary effect here is electrostatic repulsion, with less disruption of the existing multivalent contacts. The interaction probability Z-score (Figure 2f, j) also shows that the phosphorylation sites all plays essential roles in reducing multivalent interactions. Detailed protein-density plots are available for TDP43 LCD: the native protein in Figure S27 (patched CALVADOS2) and Figure S28 (patched Mpipi), and the phosphorylated protein in Figure S29 (patched CALVADOS2) and Figure S30 (patched Mpipi).

In addition to systems with extensive phosphorylation, we also benchmarked cases involving a few site-specific phosphorylation events. Slab-PTM successfully predicted both the weakening (FUS) and enhancement (NS2) of LLPS resulting from phosphorylation, regardless of whether it was applied to CALVADOS2 or Mpipi.

As shown in Figure 3a, b, the N-terminal domain of FUS contains four low-complexity regions (R1-R4). Phosphorylation at a single serine in R2 has been reported to suppress phase separation^15^. In our simulations (Figure 3c, g), both non-phosphorylated and phosphorylated FUS form condensed phases, but the non-PTM system exhibits higher protein density. CALVADOS2 further shows a lower dilute-phase protein density for phosphorylated FUS. Simulated phase diagrams across temperature gradients reveal that phosphorylated FUS has a lower critical temperature in both CALVADOS2 and Mpipi, indicating reduced LLPS propensity. Residue-level contact maps (Figure 3e, f and Figure 3i, j) show that Ser61, when unmodified, contributes to multivalent interactions, but its phosphorylation disrupts them. Figures S11 and S12 reveal that this phosphorylation site lies in a Y-rich domain, where π-π interactions are weakened upon phosphorylation. Detailed protein-density plots are available for FUS N-terminal domain: the native protein in Figure S31 (patched CALVADOS2) and Figure S32 (patched Mpipi), and the phosphorylated protein in Figure S33 (patched CALVADOS2) and Figure S34 (patched Mpipi).

**Figure 3.**
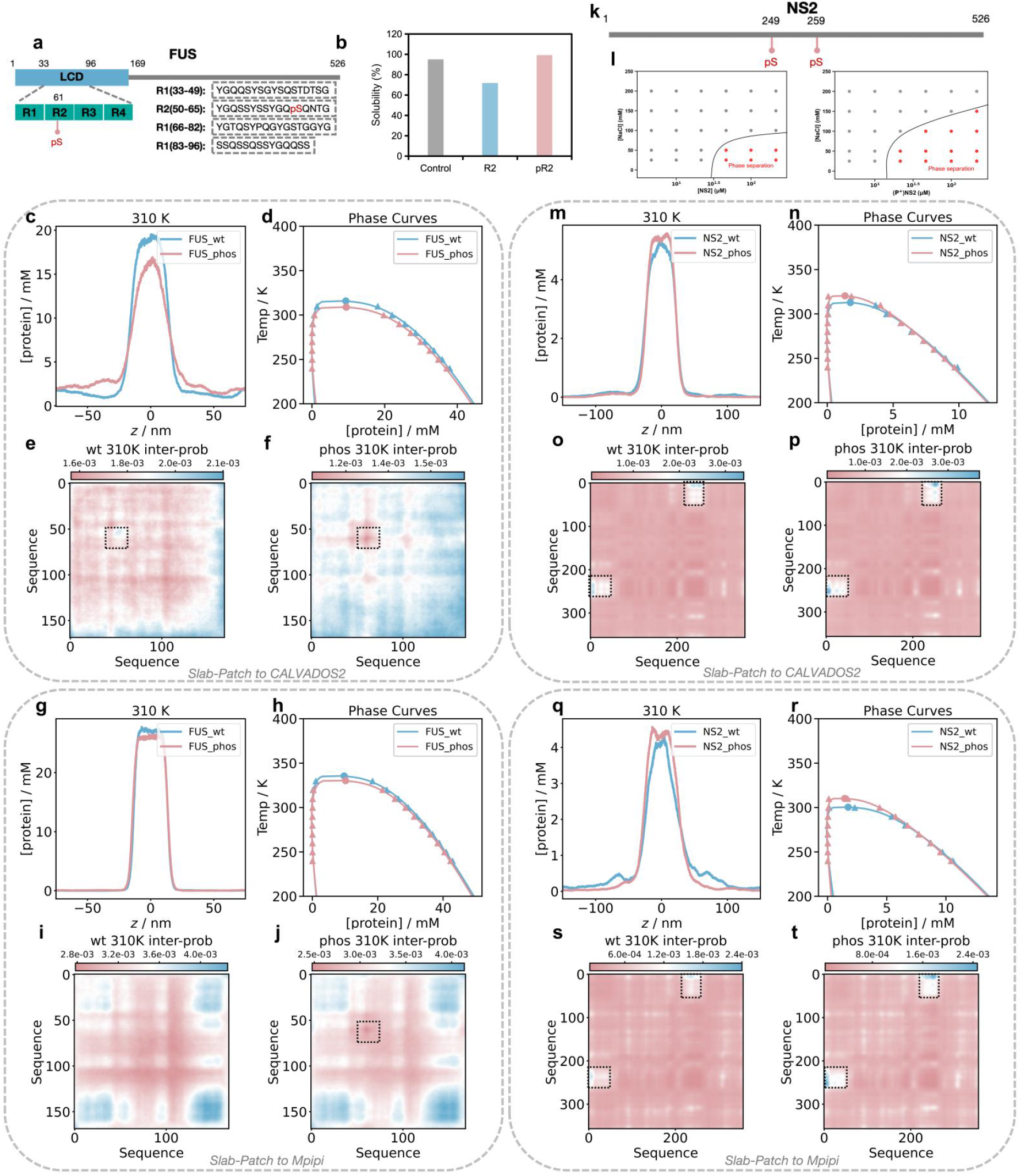
Benchmark on phosphorylated FUS N-terminal domain and NS2. **(a)** Sequence of FUS N-terminal domain with phosphorylation sites indicated. **(b)** Experimental results comparing FUS and phosphorylated FUS^15^. **(c, g)** Protein density along the z-axis for FUS and its phosphorylated form, simulated using Slab-PTM patched CALVADOS2 (c) and Mpipi (g). Blue represents the native IDP and red represents the PTM-modified IDP. (**d, h)** Phase transition curves of non-PTM and PTM systems under a temperature gradient, simulated using Slab-PTM patched CALVADOS2 (d) and Mpipi (h) Blue represents the native IDP and red represents the PTM-modified IDP. **(e, f, i, j)** Residue interaction probability maps for FUS (non-PTM: e, i) and phosphorylated FUS (f, j), simulated using Slab-PTM patched CALVADOS2 (e, f) and Mpipi. (i, j). Blue indicates stronger interactions; red indicates weaker interactions. **(k)** Sequence NS2 with phosphorylation sites indicated. **(l)** Experimental results comparing NS2 and phosphorylated NS2^45^. **(m, q)** Protein density along the z-axis for NS2 and its phosphorylated form, simulated using Slab-PTM patched CALVADOS2 (m) and Mpipi (q). Blue represents the native IDP and red represents the PTM-modified IDP. **(n, r)** Phase transition curves of non-PTM and PTM systems under a temperature gradient, simulated using Slab-PTM patched CALVADOS2 (n) and Mpipi (r) Blue represents the native IDP and red represents the PTM-modified IDP. **(o, p, s, t)** Residue interaction probability maps for NS2 (non-PTM: o, s) and phosphorylated NS2 (p, t), simulated using Slab-PTM patched CALVADOS2 (o, p) and Mpipi. (s, t). Blue indicates stronger interactions; red indicates weaker interactions.

Conversely, phosphorylation can also enhance LLPS, as seen with the NS2 protein. Prior studies report that phosphorylation at Ser249 and Ser259 enhances phase separation (Figure 3k, l)^45^. In our simulations (Figure 3m, q), both non-PTM and phosphorylated NS2 form condensed phases, but the phosphorylated form exhibits higher protein density. Mpipi simulations also indicate lower dilute-phase density in the non-PTM system. Phase diagrams confirm that phosphorylated NS2 has a higher critical temperature in both force fields (Figure 3n, r). Residue-level contact maps (Figure 3o, s and Figure 3p, t) highlight Ser249 and Ser259 as key contributors to multivalent interactions. As detailed in Figures S13 and S14, these sites lie in a negatively charged region that interacts with a positively charged N-terminal segment. Phosphorylation adds further negative charge, strengthening electrostatic interactions and promoting LLPS. Detailed protein-density plots are available for NS2: the native protein in Figure S35 (patched CALVADOS2) and Figure S36 (patched Mpipi), and the phosphorylated protein in Figure S37 (patched CALVADOS2) and Figure S38 (patched Mpipi).

### Benchmark on Acetylation System

We benchmarked lysine acetylation using the N-terminal IDR1 domain of DDX3X. As shown in Figure 4a,b, previous studies identified ten acetylated lysine sites in this region, which strongly suppress LLPS^17^. Our simulations (Figure 4c,f) show that the non-PTM DDX3X forms a condensed phase with high protein density, while the acetylated form fails to condense – only weakly forming a dense phase in Mpipi. Residue-level interaction probability maps (Figure 4d,g and Figure 4e,h) reveal that the non-PTM DDX3X exhibits higher probability contacts compared to its acetylated counterpart. The interaction probability Z-score (Figure 4j) indicates that acetylation of K81 may play a key role in reducing multivalent interactions, while acetylation of K55 and K118 also appears to be important. These findings are consistent with the experimental results shown in Figure 4i^17^. Additional sequence-level contact details are provided in Figures S15 and S16. To further assess the ability of Slab-PTM to capture the influence of single PTM sites on LLPS, we performed additional simulations for all experimentally validated single-site PTM variants. The simulated critical temperature values are shown in Figure 4k. When compared to the experimental turbidity percentage data, the simulations exhibited a relatively good correlation, suggesting that the Slab-PTM patched CALVADOS2 can, to some extent, reproduce the effect of single-site PTMs on multivalent interactions.

**Figure 4.**
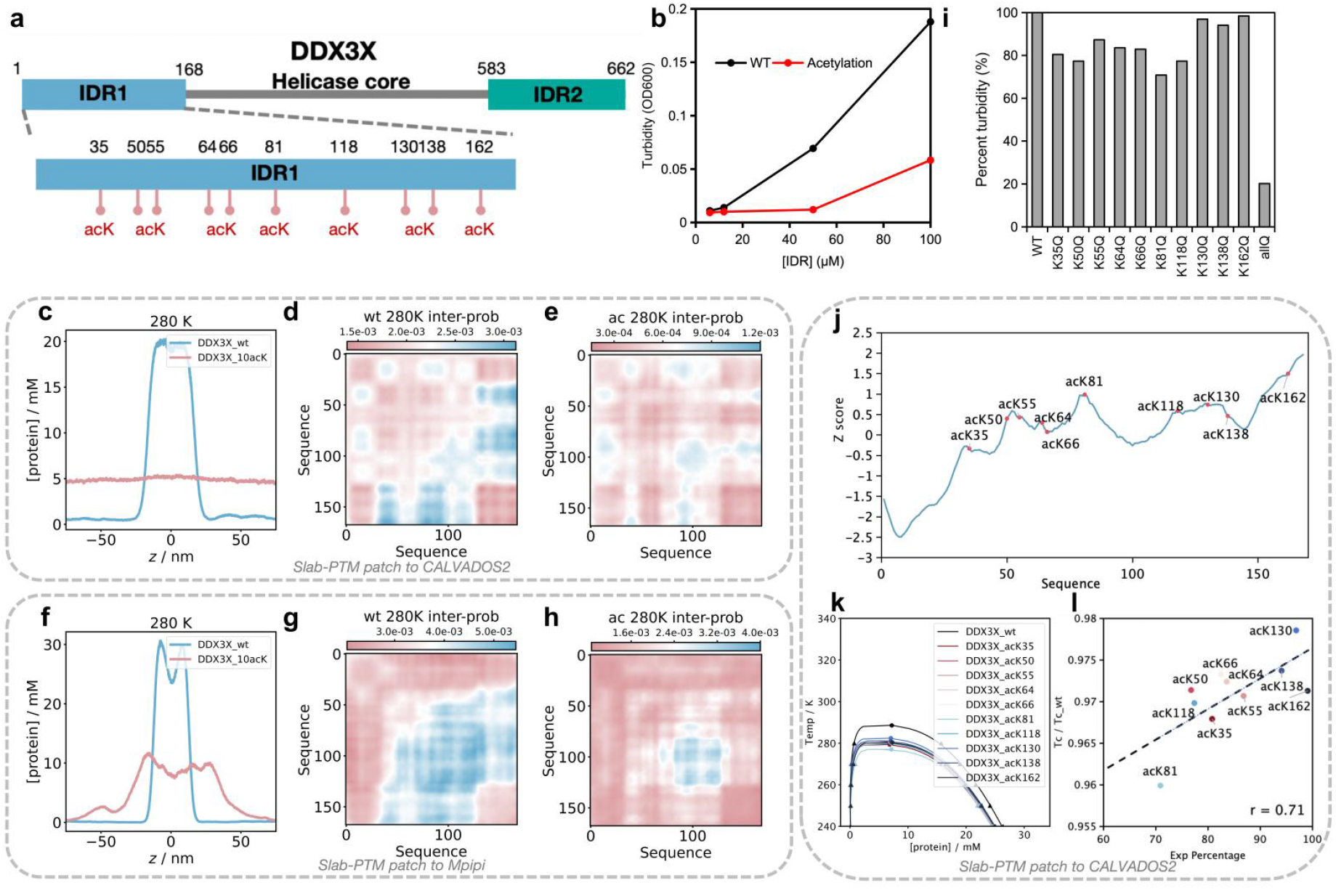
Benchmark on acetylated DDX3X IDR1 domain. **(a)** Sequence of the IDR1 of DDX3X with acetylation sites indicated. **(b)** Experimental results comparing DDX3X IDR1 and acetylated DDX3X IDR1^17^. **(c, f)** Protein density along the z-axis for DDX3X IDR1 and its acetylated form, simulated using Slab-PTM patched CALVADOS2 (c) and Mpipi (f). Blue represents the native IDP and red represents the PTM-modified IDP. **(d, e, g, h)** Residue interaction probability maps for DDX3X IDR1 (non-PTM: d, g) and acetylated DDX3X IDR1 (e, h), simulated using Slab-PTM patched CALVADOS2 (d, e) and Mpipi. (g, h). Blue indicates stronger interactions; red indicates weaker interactions. **(i)** Experimental results comparing DDX3X IDR1 and its single mutation varients^17^. **(j)** Interaction probability shuffle Z-scores of amino acid residues, with PTM sites highlighted in red. **(k)** Simulated critical temperatures (Tc) of DDX3X IDR1 and its single-point mutant variants. **(l)** Correlation between the experimentally measured percent turbidity and the simulated ratio of Tc of DDX3X IDR1 and its single-point mutant variants.

Detailed protein-density plots are available for DDX3X IDR1: the native protein in Figure S39 (patched CALVADOS2) and Figure S40 (patched Mpipi), and the acetylated protein in Figure S41 (patched CALVADOS2) and Figure S42 (patched Mpipi). All the detailed protein-density plots of single-site PTMs could be seen in Figure S47-S56.

### Benchmark on Methylation System

To evaluate the effect of asymmetric dimethylation on arginine residues, we tested two systems–hnRNP A2 and DDX4– both of which demonstrated Slab-PTM’s ability to capture the impact of methylation on LLPS.

As shown in Figure 5a, b, previous studies reported that arginine methylation in the hnRNP A2 LCD significantly suppresses its phase separation^46^. Our simulations confirmed this: the non-PTM hnRNP A2 LCD forms a dense phase in the slab box, whereas the methylated form fails to do so in both CALVADOS2 and Mpipi models (Figure 5c, g). Phase diagrams derived from simulations across temperature gradients show a much lower critical temperature for the methylated variant, indicating reduced LLPS propensity (Figure 5d, h). Residue-level interaction probability maps (Figure 5e, i and Figure 5f, j) reveal minimal contact probability in the methylated regions. Sequence-level analyses (Figures S17 and S18) show that in the non-PTM state, arginines in the N-terminal region contribute cation-n interactions, which are diminished by methylation. In the methylated variant, remaining multivalent contacts primarily originate from arginines in the C-terminal region. Detailed protein-density plots are available for the methylated hnRNP A2 LCD in Figure S23 (patched CALVADOS2) and Figure S24 (patched Mpipi).

**Figure 5.**
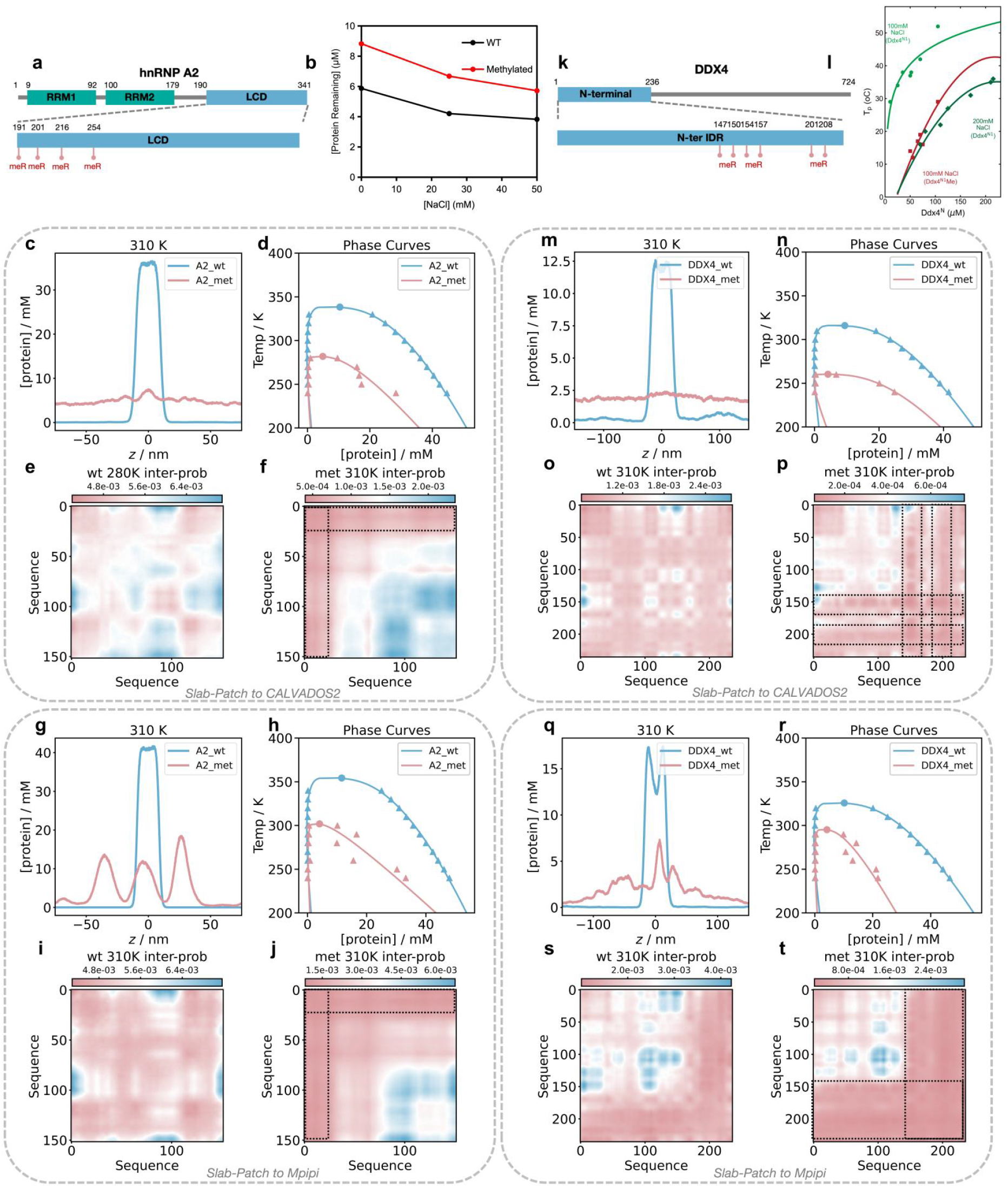
Benchmark on asymmetrically dimethylated hnRNP A2 Low-complex domain (LCD) and DDX4 N-terminal domain. **(a)** Sequence of hnRNP A2 LCD with asymmetric dimethylation sites indicated. **(b)** Experimental results comparing A2 LCD and asymmetrically dimethylated A2 LCD^46^. **(c, g)** Protein density along the z-axis for A2 LCD and its methylated form, simulated using Slab-PTM patched CALVADOS2 (c) and Mpipi (g). Blue represents the native IDP and red represents the PTM-modified IDP. (**d, h)** Phase transition curves of non-PTM and PTM systems under a temperature gradient, simulated using Slab-PTM patched CALVADOS2 (d) and Mpipi (h) Blue represents the native IDP and red represents the PTM-modified IDP. **(e, f, i, j)** Residue interaction probability maps for A2 LCD (non-PTM: e, i) and methylated A2 LCD (f, j), simulated using Slab-PTM patched CALVADOS2 (e, f) and Mpipi. (i, j). Blue indicates stronger interactions; red indicates weaker interactions. **(k)** Sequence DDX4 N-terminal domain with asymmetric dimethylation sites indicated. **(l)** Experimental results comparing DDX4 and methylated DDX4^16^. **(m, q)** Protein density along the z-axis for DDX4 and its methylated form, simulated using Slab-PTM patched CALVADOS2 (m) and Mpipi (q). Blue represents the native IDP and red represents the PTM-modified IDP. **(n, r)** Phase transition curves of non-PTM and PTM systems under a temperature gradient, simulated using Slab-PTM patched CALVADOS2 (n) and Mpipi (r) Blue represents the native IDP and red represents the PTM-modified IDP. **(o, p, s, t)** Residue interaction probability maps for DDX4 (non-PTM: o, s) and methylated DDX4 (p, t), simulated using Slab-PTM patched CALVADOS2 (o, p) and Mpipi. (s, t). Blue indicates stronger interactions; red indicates weaker interactions.

Similar results were observed for the DDX4 N-terminal domain. As reported previously (Figure 5k,l), methylation in this region suppresses LLPS^16^. Our simulations show that while the non-PTM DDX4 forms a dense phase, the methylated version does not, in both CALVADOS2 and Mpipi (Figure 5m, q). Phase diagrams confirm that methylation lowers the critical temperature substantially (Figure 5n, r). Residue-level contact maps (Figure 5o, s and Figure 5p,t) again highlight reduced contact probability in methylated regions. Detailed interaction profiles are shown in Figures S19 and S20. Detailed protein-density plots are available for DDX4 N-terminal domain: the native protein in Figure S43 (patched CALVADOS2) and Figure S44 (patched Mpipi), and the acetylated protein in Figure S45 (patched CALVADOS2) and Figure S46 (patched Mpipi).

## Discussion

PTMs are central to the regulation of biomolecular condensates, yet their impact remains challenging to capture with current coarse-grained modeling frameworks. In this study, we addressed this limitation by developing Slab-PTM, a transferable parameter patch that integrates PMF-derived interaction strengths for five prevalent PTM types into widely used LLPS simulation models.

Our results demonstrate that Slab-PTM reliably reproduces experimentally observed changes in phase behavior across a diverse set of IDPs and PTMs, including both global and site-specific modifications. Notably, it captures both suppression and enhancement of phase separation, suggesting that the patch accurately reflects the biophysical consequences of PTMs on residue-level interactions. Notably, despite substantial differences in their baseline parameterizations, both CALVADOS2 and Mpipi produced highly consistent LLPS behaviors for PTM-containing IDPs when patched with Slab-PTM. This cross-model agreement highlights the strong transferability of our approach. As a result, Slab-PTM is well-positioned to support future developments in coarse-grained force fields by providing reliable PTM interaction parameters across diverse simulation platforms. To facilitate broader adoption, we have made an automated script available on https://github.com/decodermu/Slab-PTM for generating Slab-PTM patches compatible with new or evolving IDP models.

Beyond reproducing known LLPS outcomes, Slab-PTM enables deeper insight into the molecular grammar underlying these effects. By resolving residue-level interaction probabilities, we observed consistent patterns of PTM-induced disruption or promotion of multivalent contacts, often linked to electrostatic or π-interaction changes. These results highlight Slab-PTM not just as a practical modeling tool, but also to mechanistically decode the PTM-regulated IDP molecular grammar that controls the phase behavior.

However, substantial challenges remain. Currently, Slab-PTM supports only five types of PTMs, while a broader spectrum of modifications is known to modulate IDP phase behavior. For instance, glycosylation^47^, ubiquitination^48^, PARylation^49^, Neddylation^50^, and SUMOylation^51^ all play regulatory roles in LLPS across various biological systems. Many of these involve complex chemical moieties or structural alterations that extend beyond simple side-chain modifications, requiring new strategies for parameterization.

Additionally, some PTMs occur outside of canonical side chains. N-terminal acetylation, as reported in FUS, exemplifies modifications that require position-specific treatment and may not be well-captured by current residue-centered coarse-graining schemes^52^. Beyond proteins, RNA is a frequent and critical component of biomolecular condensates. PTMs on IDPs could also influence the interaction with RNA^45,52^.

In summary, Slab-PTM provides a practical and extensible parameter patch that facilitates the simulation of PTM-regulated IDP phase separation. By accurately capturing the residue-level interaction changes introduced by PTMs, it offers a valuable tool for probing the molecular grammar through which PTMs control condensate formation. Looking forward, Slab-PTM may also enable the rational design of synthetic, PTM-responsive regulatory elements within IDPs, offering a new avenue for engineering tunable biomolecular condensates in synthetic biology and cell programming applications.

## Supplemental Information

Figure S1-S56 and Table S1-S3.

## Acknowledgments

This work was supported in part by the National Natural Science Foundation of China (22237002 and T2321001).

