## Supplemental Information for "Slab-PTM: A Coarse-Grained Force-Field Parameter Patch for Modeling Post-Translational Modification Effects on Biomolecular Condensates"

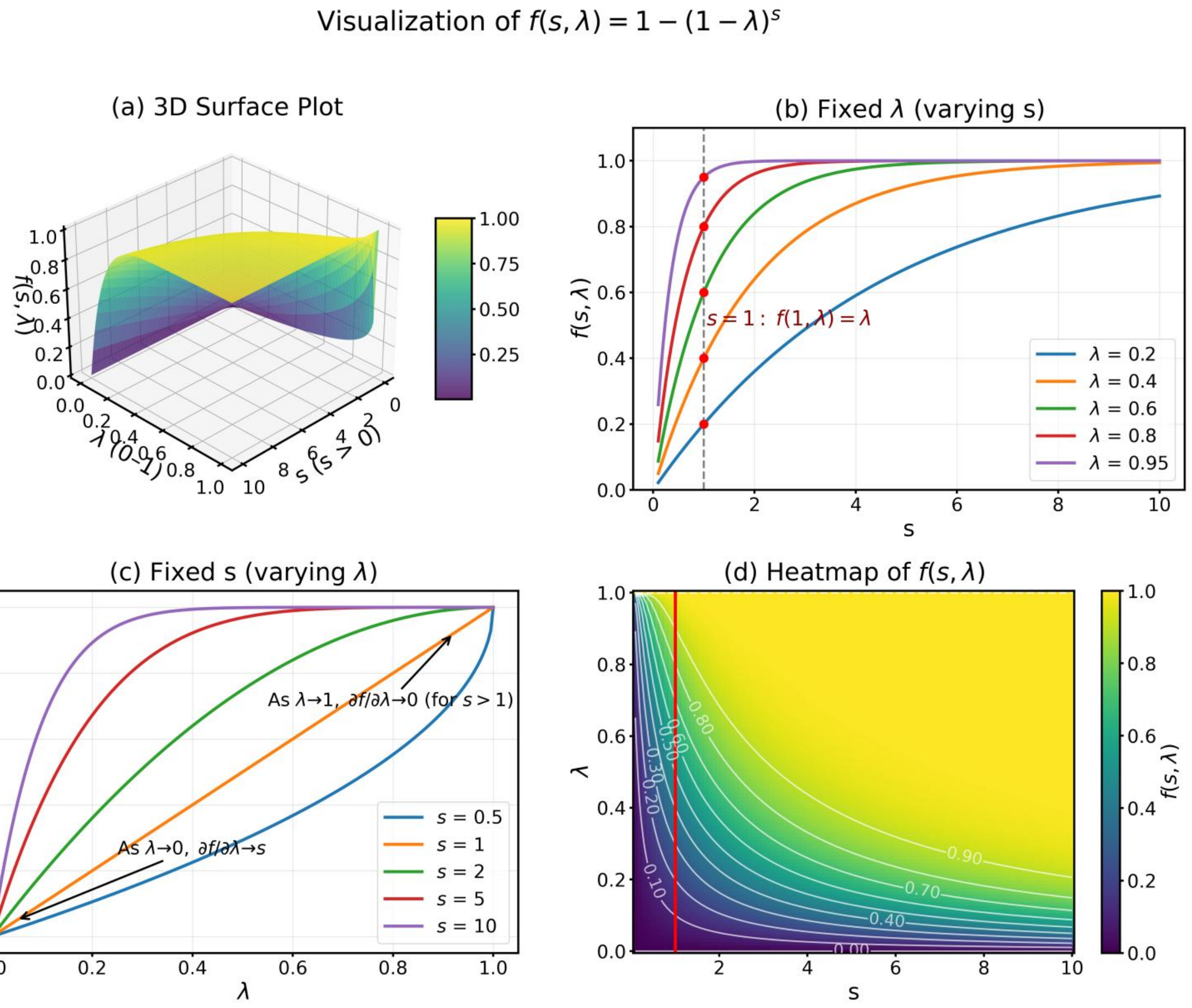


**Figure S1.** The detailed visualization of Eq. 15. (a) The 3D surface of all resulting λ under various inputs of s and λ. (b) The resulting λ under various inputs of s and fixed λ. When s=1, the result λ always equal to the input one. (c) The resulting λ under various inputs of λ and fixed s. When λ is close to 1, the partial derivative of the function with respect to λ approaches 0; when λ is close to 0, the partial derivative of the function with respect to λ approaches s. (d) The heat-map of formation of (a).

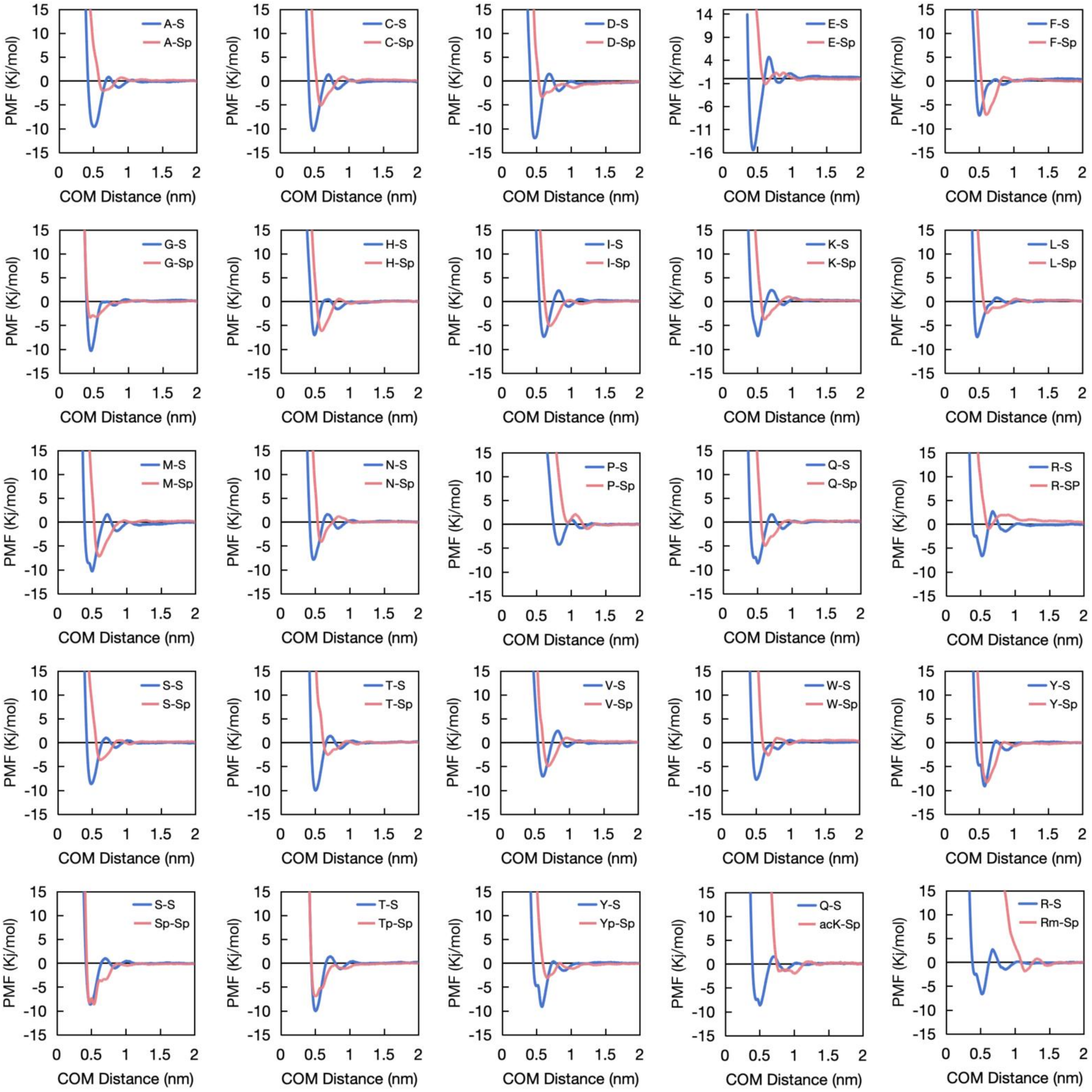


**Figure S2.** The PMF curves of phosphoserine (Sp) associated amino acid dipeptide pairs (pink curves) and their reference dipeptide pairs (blue curves).

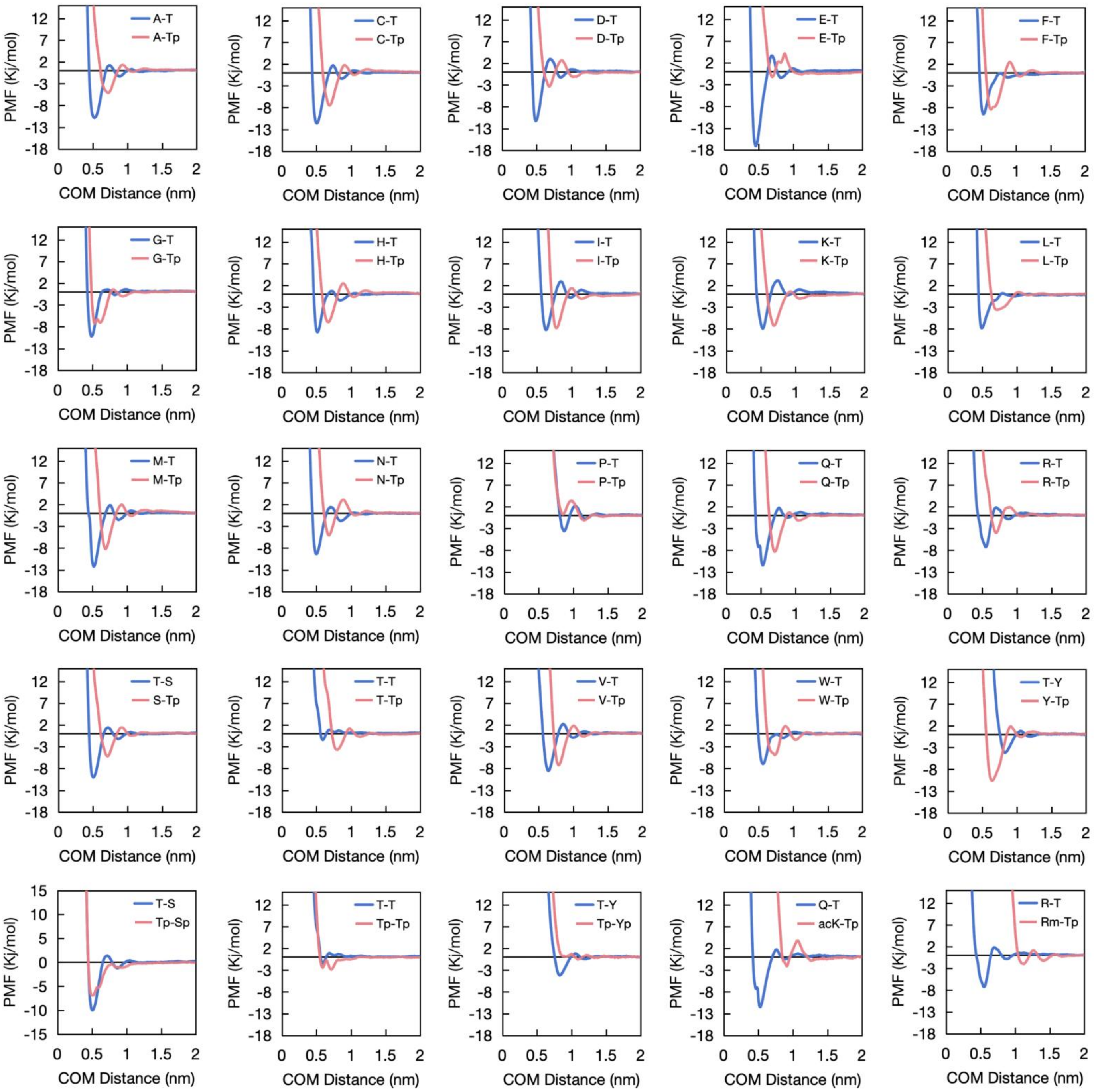

**Figure S3.** The PMF curves of phosphothreonine (Tp) associated amino acid dipeptide pairs (pink curves) and their reference dipeptide pairs (blue curves).

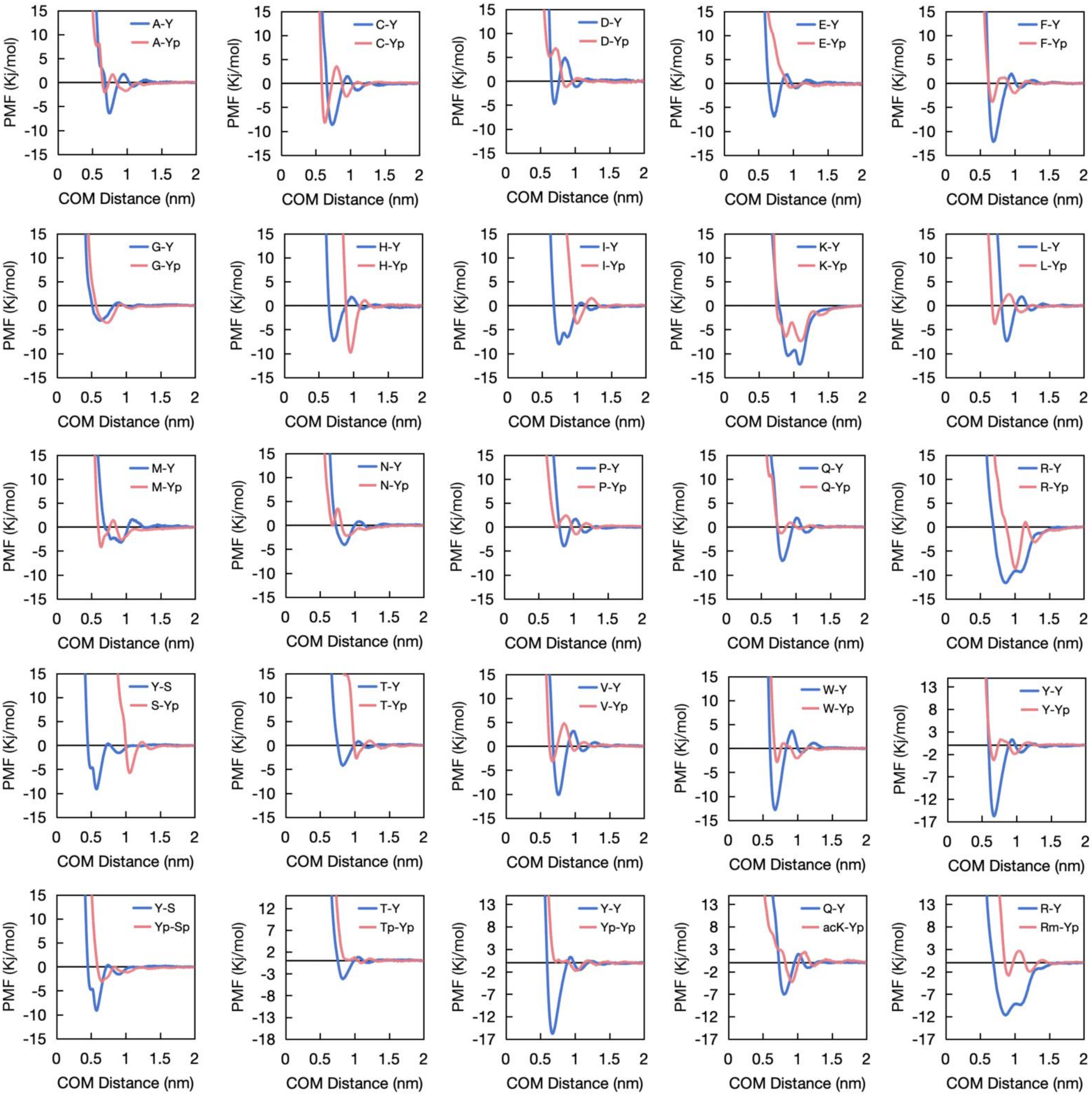


**Figure S4.** The PMF curves of phosphotyrosine (Yp) associated amino acid dipeptide pairs (pink curves) and their reference dipeptide pairs (blue curves).

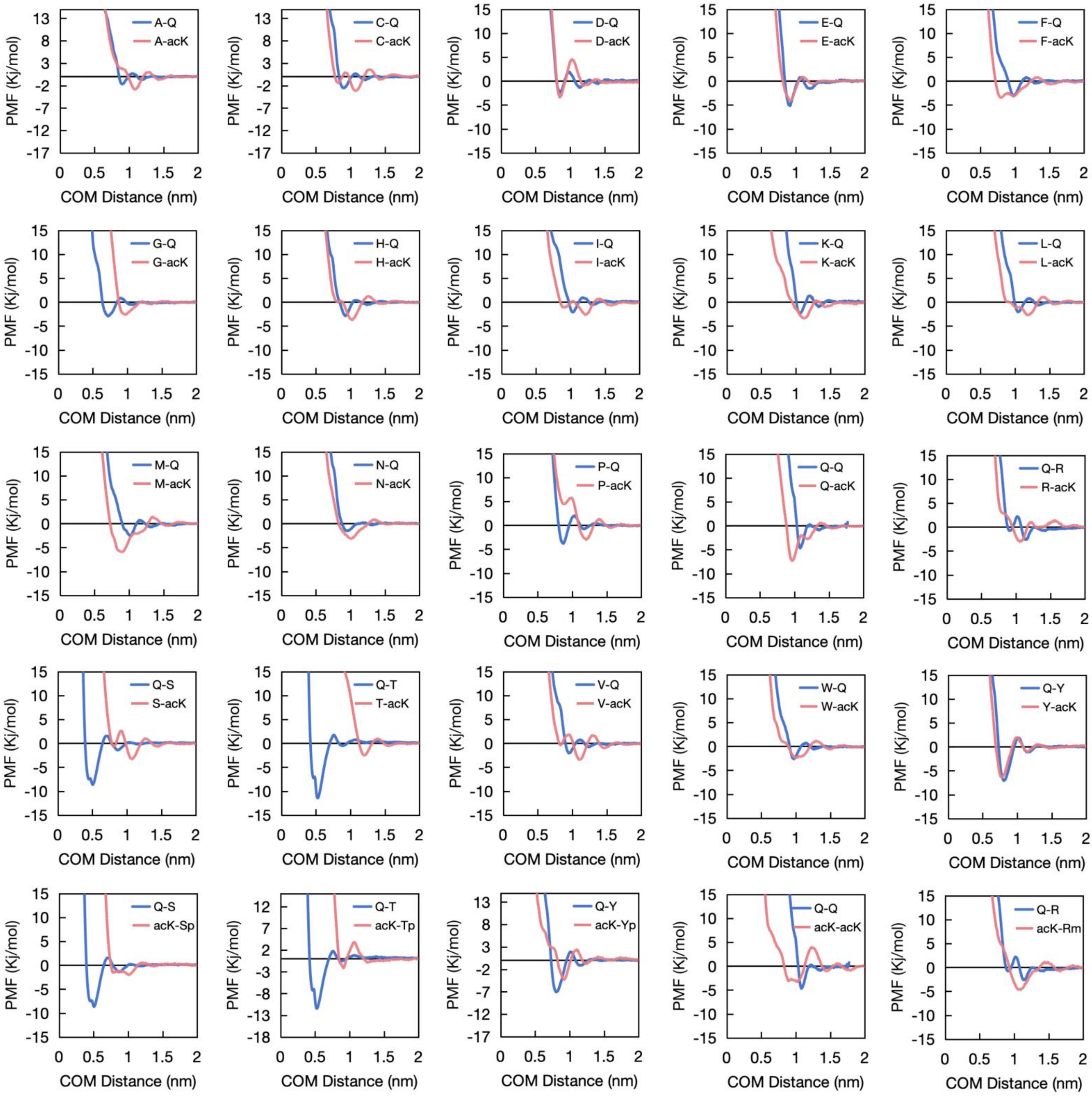


**Figure S5.** The PMF curves of acetyllysine (acK) associated amino acid dipeptide pairs (pink curves) and their reference dipeptide pairs (blue curves).

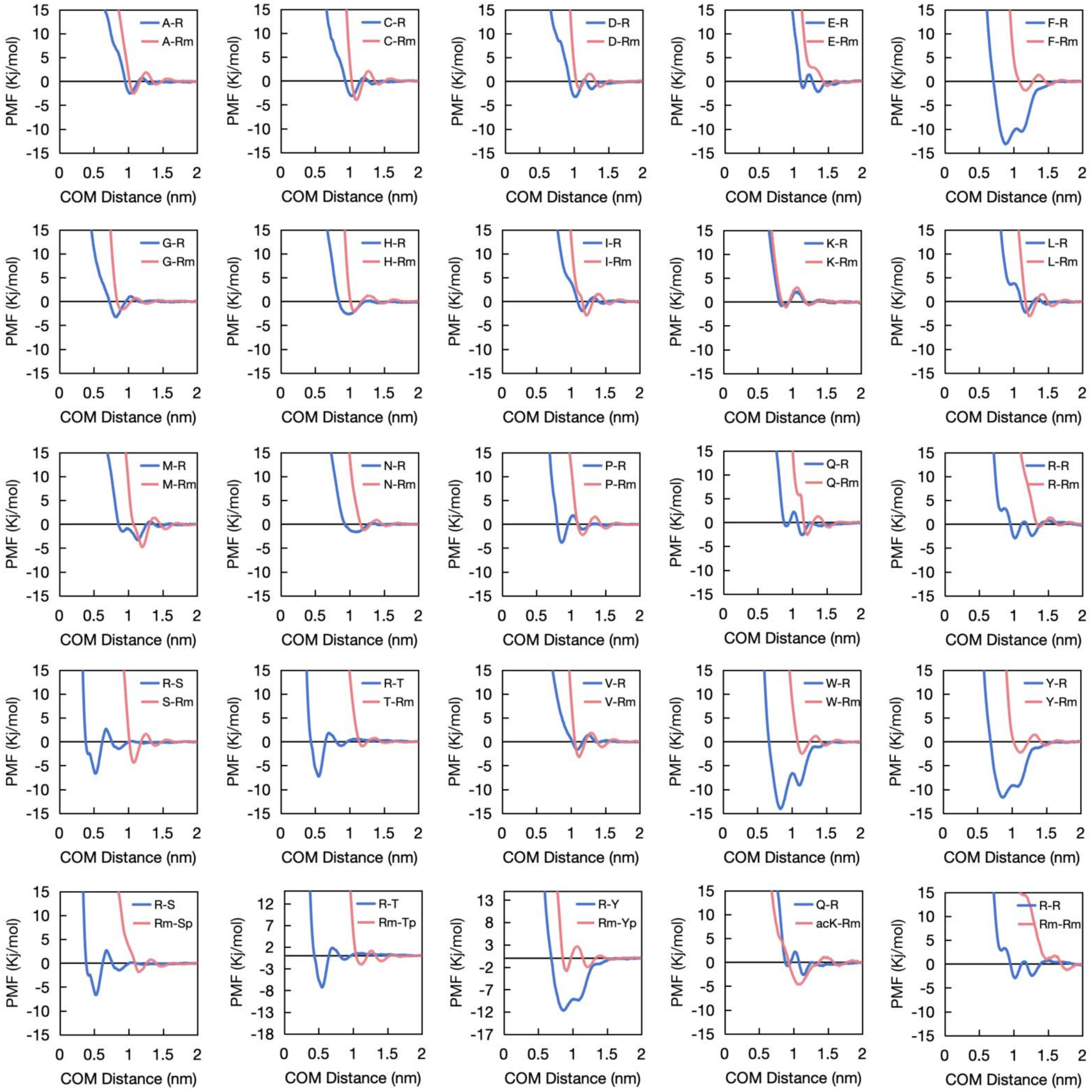


**Figure S6.** The PMF curves of asymmetric dimethylarginine (Rm) associated amino acid dipeptide pairs (pink curves) and their reference dipeptide pairs (blue curves).

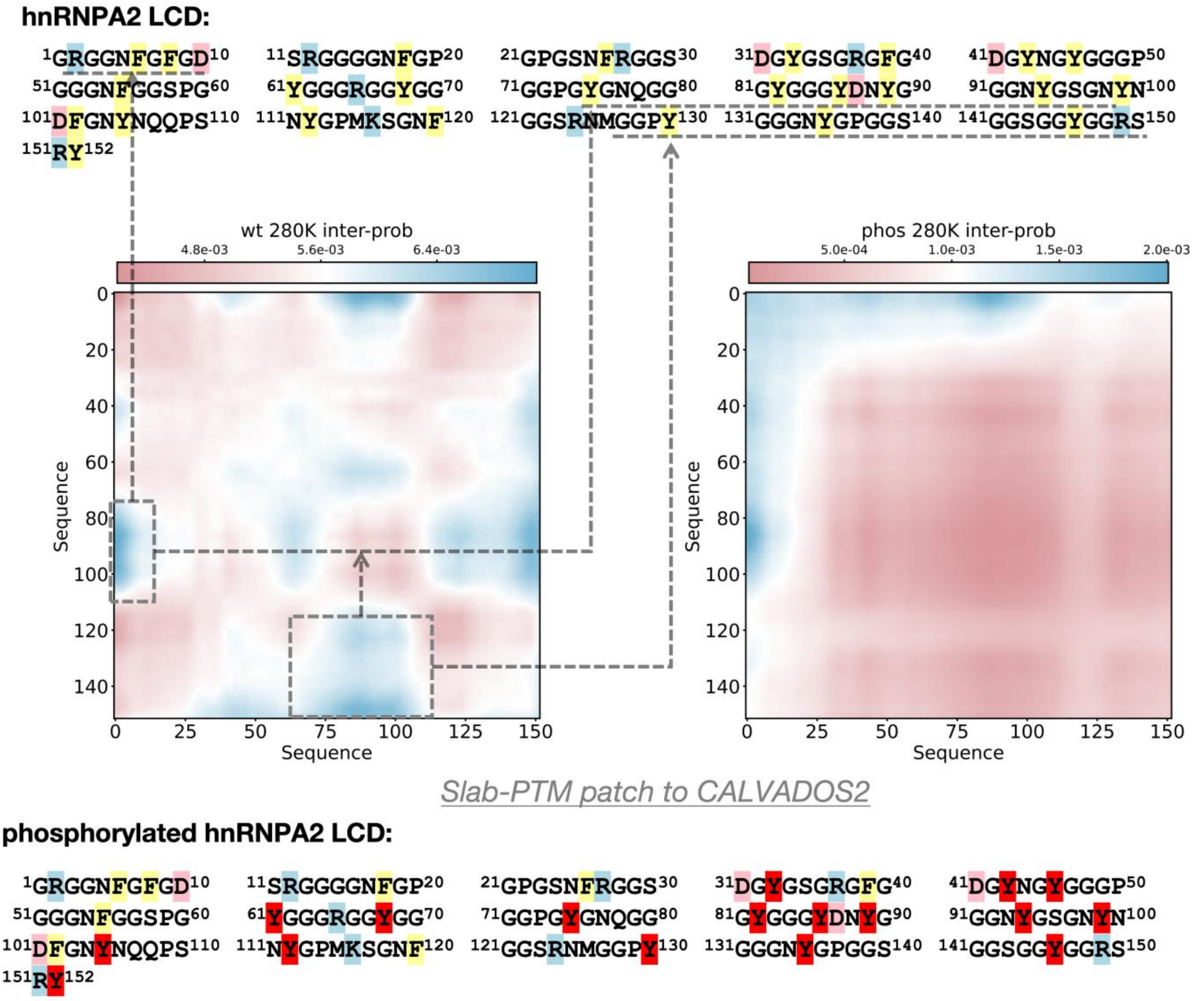


**Figure S7.** The sequence of hnRNPA2 LCD and the residue-level averaged interaction probability map derived from the LLPS MD trajectory, simulated using the Slab-PTM–patched CALVADOS2 model. Positively charged residues (R, K) are shown with a light blue background; negatively charged residues (D, E) with a pink background; aromatic residues (F, Y, W) with a yellow background; and PTM-modified sites with a red background.

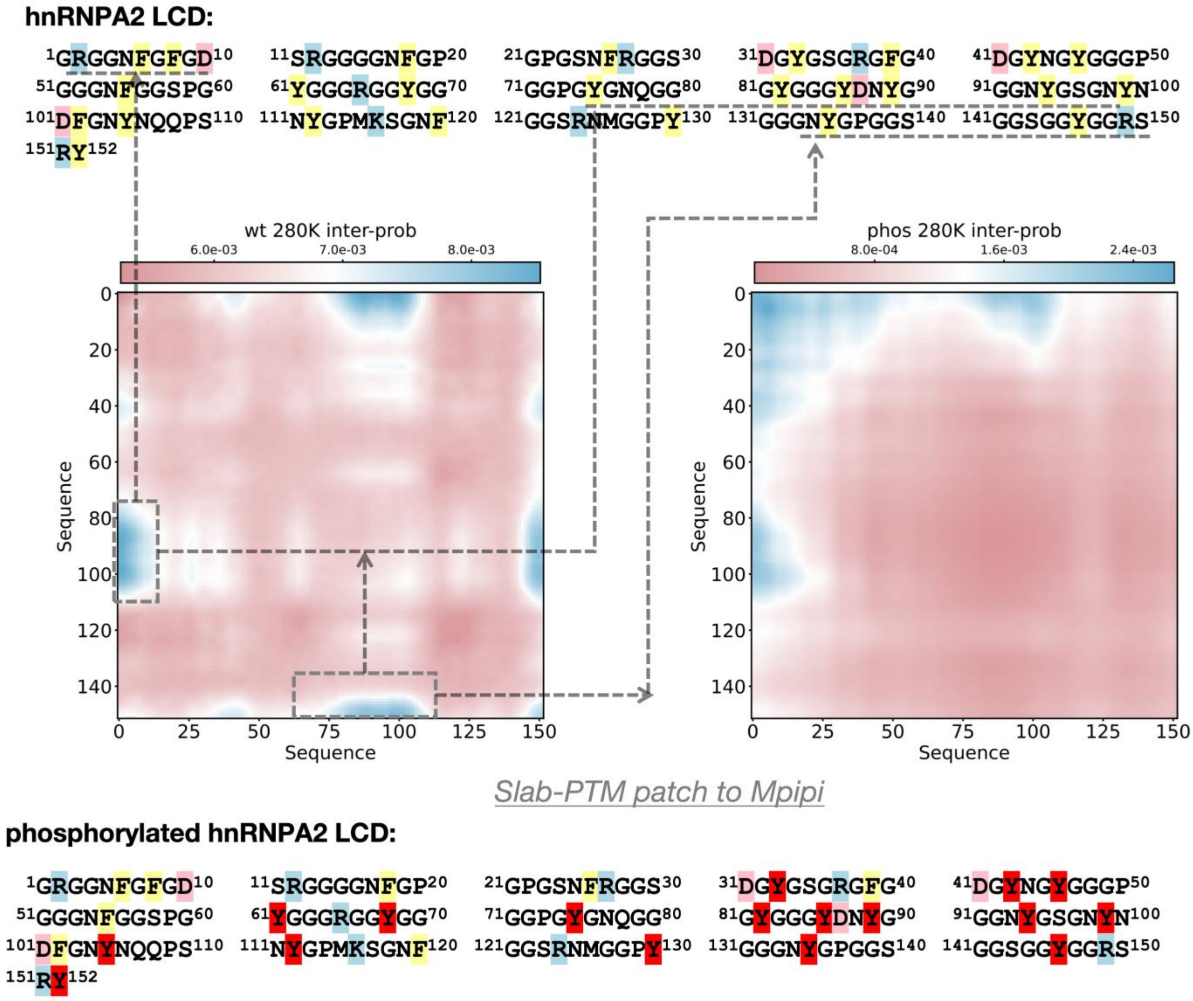


**Figure S8.** The sequence of hnRNPA2 LCD and the residue-level averaged interaction probability map derived from the LLPS MD trajectory, simulated using the Slab-PTM–patched Mpipi model. Positively charged residues (R, K) are shown with a light blue background; negatively charged residues (D, E) with a pink background; aromatic residues (F, Y, W) with a yellow background; and PTM-modified sites with a red background.

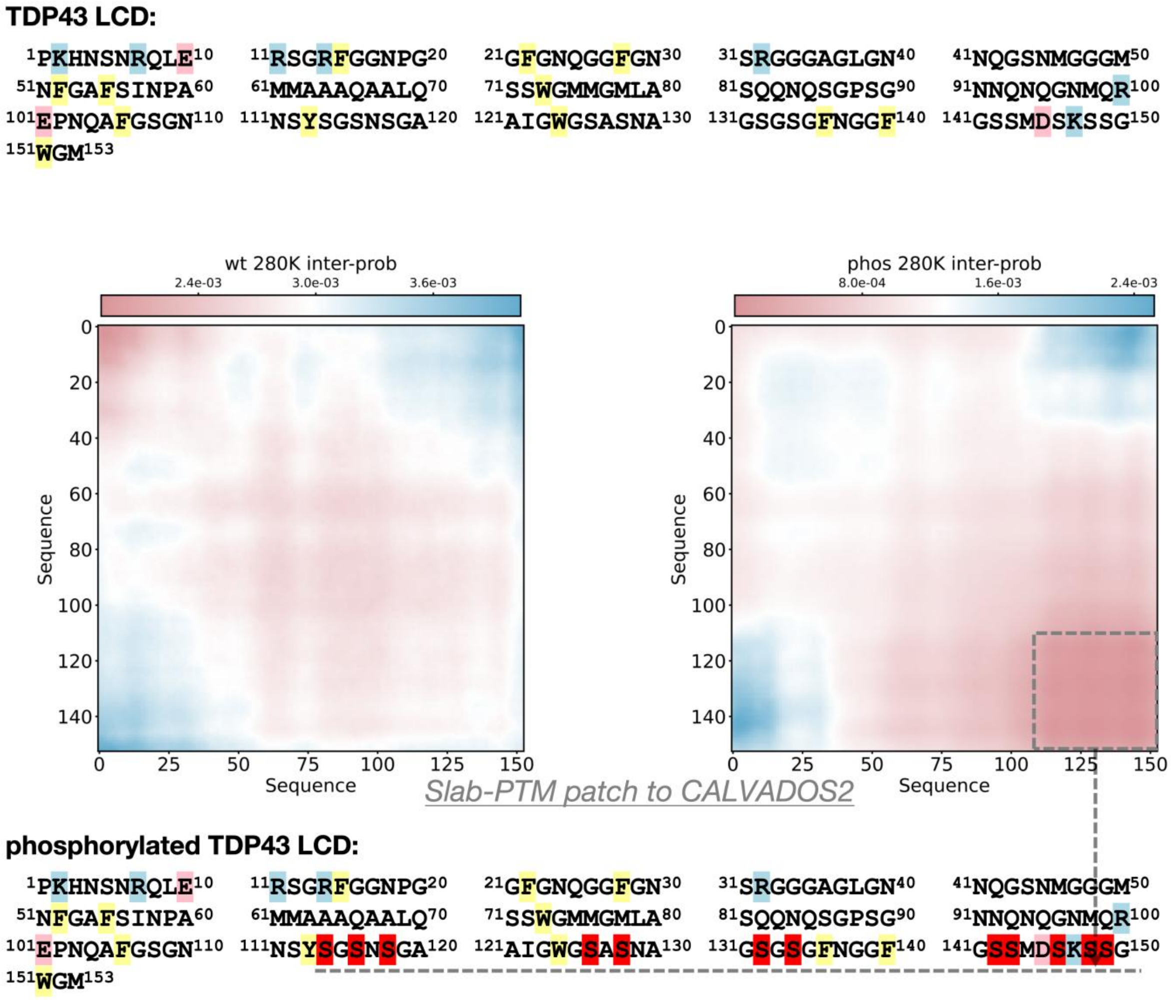


**Figure S9.** The sequence of TDP43 LCD and the residue-level averaged interaction probability map derived from the LLPS MD trajectory, simulated using the Slab-PTM–patched CALVADOS2 model. Positively charged residues (R, K) are shown with a light blue background; negatively charged residues (D, E) with a pink background; aromatic residues (F, Y, W) with a yellow background; and PTM-modified sites with a red background.

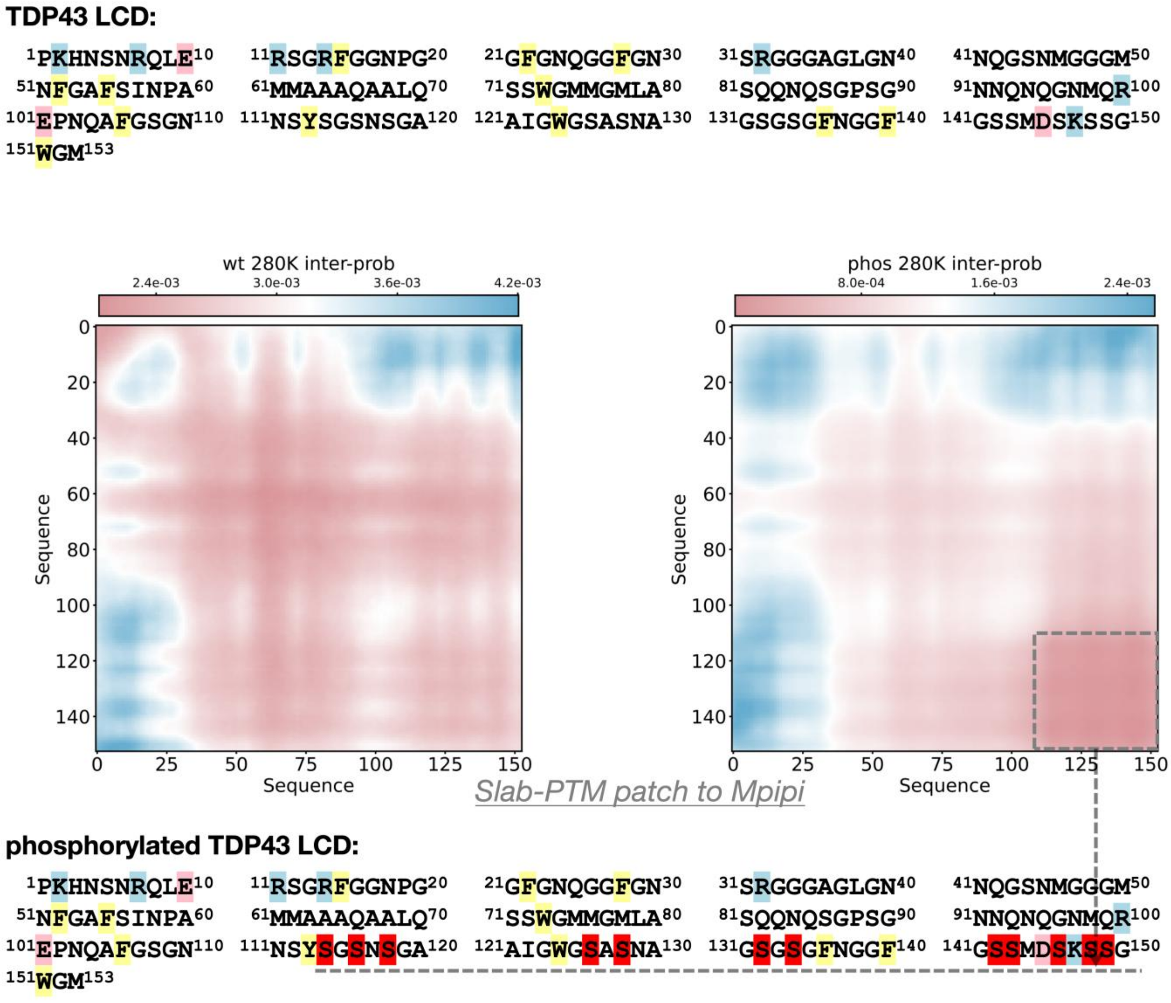


**Figure S10.** The sequence of TDP43 LCD and the residue-level averaged interaction probability map derived from the LLPS MD trajectory, simulated using the Slab-PTM–patched Mpipi model. Positively charged residues (R, K) are shown with a light blue background; negatively charged residues (D, E) with a pink background; aromatic residues (F, Y, W) with a yellow background; and PTM-modified sites with a red background.

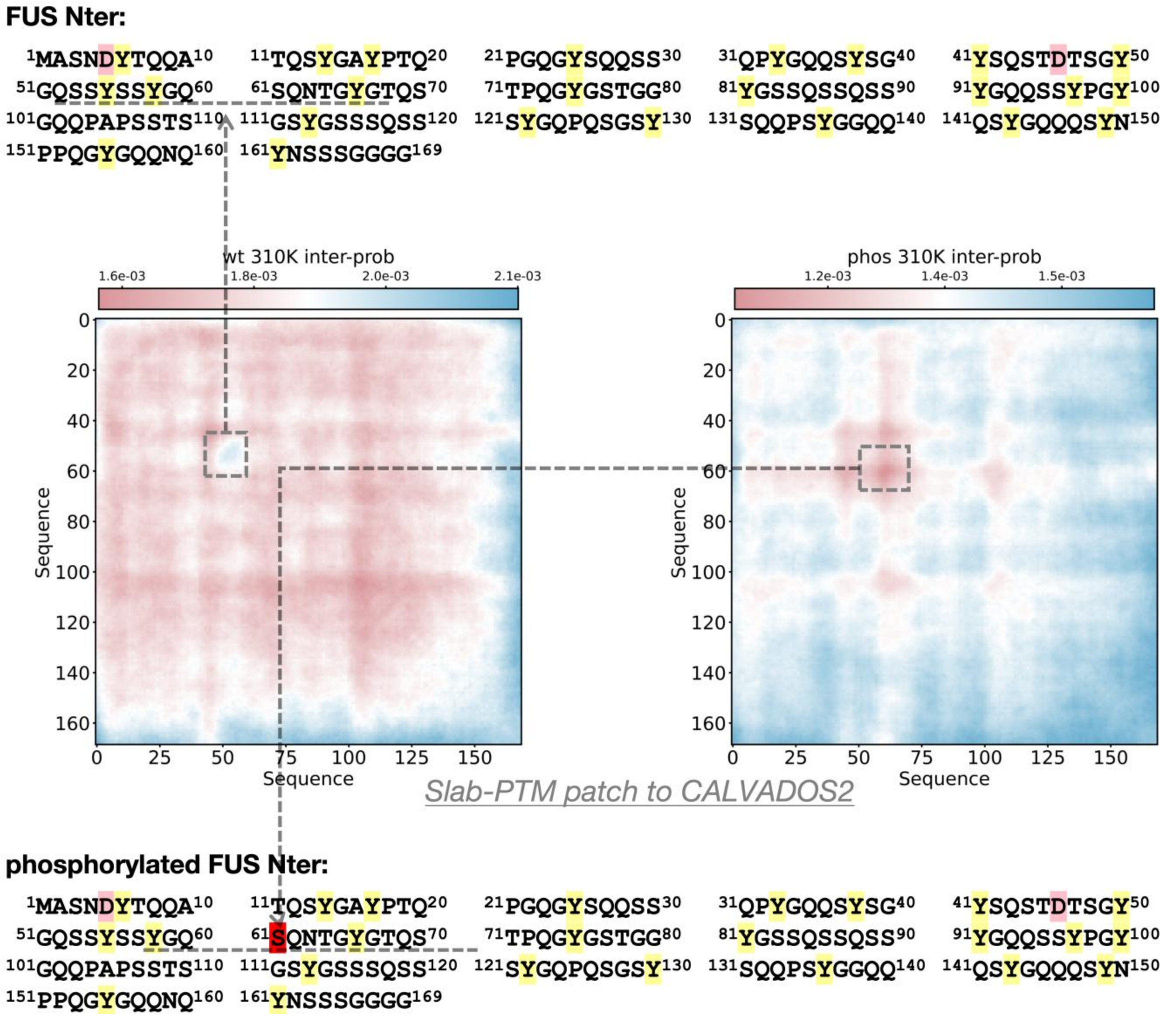


**Figure S11.** The sequence of FUS N-terminal domain and the residue-level averaged interaction probability map derived from the LLPS MD trajectory, simulated using the Slab-PTM–patched CALVADOS2 model. Positively charged residues (R, K) are shown with a light blue background; negatively charged residues (D, E) with a pink background; aromatic residues (F, Y, W) with a yellow background; and PTM-modified sites with a red background.

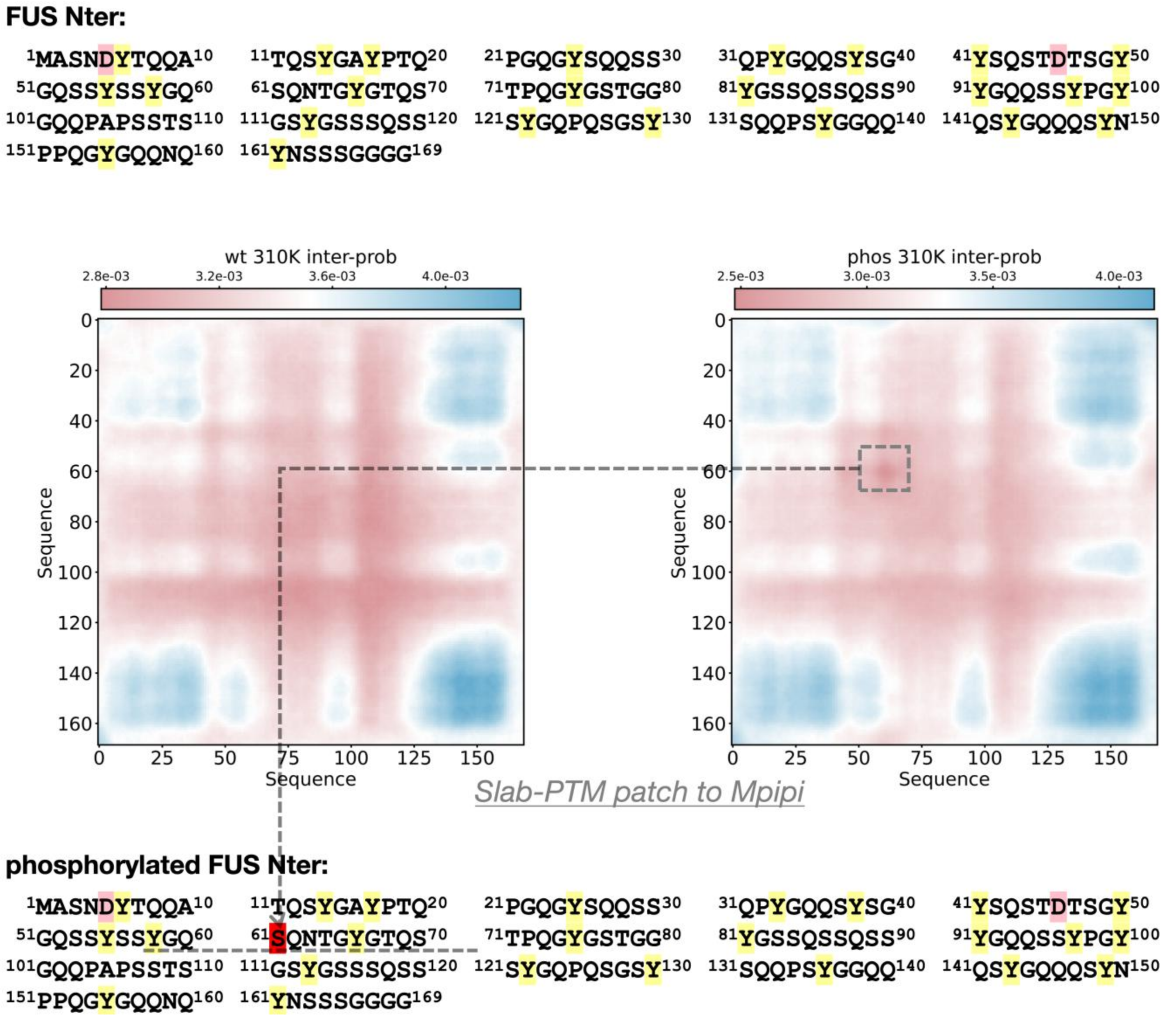


**Figure S12.** The sequence of FUS N-terminal domain and the residue-level averaged interaction probability map derived from the LLPS MD trajectory, simulated using the Slab-PTM–patched Mpipi model. Positively charged residues (R, K) are shown with a light blue background; negatively charged residues (D, E) with a pink background; aromatic residues (F, Y, W) with a yellow background; and PTM-modified sites with a red background.

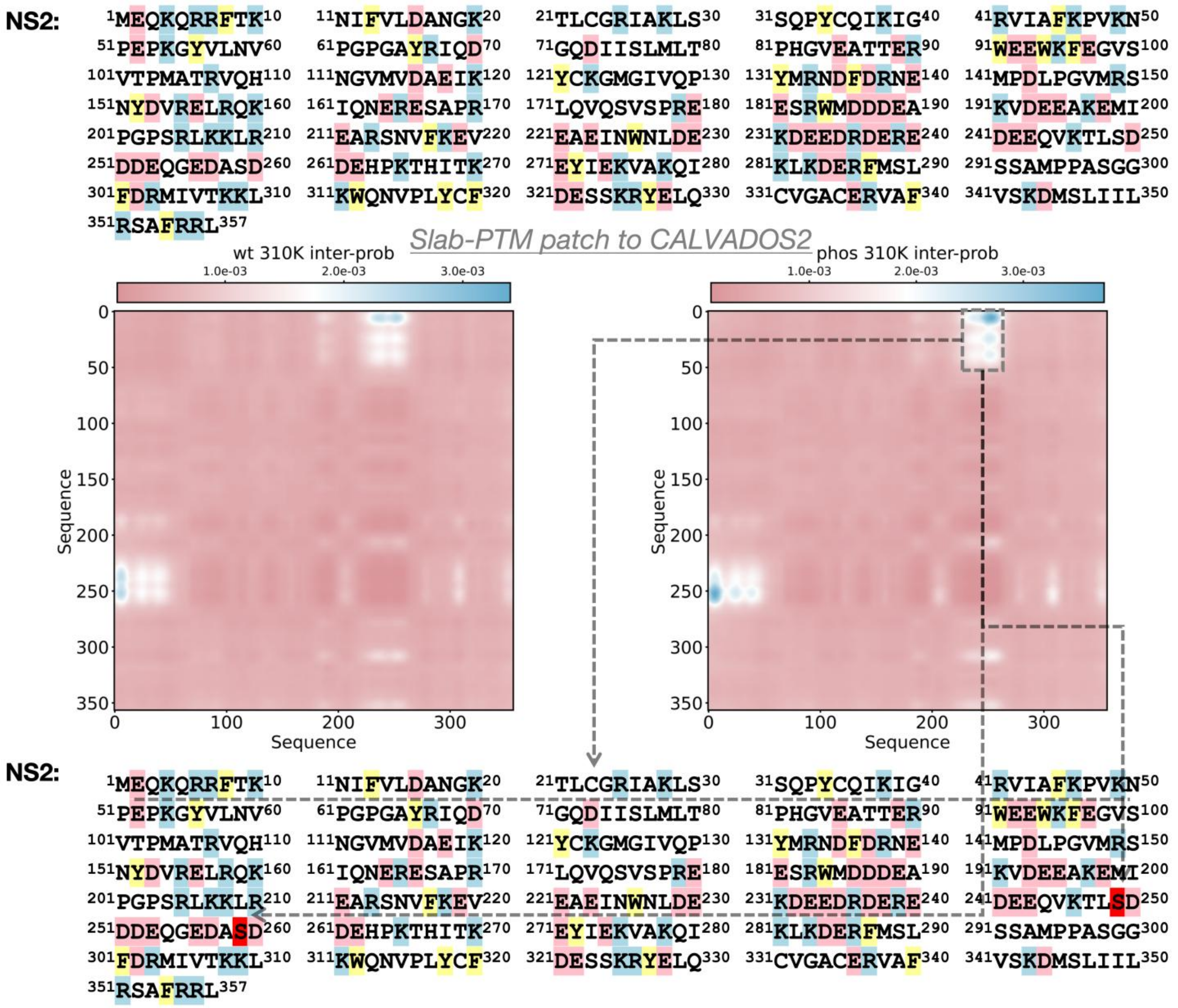


**Figure S13.** The sequence of NS2 and the residue-level averaged interaction probability map derived from the LLPS MD trajectory, simulated using the Slab-PTM–patched CALVADOS2 model. Positively charged residues (R, K) are shown with a light blue background; negatively charged residues (D, E) with a pink background; aromatic residues (F, Y, W) with a yellow background; and PTM-modified sites with a red background.

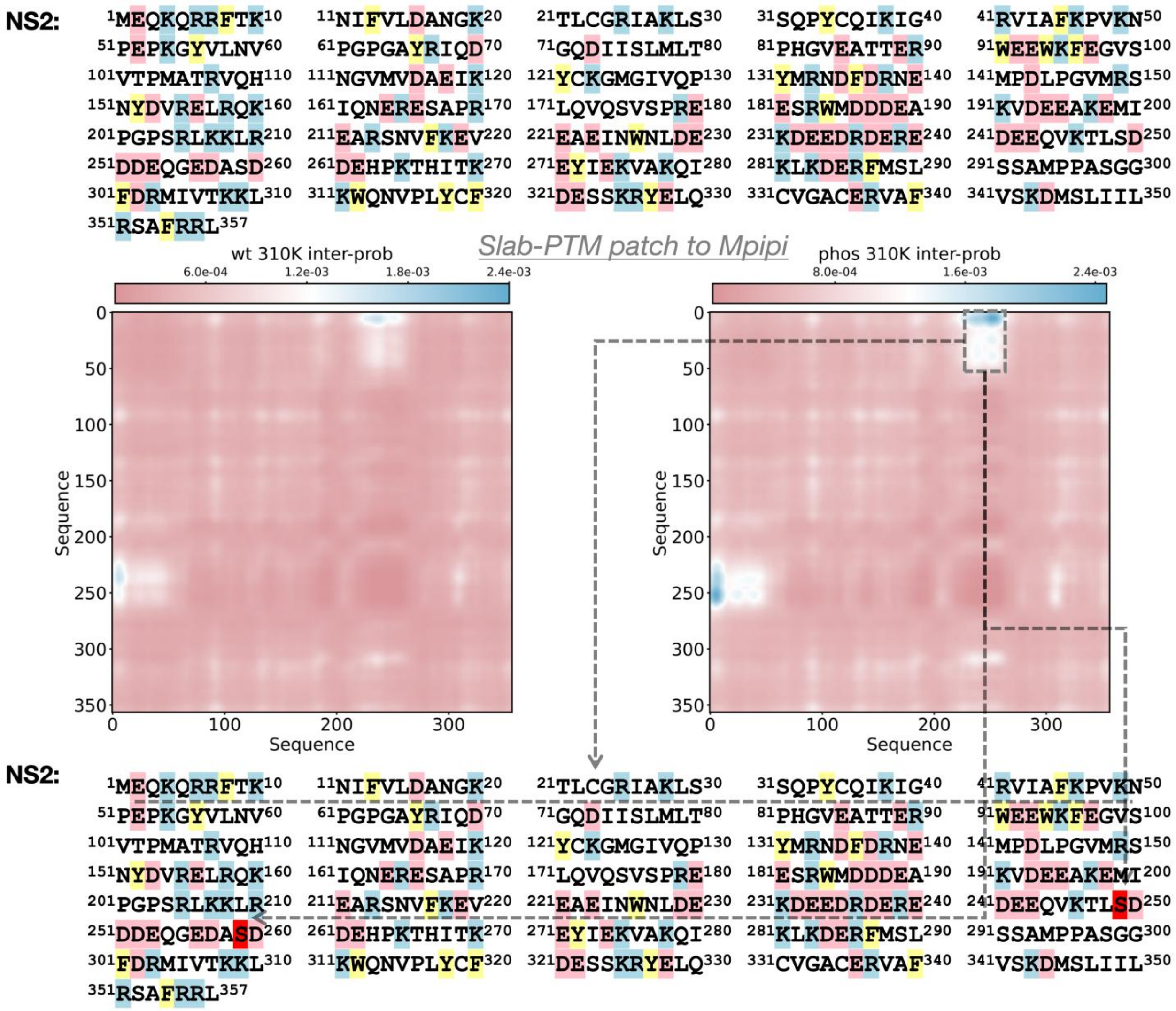


**Figure S14.** The sequence of NS2 and the residue-level averaged interaction probability map derived from the LLPS MD trajectory, simulated using the Slab-PTM–patched Mpipi model. Positively charged residues (R, K) are shown with a light blue background; negatively charged residues (D, E) with a pink background; aromatic residues (F, Y, W) with a yellow background; and PTM-modified sites with a red background.

**DDX3X IDR1:**

$^{1}$MSHVAVENAL$^{10}$ $^{11}$GLDQQFAGLD$^{20}$ $^{21}$LNSSDNQSGG$^{30}$ $^{31}$STASKGRYIP$^{40}$ $^{41}$PHLRNREATK$^{50}$
$^{51}$GFYDKDSSGW$^{60}$ $^{61}$SSSKDKDAYS$^{70}$ $^{71}$SFGSRSDSRG$^{80}$ $^{81}$KSSFFSDRGS$^{90}$ $^{91}$GSRGRFDDRG$^{100}$
$^{101}$RSDYDGIGSR$^{110}$ $^{111}$GDRSGFGKFE$^{120}$ $^{121}$RGGNSRWCDK$^{130}$ $^{131}$SDEDDWSKPL$^{140}$ $^{141}$PPSERLEQEL$^{150}$
$^{151}$FSGGNTGINF$^{160}$ $^{161}$EKYDDIPV$^{168}$

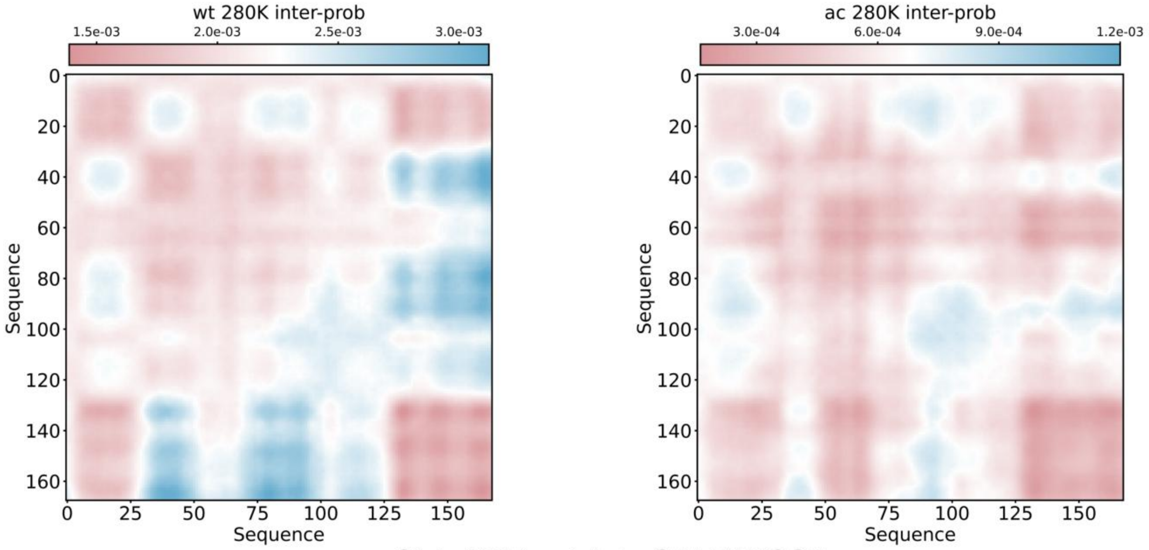


Slab-PTM patch to CALVADOS2

**acetylated DDX3X IDR1:**

$^{1}$MSHVAVENAL$^{10}$ $^{11}$GLDQQFAGLD$^{20}$ $^{21}$LNSSDNQSGG$^{30}$ $^{31}$STASKGRYIP$^{40}$ $^{41}$PHLRNREATK$^{50}$
$^{51}$GFYDKDSSGW$^{60}$ $^{61}$SSSKDKDAYS$^{70}$ $^{71}$SFGSRSDSRG$^{80}$ $^{81}$KSSFFSDRGS$^{90}$ $^{91}$GSRGRFDDRG$^{100}$
$^{101}$RSDYDGIGSR$^{110}$ $^{111}$GDRSGFGKFE$^{120}$ $^{121}$RGGNSRWCDK$^{130}$ $^{131}$SDEDDWSKPL$^{140}$ $^{141}$PPSERLEQEL$^{150}$
$^{151}$FSGGNTGINF$^{160}$ $^{161}$EKYDDIPV$^{168}$

**Figure S15.** The sequence of DDX3X IDR1 domain and the residue-level averaged interaction probability map derived from the LLPS MD trajectory, simulated using the Slab-PTM–patched CALVADOS2 model. Positively charged residues (R, K) are shown with a light blue background; negatively charged residues (D, E) with a pink background; aromatic residues (F, Y, W) with a yellow background; and PTM-modified sites with a red background.

**DDX3X IDR1:**

1MSHVAVENAL10 11GLDQQFAGLD20 21LNSSDNQSGG30 31STASKGRYIP40 41PHLRNREATK50
51GFYDKDSSGW60 61SSSKDKDAYS70 71SFGSRSDSRG80 81KSSFFSDRGS90 91GSRGRFDDRG100
101RSDYDGIGSR110 111GDRSGFGKFE120 121RGGNSRWCDK130 131SDEDDWSKPL140 141PPSERLEQEL150
151FSGGNTGINF160 161EKYDDIPV168

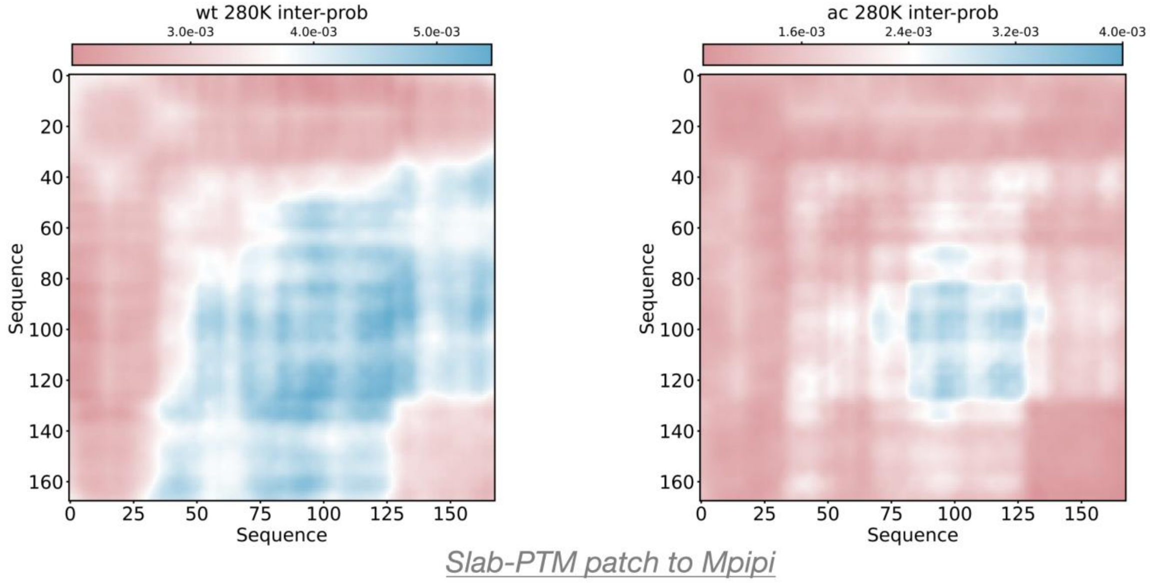


**acetylated DDX3X IDR1:**

1MSHVAVENAL10 11GLDQQFAGLD20 21LNSSDNQSGG30 31STASKGRYIP40 41PHLRNREATK50
51GFYDKDSSGW60 61SSSKDKDAYS70 71SFGSRSDSRG80 81KSSFFSDRGS90 91GSRGRFDDRG100
101RSDYDGIGSR110 111GDRSGFGKFE120 121RGGNSRWCDK130 131SDEDDWSKPL140 141PPSERLEQEL150
151FSGGNTGINF160 161EKYDDIPV168

**Figure S16.** The sequence of DDX3X IDR1 domain and the residue-level averaged interaction probability map derived from the LLPS MD trajectory, simulated using the Slab-PTM–patched Mpipi model. Positively charged residues (R, K) are shown with a light blue background; negatively charged residues (D, E) with a pink background; aromatic residues (F, Y, W) with a yellow background; and PTM-modified sites with a red background.

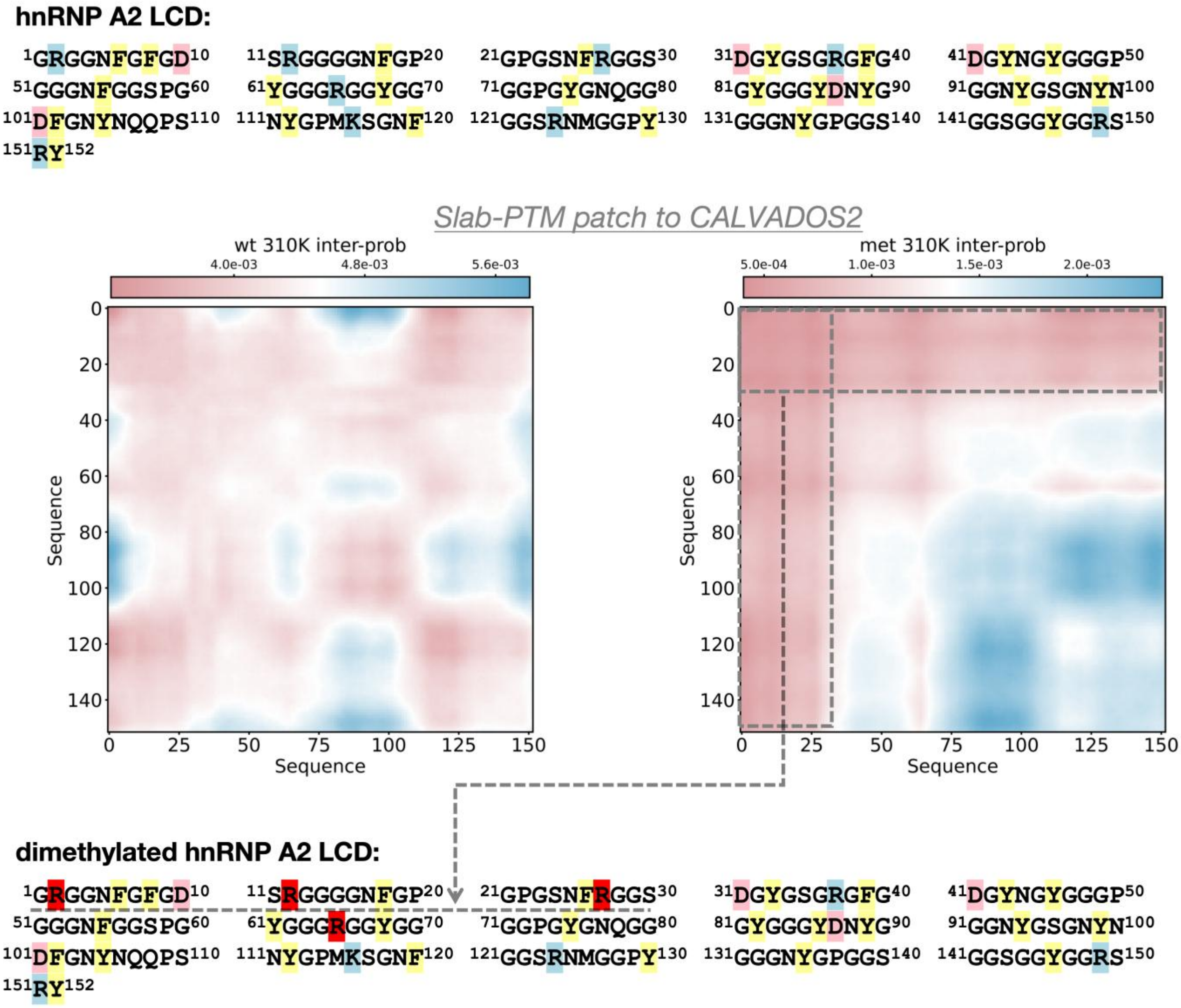


**Figure S17.** The sequence of hnRNPA2 LCD domain and the residue-level averaged interaction probability map derived from the LLPS MD trajectory, simulated using the Slab-PTM–patched CALVADOS2 model. Positively charged residues (R, K) are shown with a light blue background; negatively charged residues (D, E) with a pink background; aromatic residues (F, Y, W) with a yellow background; and PTM-modified sites with a red background.

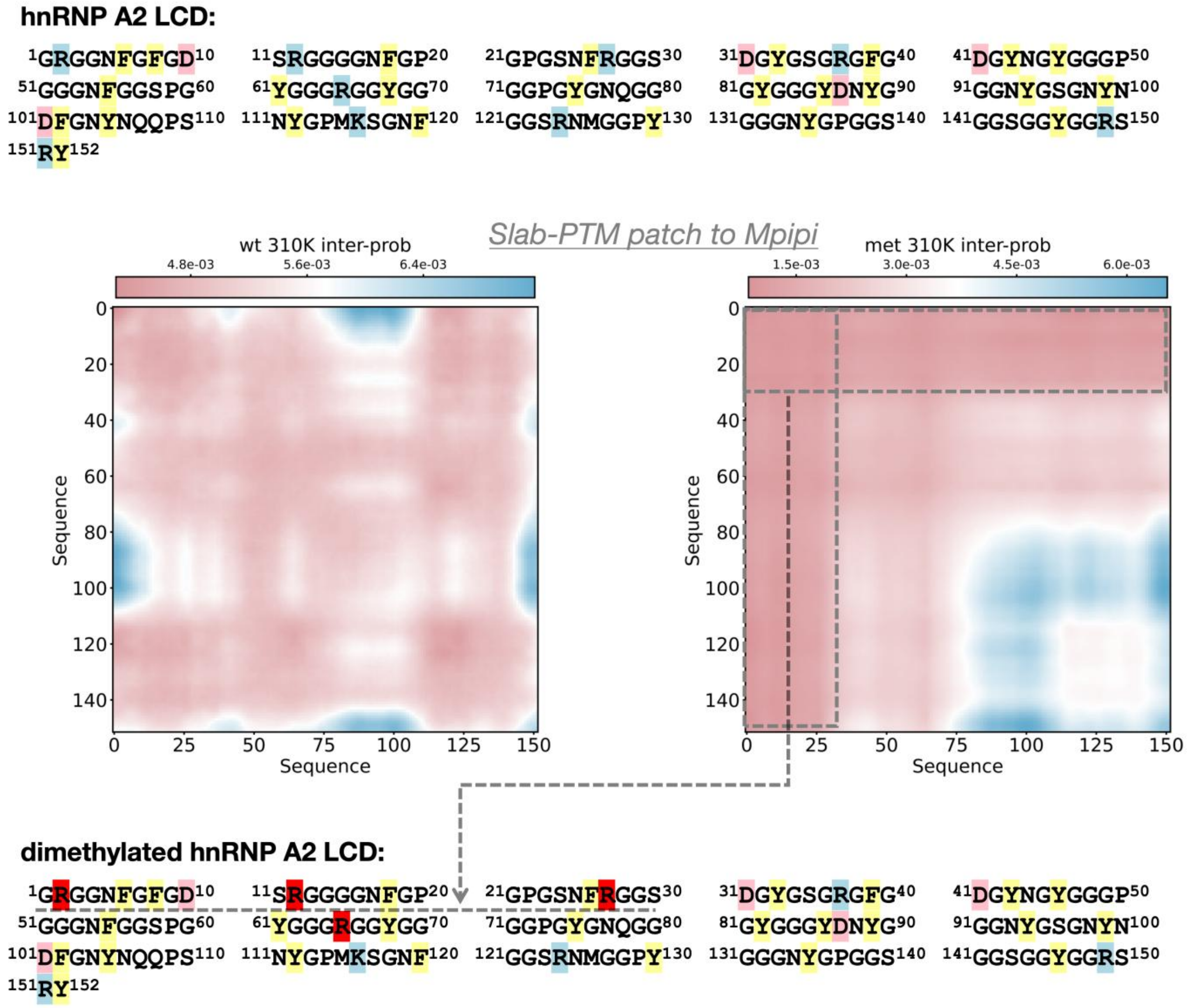


**Figure S18.** The sequence of hnRNPA2 LCD domain and the residue-level averaged interaction probability map derived from the LLPS MD trajectory, simulated using the Slab-PTM–patched Mpipi model. Positively charged residues (R, K) are shown with a light blue background; negatively charged residues (D, E) with a pink background; aromatic residues (F, Y, W) with a yellow background; and PTM-modified sites with a red background.

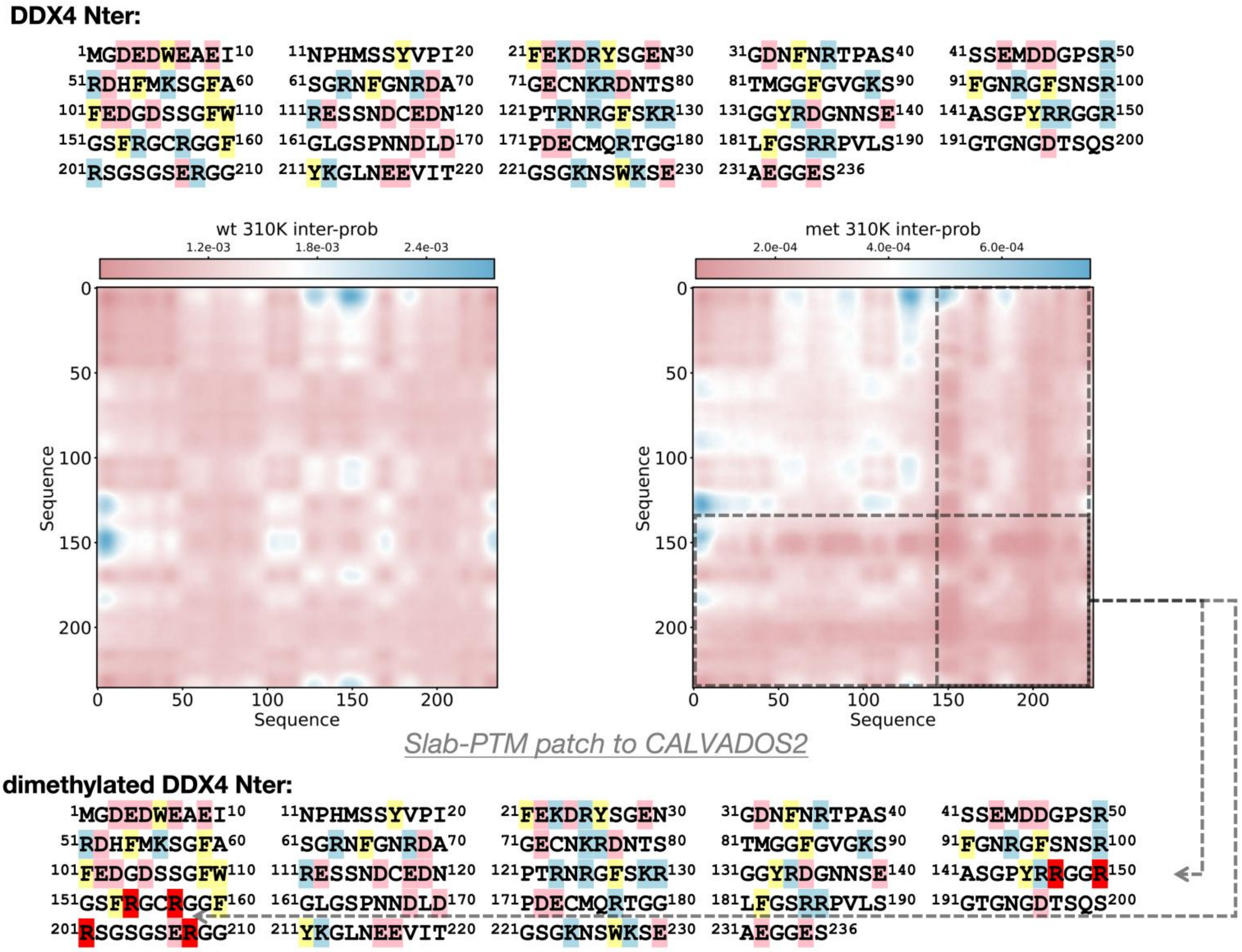


**Figure S19.** The sequence of DDX4 N-terminal domain and the residue-level averaged interaction probability map derived from the LLPS MD trajectory, simulated using the Slab-PTM–patched CALVADOS2 model. Positively charged residues (R, K) are shown with a light blue background; negatively charged residues (D, E) with a pink background; aromatic residues (F, Y, W) with a yellow background; and PTM-modified sites with a red background.

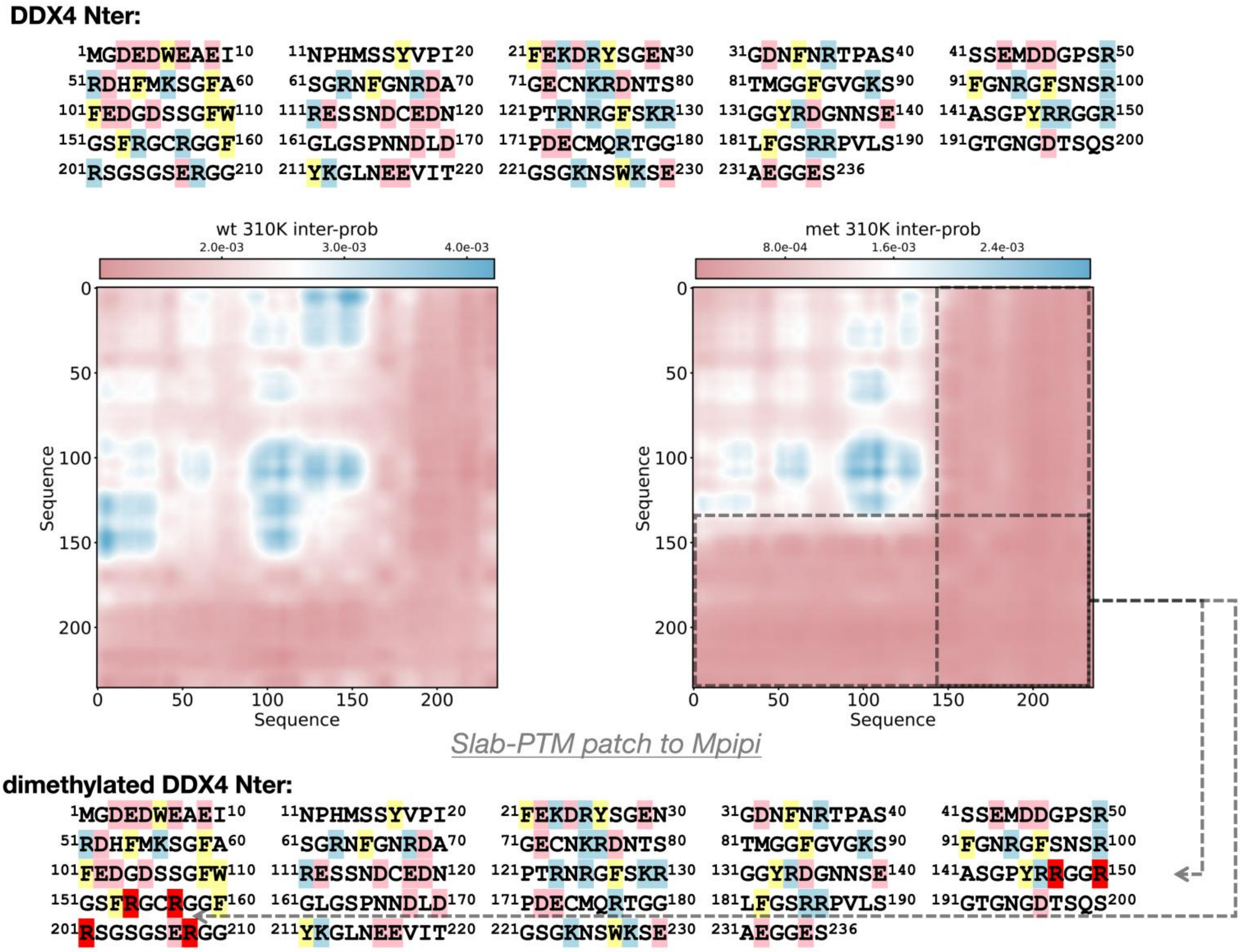


**Figure S20.** The sequence of DDX4 N-terminal domain and the residue-level averaged interaction probability map derived from the LLPS MD trajectory, simulated using the Slab-PTM–patched Mpipi model. Positively charged residues (R, K) are shown with a light blue background; negatively charged residues (D, E) with a pink background; aromatic residues (F, Y, W) with a yellow background; and PTM-modified sites with a red background.

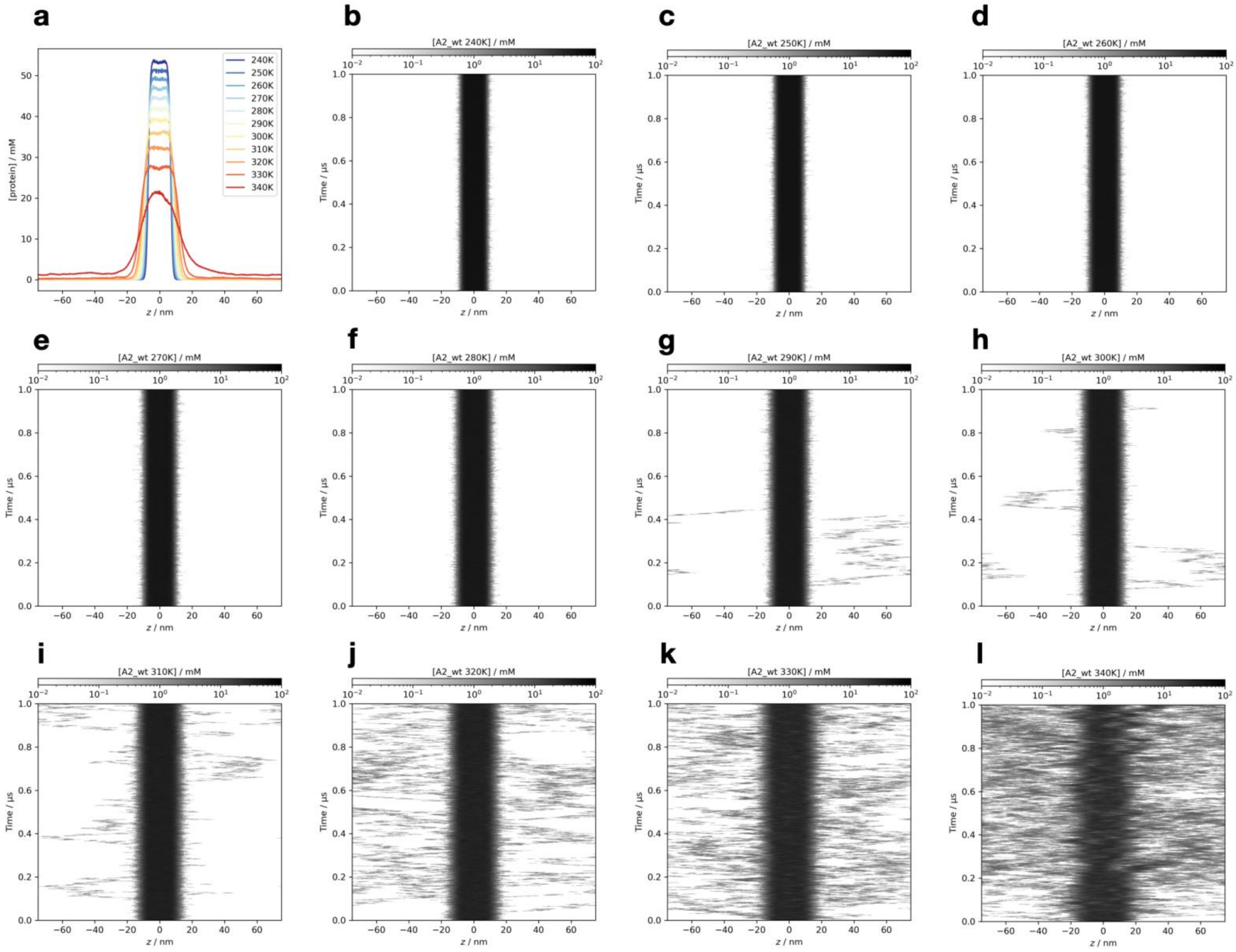

**Figure S21.** The detailed simulation results of the non-PTM hnRNP A2 LCD at all temperatures, simulated using the Slab-PTM patched CALVADOS2 model. (a) The average protein density along the z-axis. (b-l) The time-resolved average protein density along the z-axis from simulation trajectories at temperatures ranging from 240 K to 340 K.

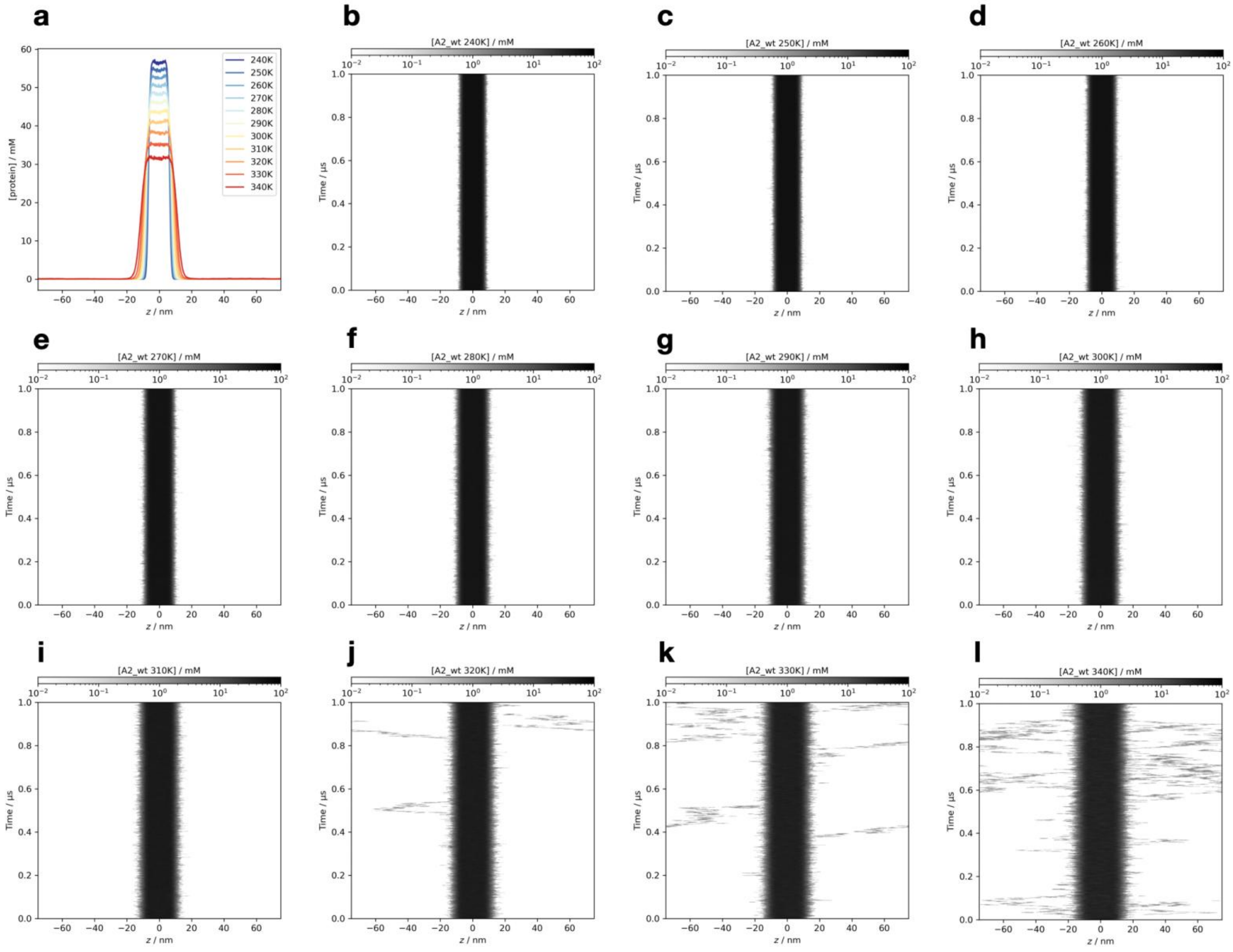


**Figure S22.** The detailed simulation results of the non-PTM hnRNP A2 LCD at all temperatures, simulated using the Slab-PTM patched Mpipi model. (a) The average protein density along the z-axis. (b-l) The time-resolved average protein density along the z-axis from simulation trajectories at temperatures ranging from 240 K to 340 K.

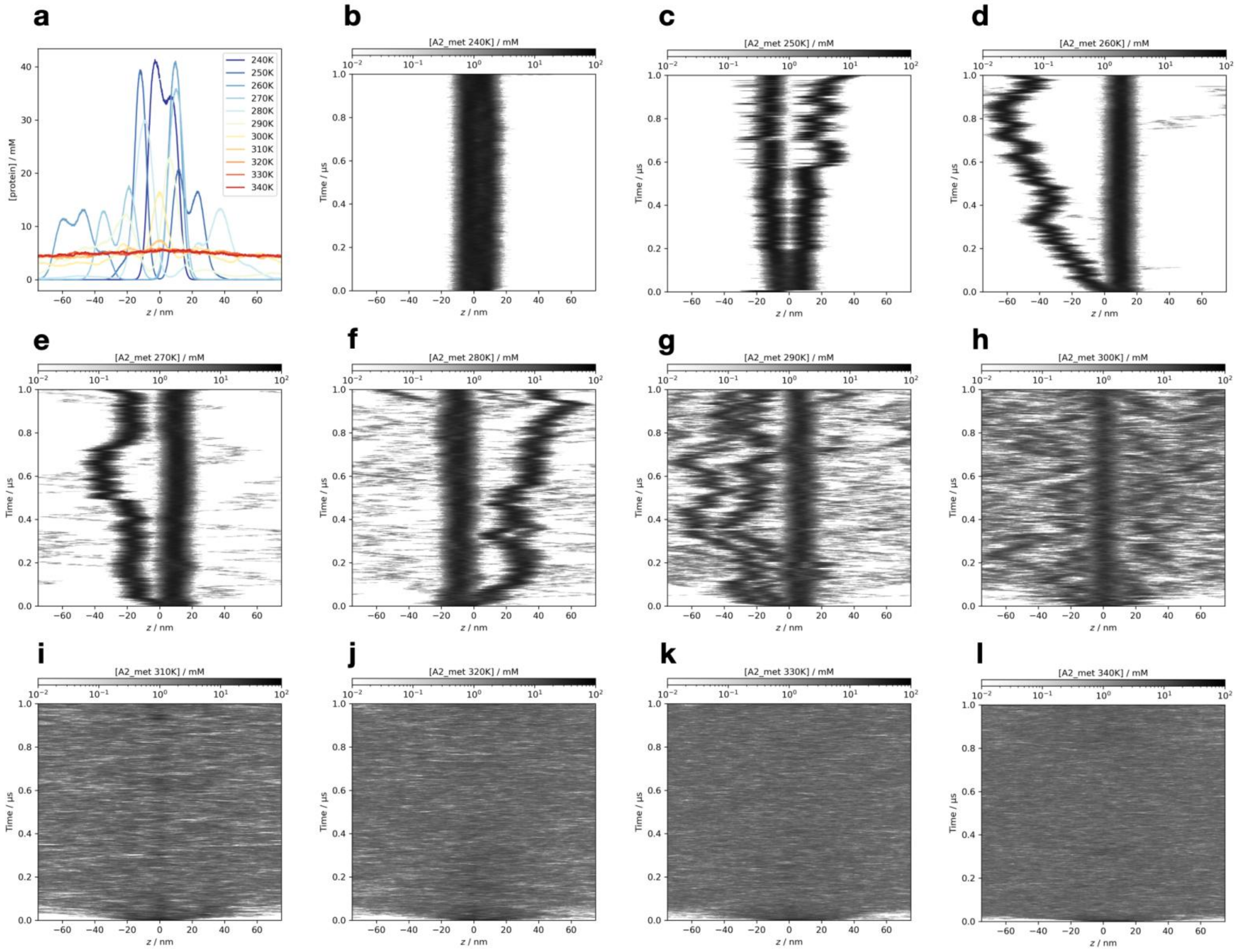


**Figure S23.** The detailed simulation results of the asymmetrically dimethylated hnRNP A2 LCD at all temperatures, simulated using the Slab-PTM patched CALVADOS2 model. (a) The average protein density along the z-axis. (b-l) The time-resolved average protein density along the z-axis from simulation trajectories at temperatures ranging from 240 K to 340 K.

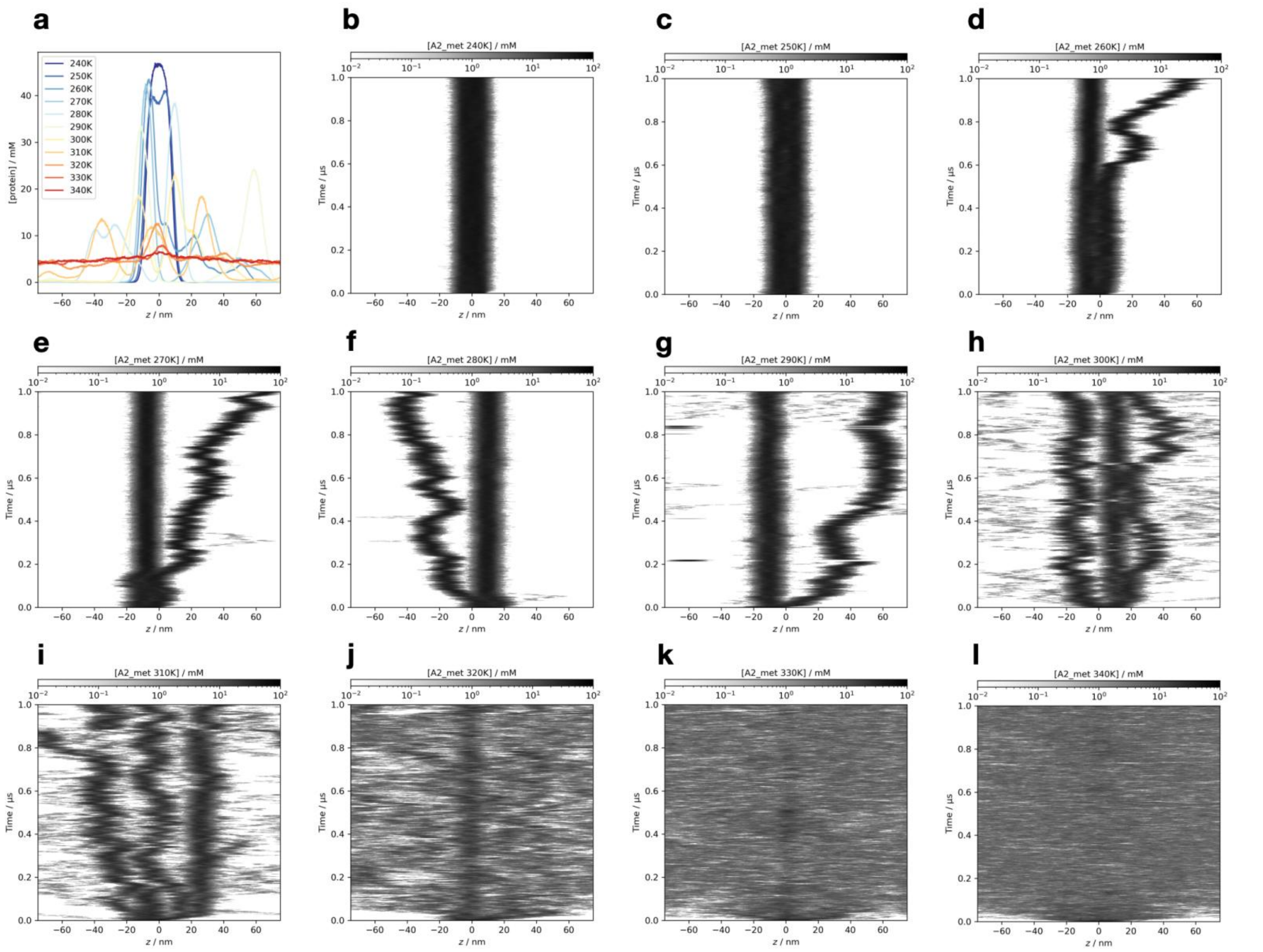


**Figure S24.** The detailed simulation results of the asymmetrically dimethylated hnRNP A2 LCD at all temperatures, simulated using the Slab-PTM patched Mpipi model. (a) The average protein density along the z-axis. (b-l) The time-resolved average protein density along the z-axis from simulation trajectories at temperatures ranging from 240 K to 340 K.

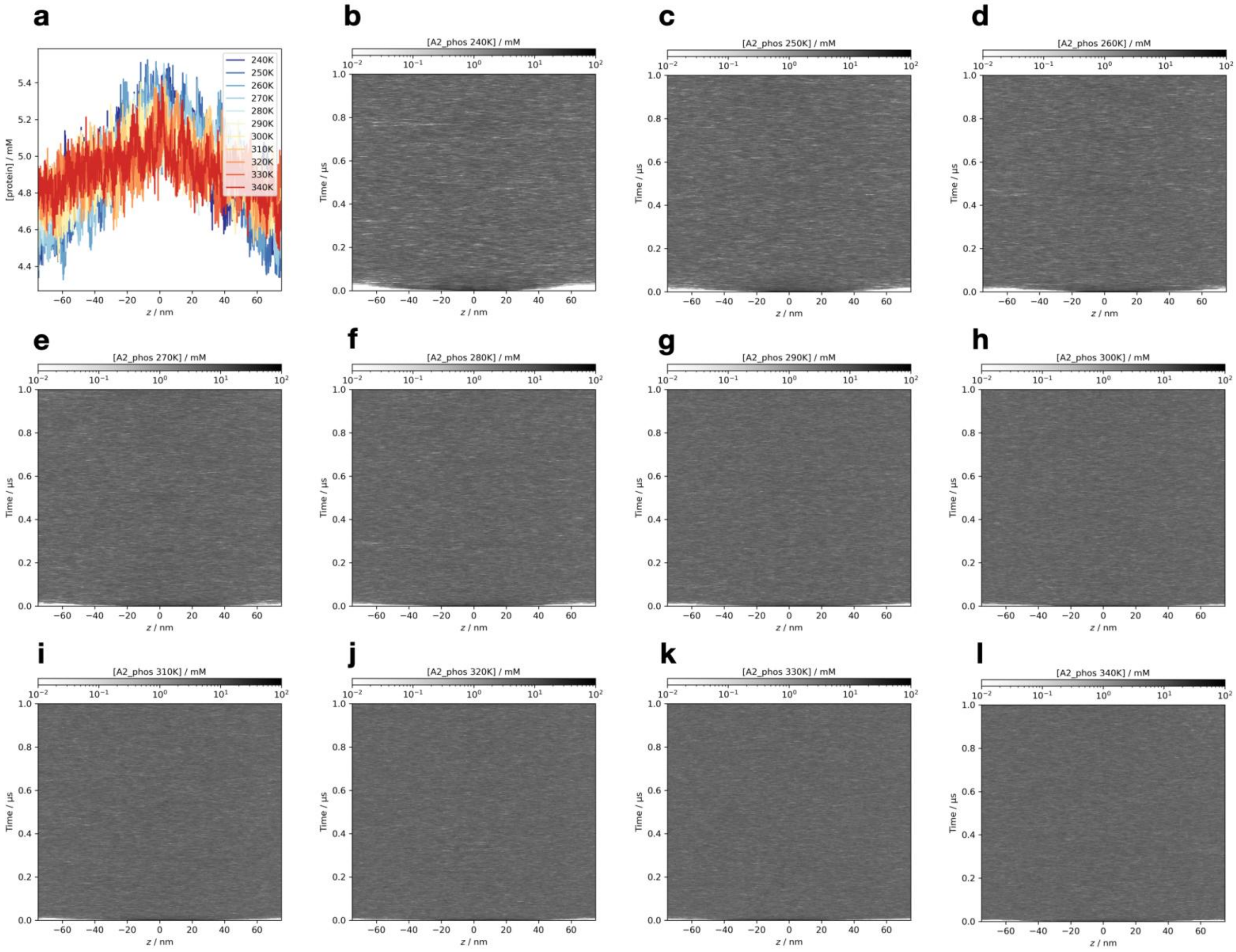


**Figure S25.** The detailed simulation results of the phosphorylated hnRNP A2 LCD at all temperatures, simulated using the Slab-PTM patched CALVADOS2 model. (a) The average protein density along the z-axis. (b-l) The time-resolved average protein density along the z-axis from simulation trajectories at temperatures ranging from 240 K to 340 K.

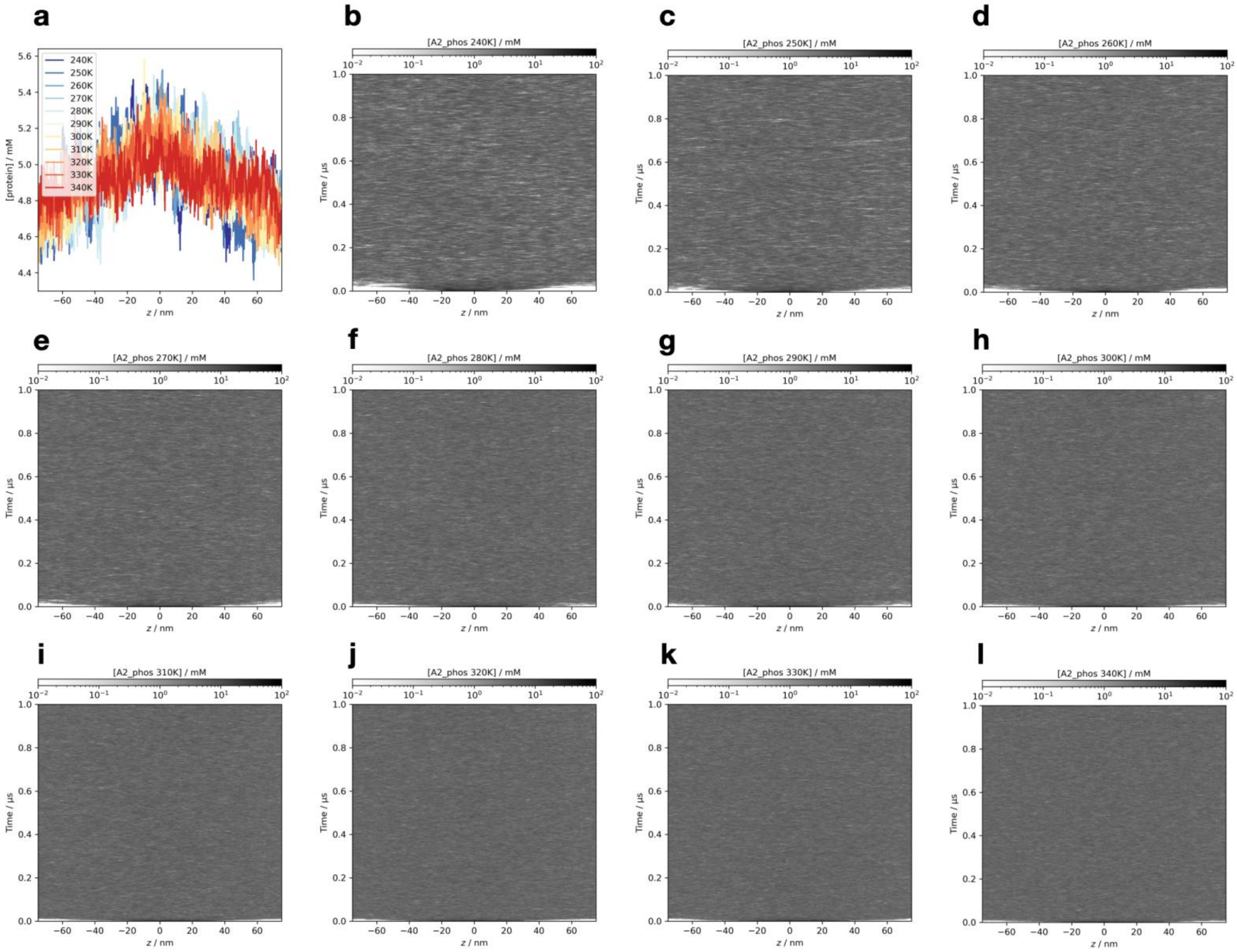


**Figure S26.** The detailed simulation results of the phosphorylated hnRNP A2 LCD at all temperatures, simulated using the Slab-PTM patched Mpipi model. (a) The average protein density along the z-axis. (b-l) The time-resolved average protein density along the z-axis from simulation trajectories at temperatures ranging from 240 K to 340 K.

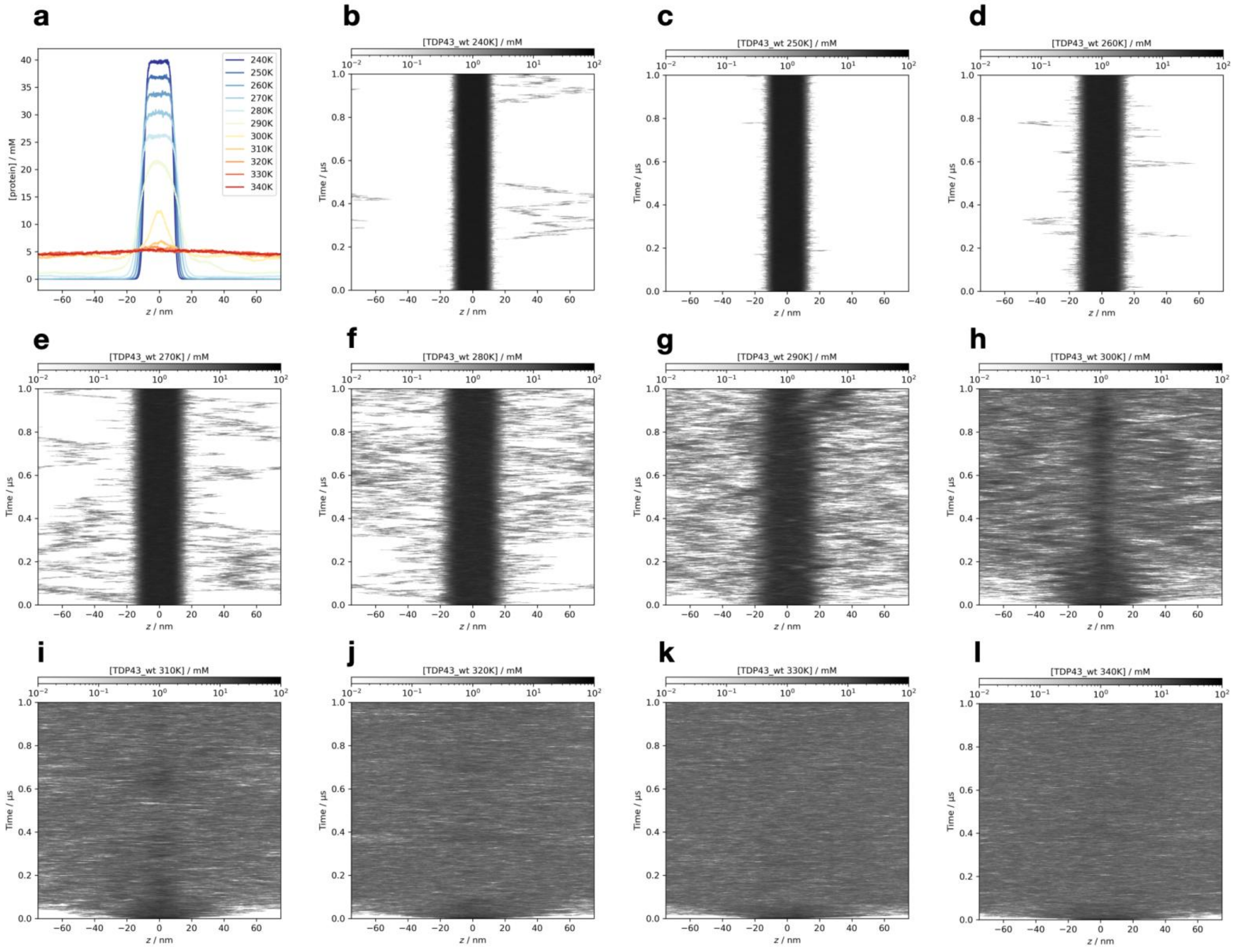

**Figure S27.** The detailed simulation results of the non-PTM TDP43 LCD at all temperatures, simulated using the Slab-PTM patched CALVADOS2 model. (a) The average protein density along the z-axis. (b-l) The time-resolved average protein density along the z-axis from simulation trajectories at temperatures ranging from 240 K to 340 K.

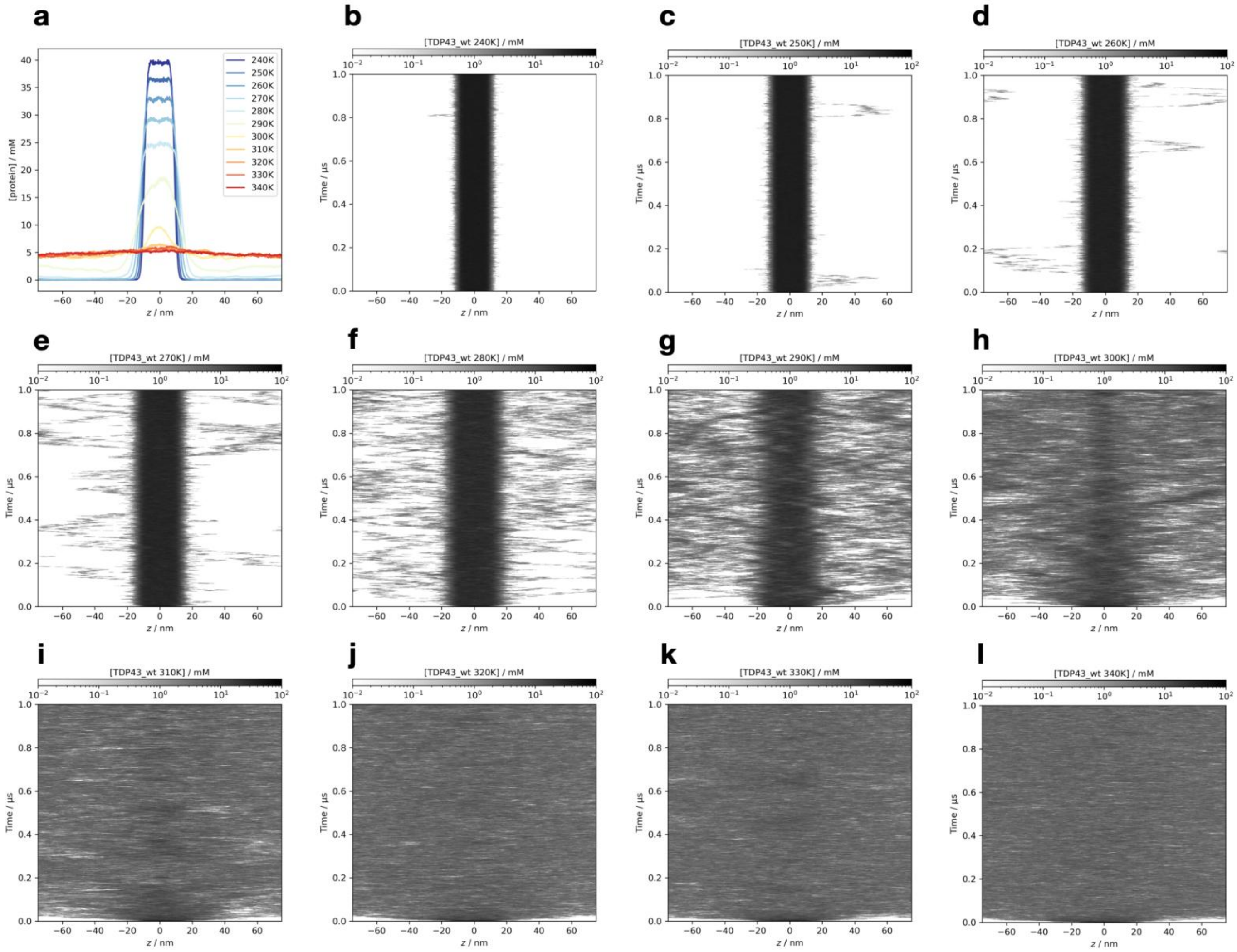


**Figure S28.** The detailed simulation results of the non-PTM TDP43 LCD at all temperatures, simulated using the Slab-PTM patched Mpipi model. (a) The average protein density along the z-axis. (b-l) The time-resolved average protein density along the z-axis from simulation trajectories at temperatures ranging from 240 K to 340 K.

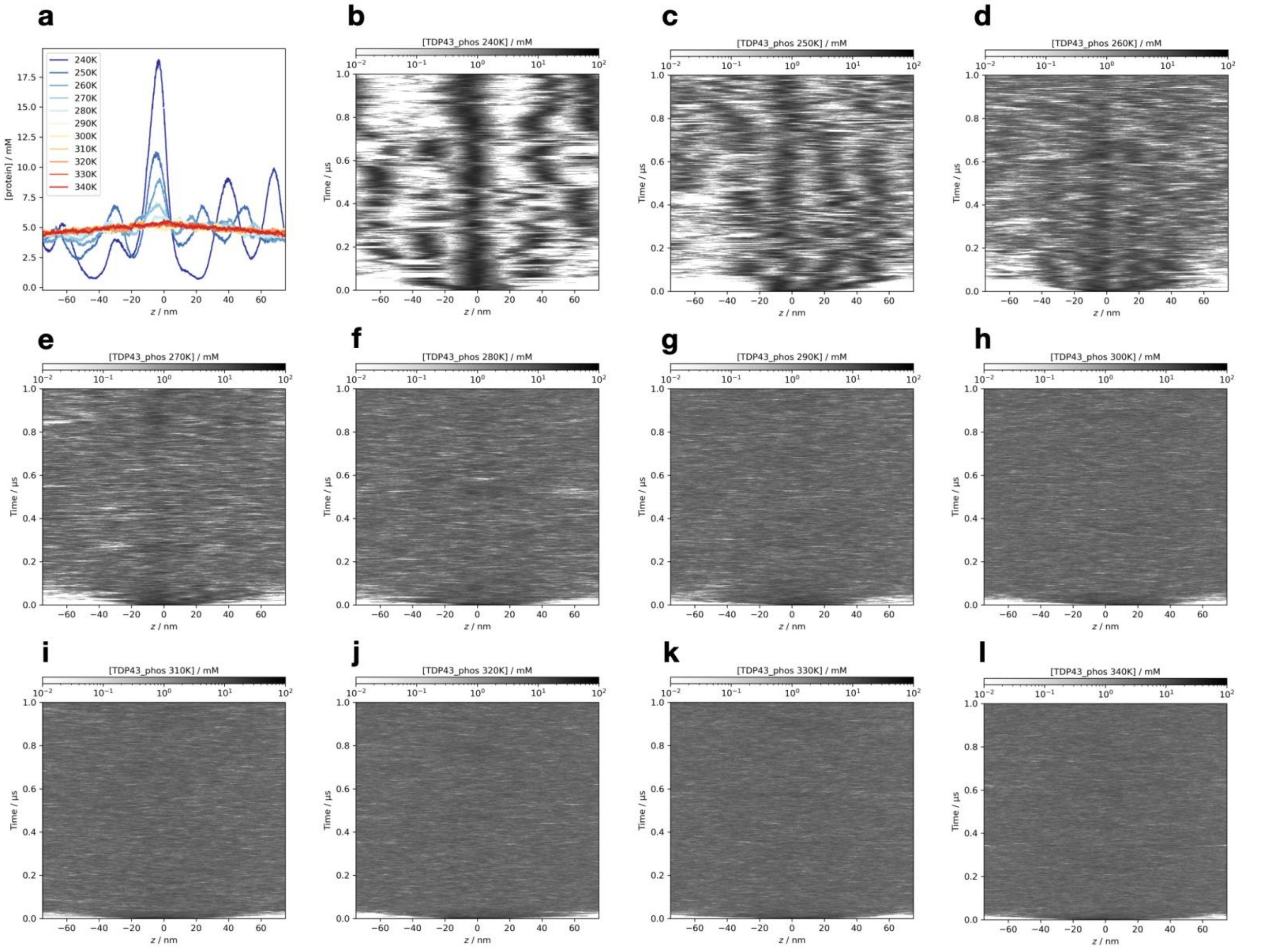


**Figure S29.** The detailed simulation results of the phosphorylated TDP43 LCD at all temperatures, simulated using the Slab-PTM patched CALVADOS2 model. (a) The average protein density along the z-axis. (b-l) The time-resolved average protein density along the z-axis from simulation trajectories at temperatures ranging from 240 K to 340 K.

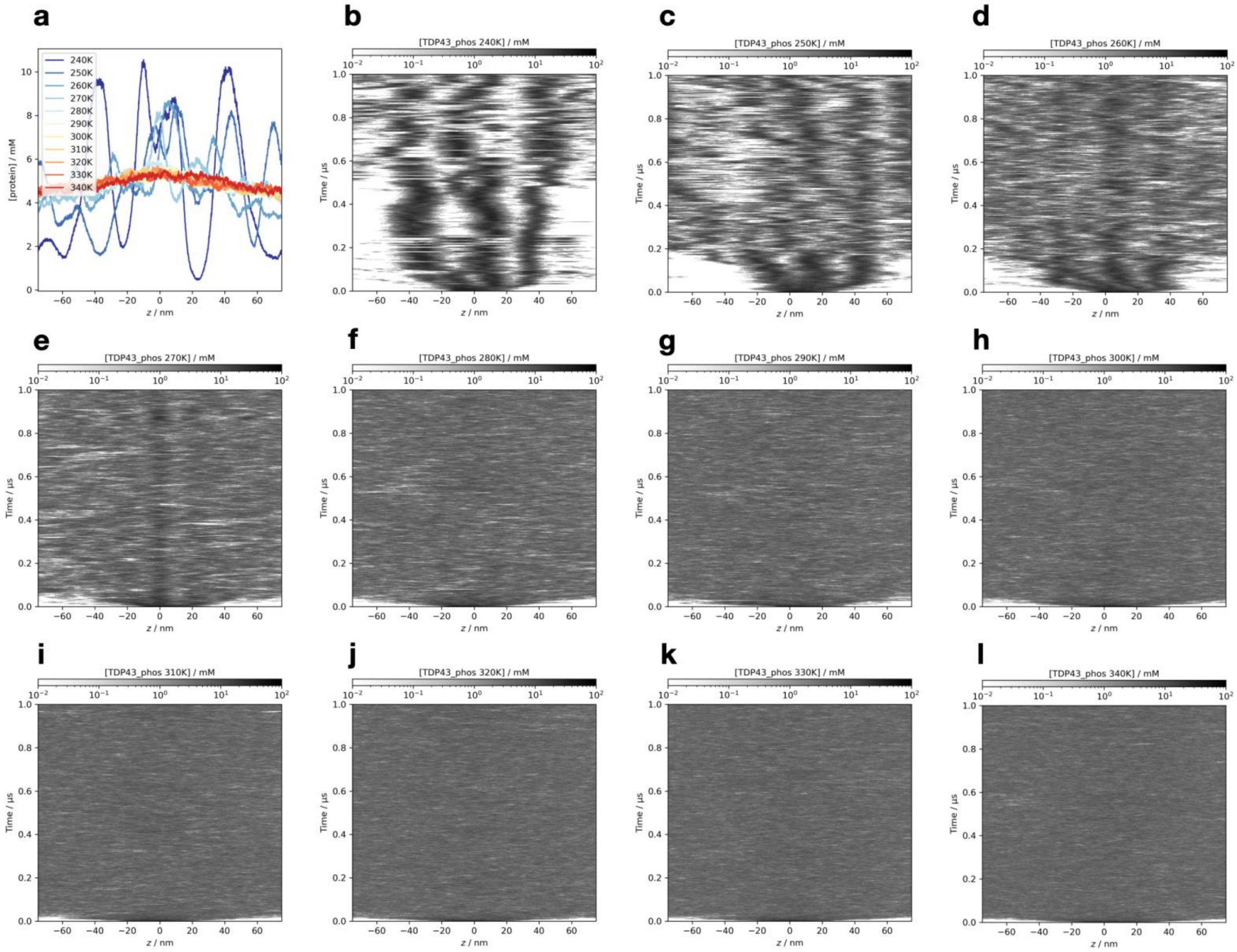


**Figure S30.** The detailed simulation results of the phosphorylated TDP43 LCD at all temperatures, simulated using the Slab-PTM patched Mpipi model. (a) The average protein density along the z-axis. (b-l) The time-resolved average protein density along the z-axis from simulation trajectories at temperatures ranging from 240 K to 340 K.

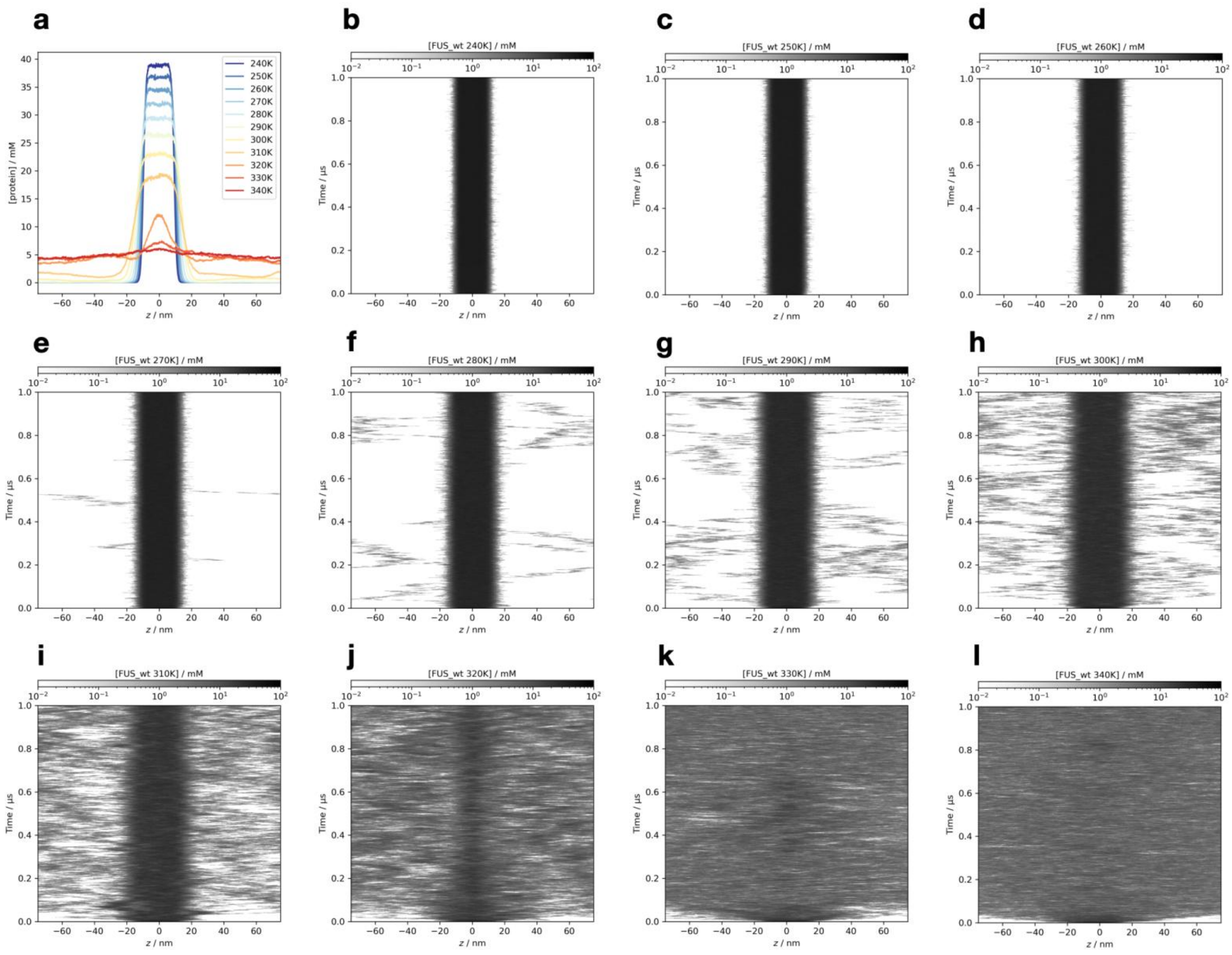


**Figure S31.** The detailed simulation results of the non-PTM FUS N-terminal domain at all temperatures, simulated using the Slab-PTM patched CALVADOS2 model. (a) The average protein density along the z-axis. (b-l) The time-resolved average protein density along the z-axis from simulation trajectories at temperatures ranging from 240 K to 340 K.

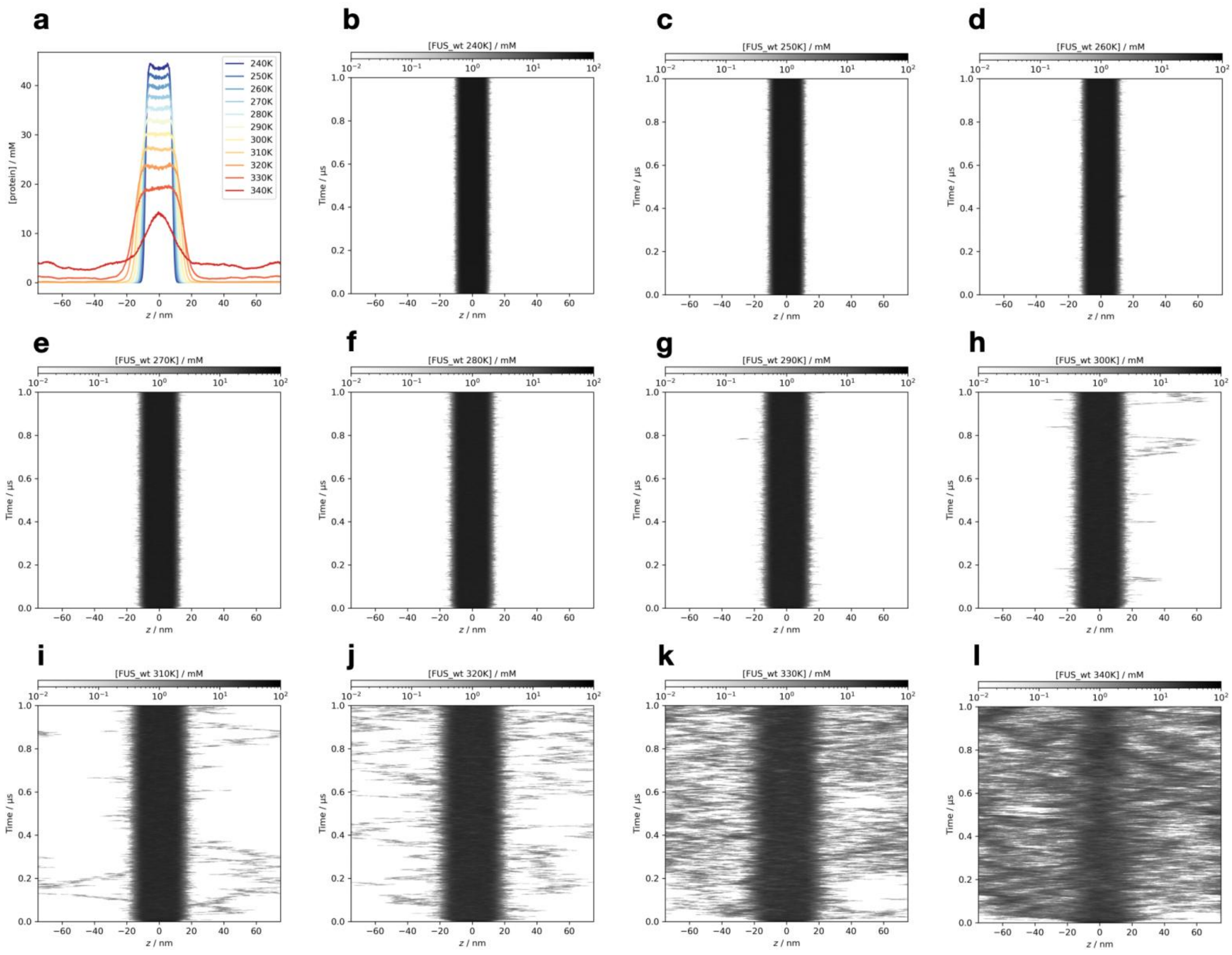


**Figure S32.** The detailed simulation results of the FUS N-terminal domain at all temperatures, simulated using the Slab-PTM patched Mpipi model. (a) The average protein density along the z-axis. (b-l) The time-resolved average protein density along the z-axis from simulation trajectories at temperatures ranging from 240 K to 340 K.

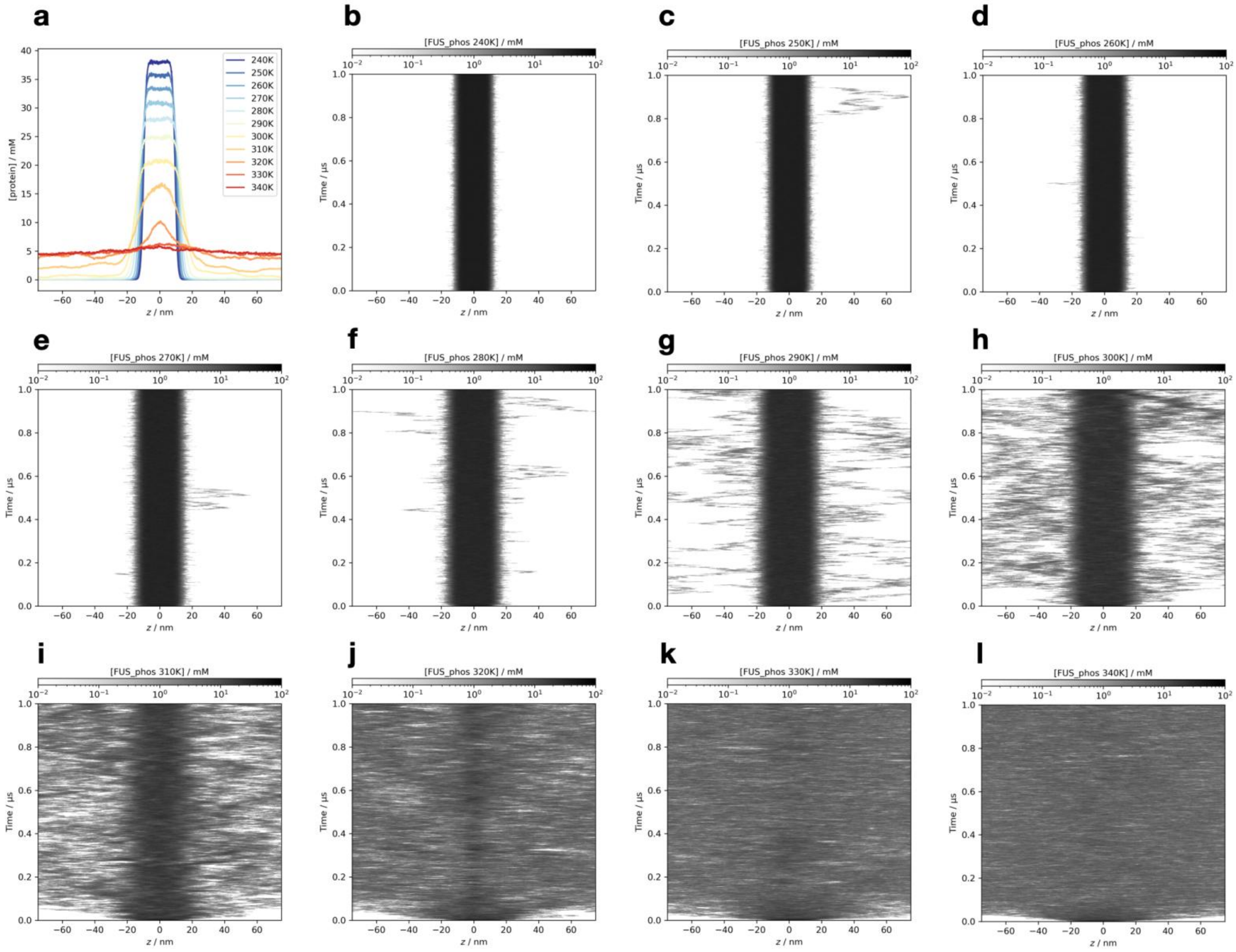

**Figure S33.** The detailed simulation results of the phosphorylated FUS N-terminal domain at all temperatures, simulated using the Slab-PTM patched CALVADOS2 model. (a) The average protein density along the z-axis. (b-l) The time-resolved average protein density along the z-axis from simulation trajectories at temperatures ranging from 240 K to 340 K.

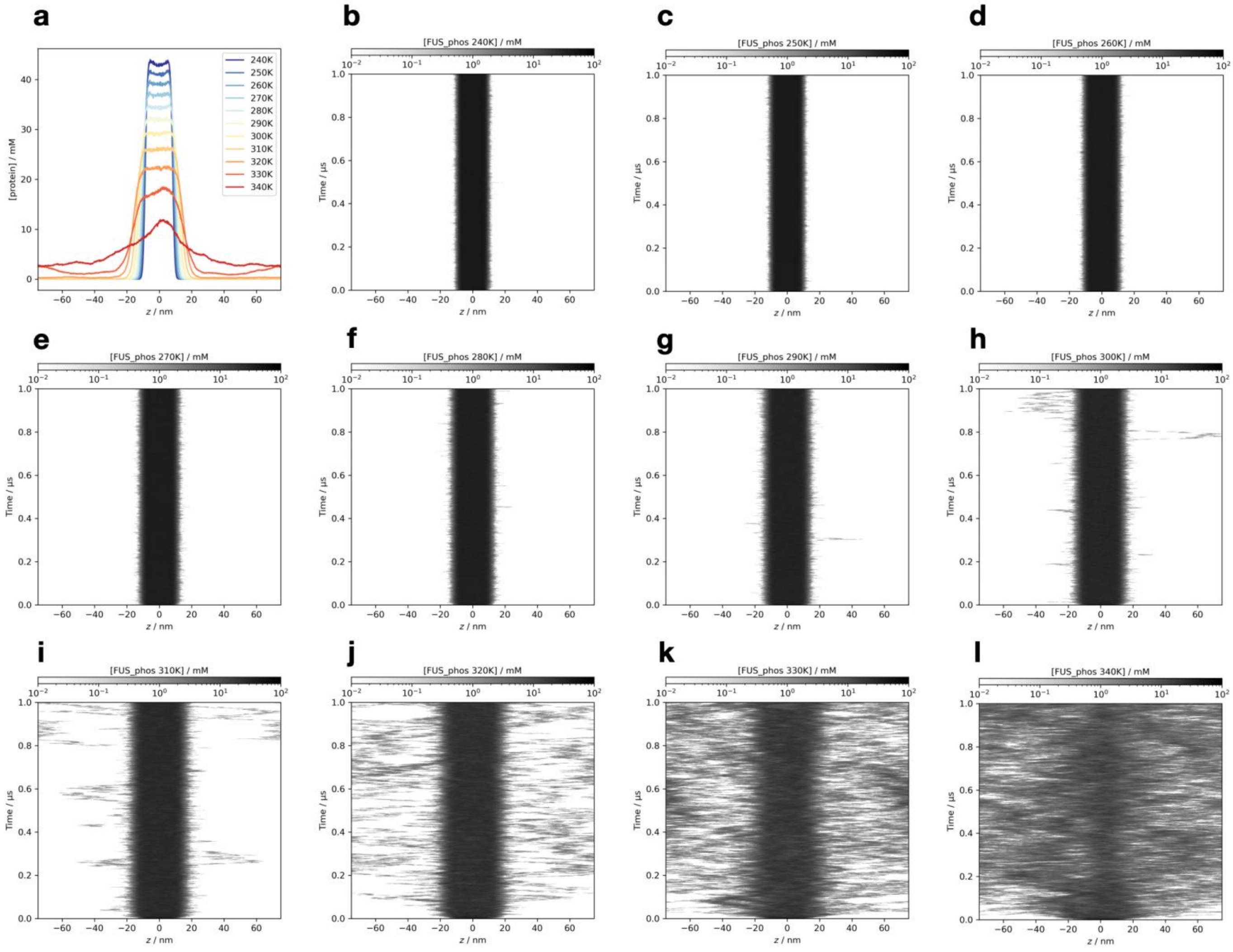

**Figure S34.** The detailed simulation results of the phosphorylated FUS N-terminal domain at all temperatures, simulated using the Slab-PTM patched Mpipi model. (a) The average protein density along the z-axis. (b-l) The time-resolved average protein density along the z-axis from simulation trajectories at temperatures ranging from 240 K to 340 K.

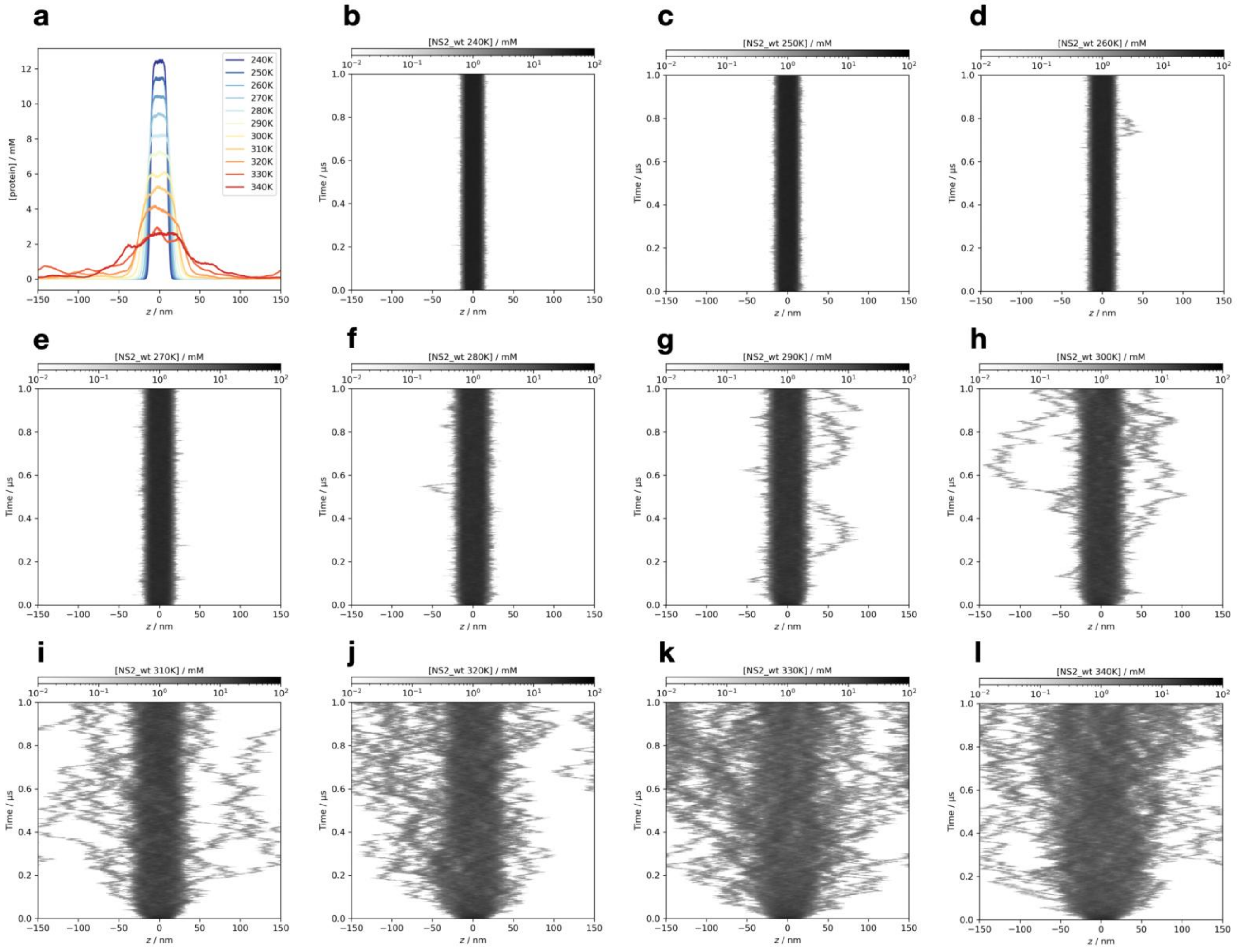


**Figure S35.** The detailed simulation results of the non-PTM NS2 at all temperatures, simulated using the Slab-PTM patched CALVADOS2 model. (a) The average protein density along the z-axis. (b-l) The time-resolved average protein density along the z-axis from simulation trajectories at temperatures ranging from 240 K to 340 K.

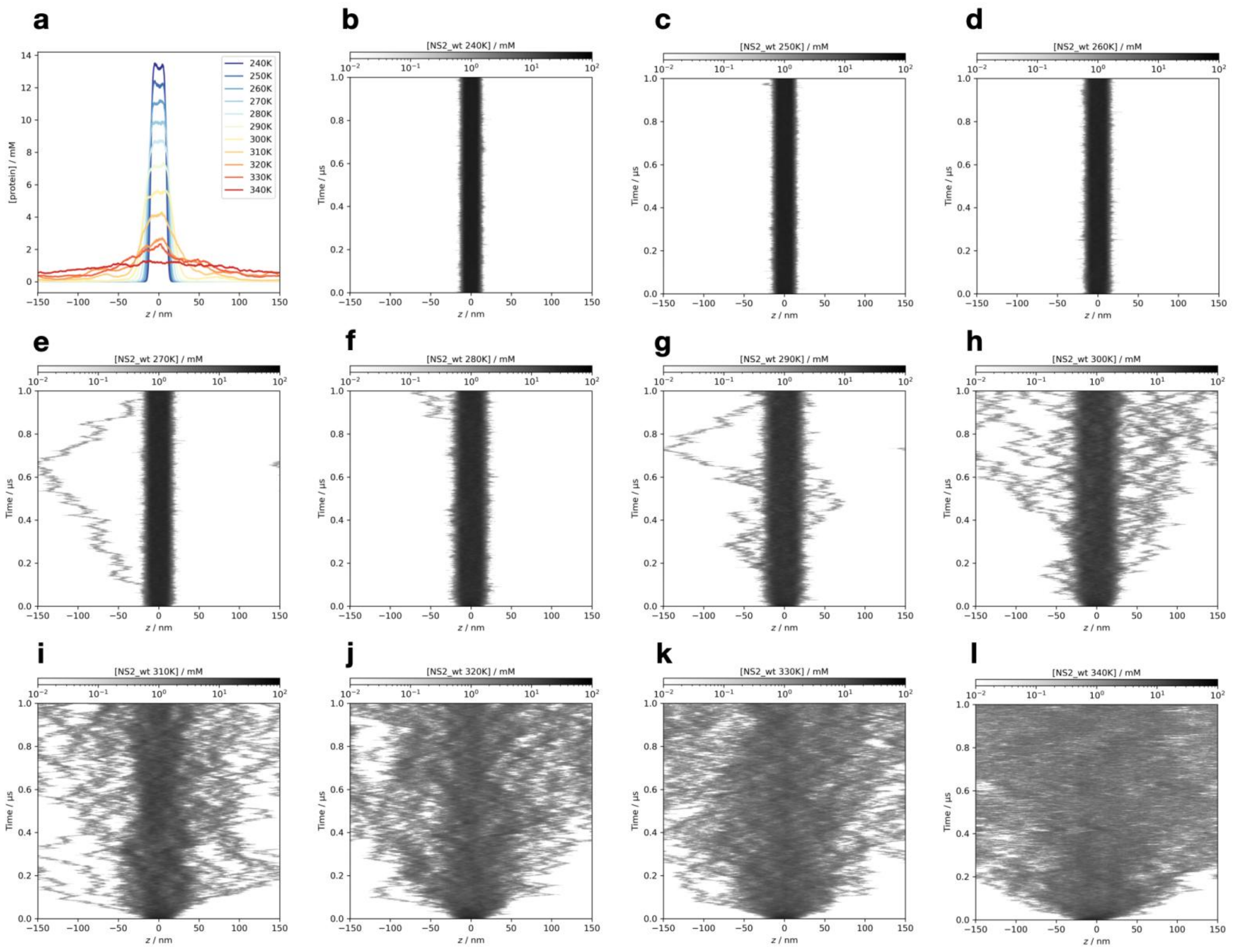


**Figure S36.** The detailed simulation results of the non-PTM NS2 at all temperatures, simulated using the Slab-PTM patched Mpipi model. (a) The average protein density along the z-axis. (b-l) The time-resolved average protein density along the z-axis from simulation trajectories at temperatures ranging from 240 K to 340 K.

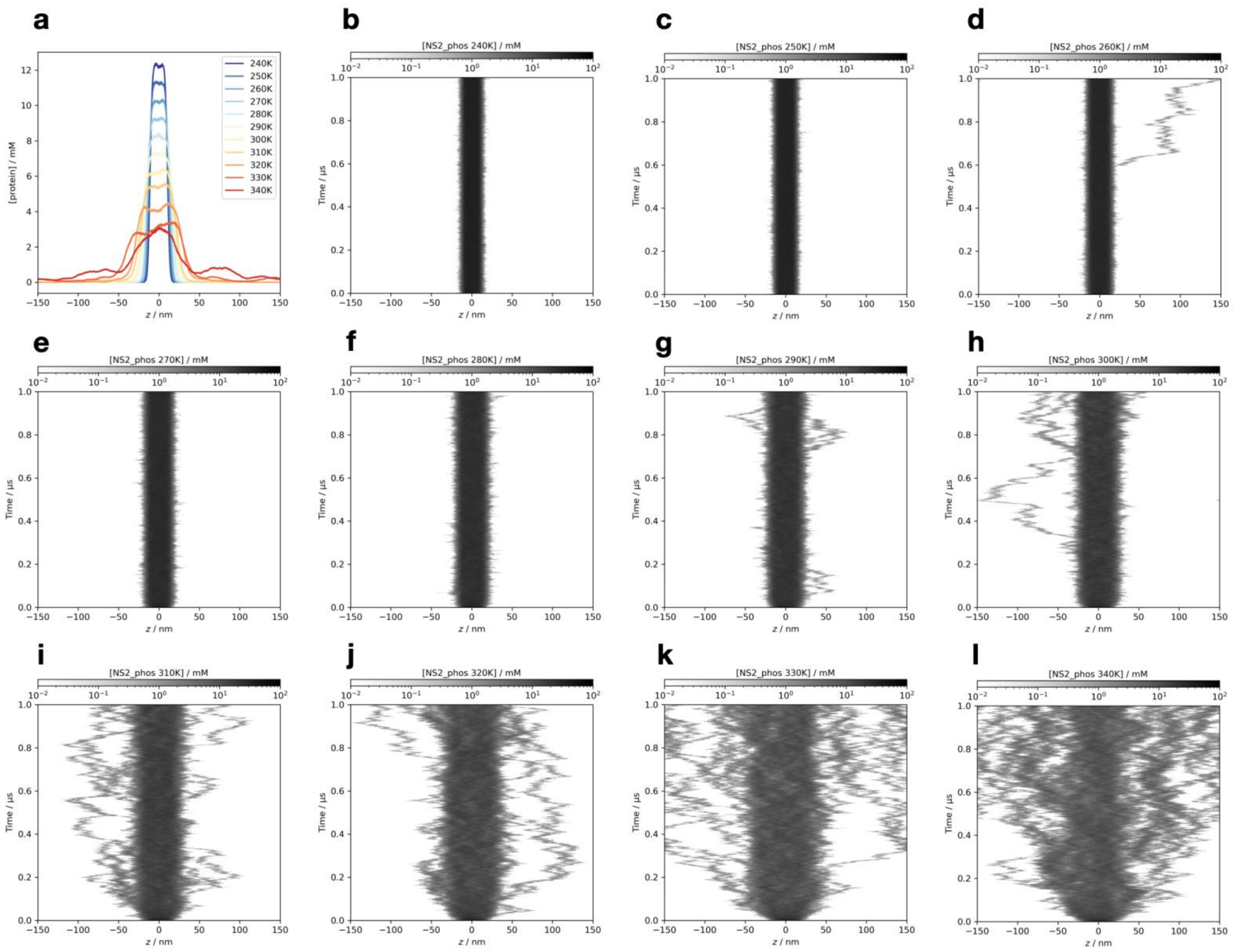


**Figure S37.** The detailed simulation results of the phosphorylated NS2 at all temperatures, simulated using the Slab-PTM patched CALVADOS2 model. (a) The average protein density along the z-axis. (b-l) The time-resolved average protein density along the z-axis from simulation trajectories at temperatures ranging from 240 K to 340 K

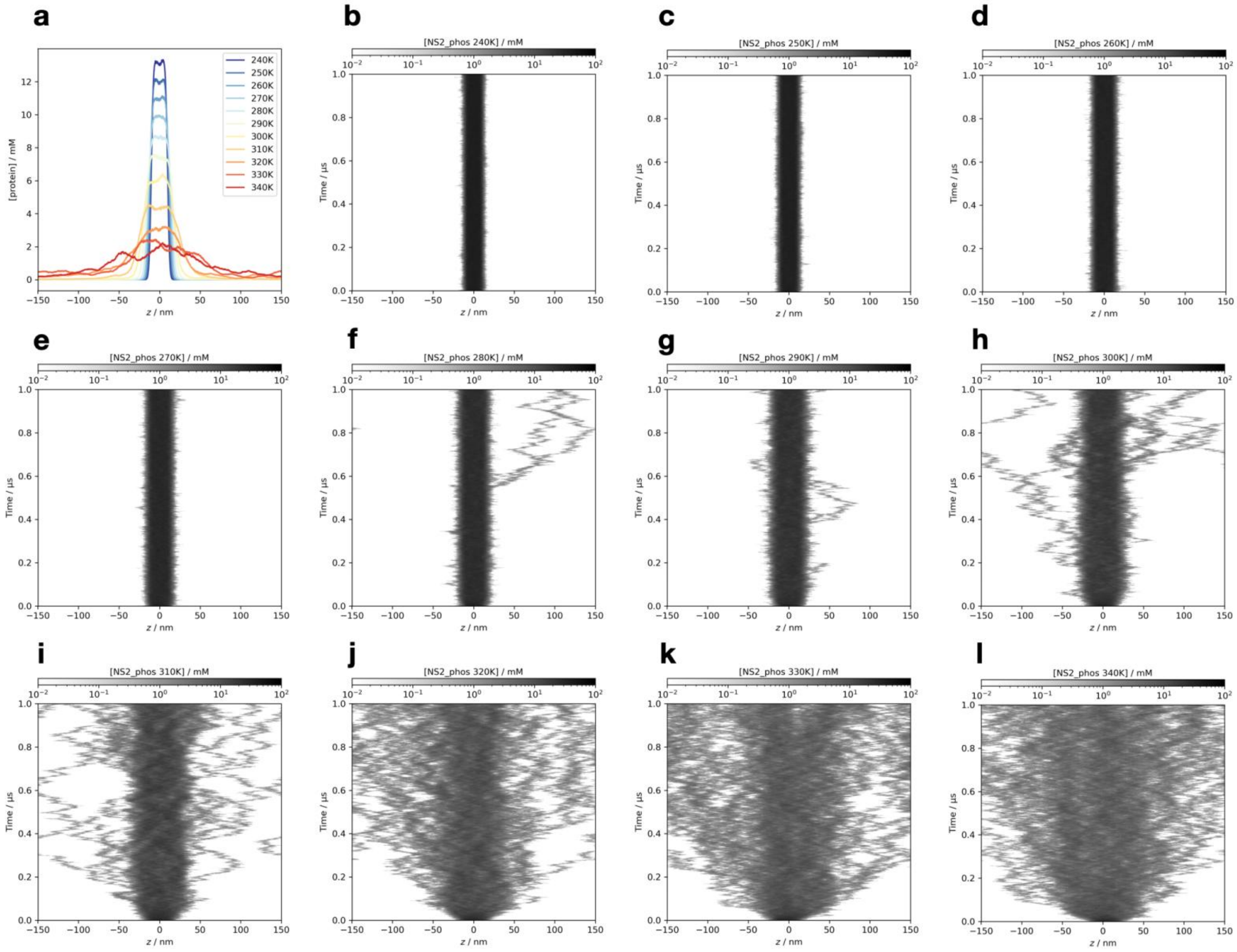

**Figure S38.** The detailed simulation results of the phosphorylated NS2 at all temperatures, simulated using the Slab-PTM patched Mpipi model. (a) The average protein density along the z-axis. (b-l) The time-resolved average protein density along the z-axis from simulation trajectories at temperatures ranging from 240 K to 340 K.

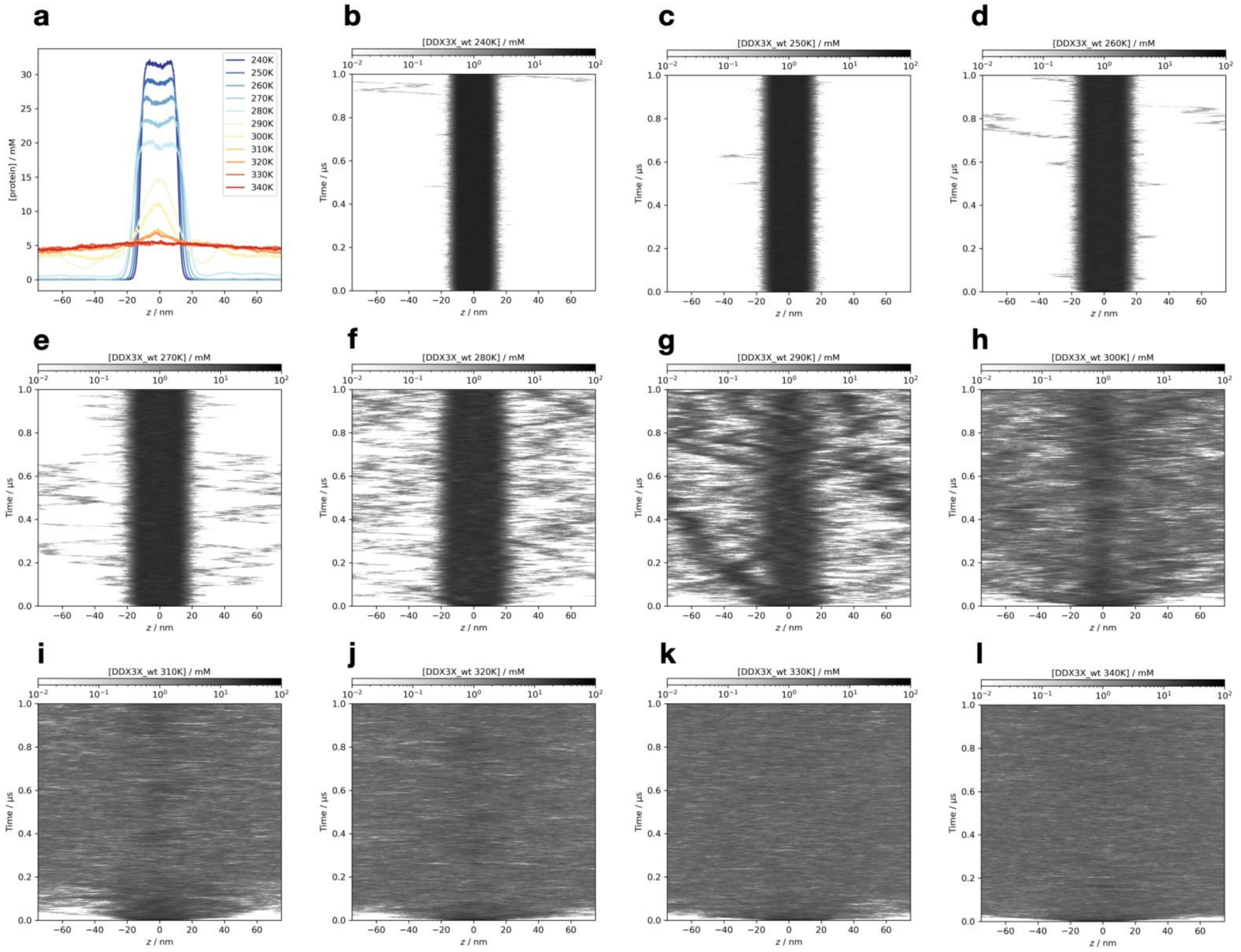


**Figure S39.** The detailed simulation results of the non-PTM DDX3X IDR1 domain at all temperatures, simulated using the Slab-PTM patched CALVADOS2 model. (a) The average protein density along the z-axis. (b-l) The time-resolved average protein density along the z-axis from simulation trajectories at temperatures ranging from 240 K to 340 K.

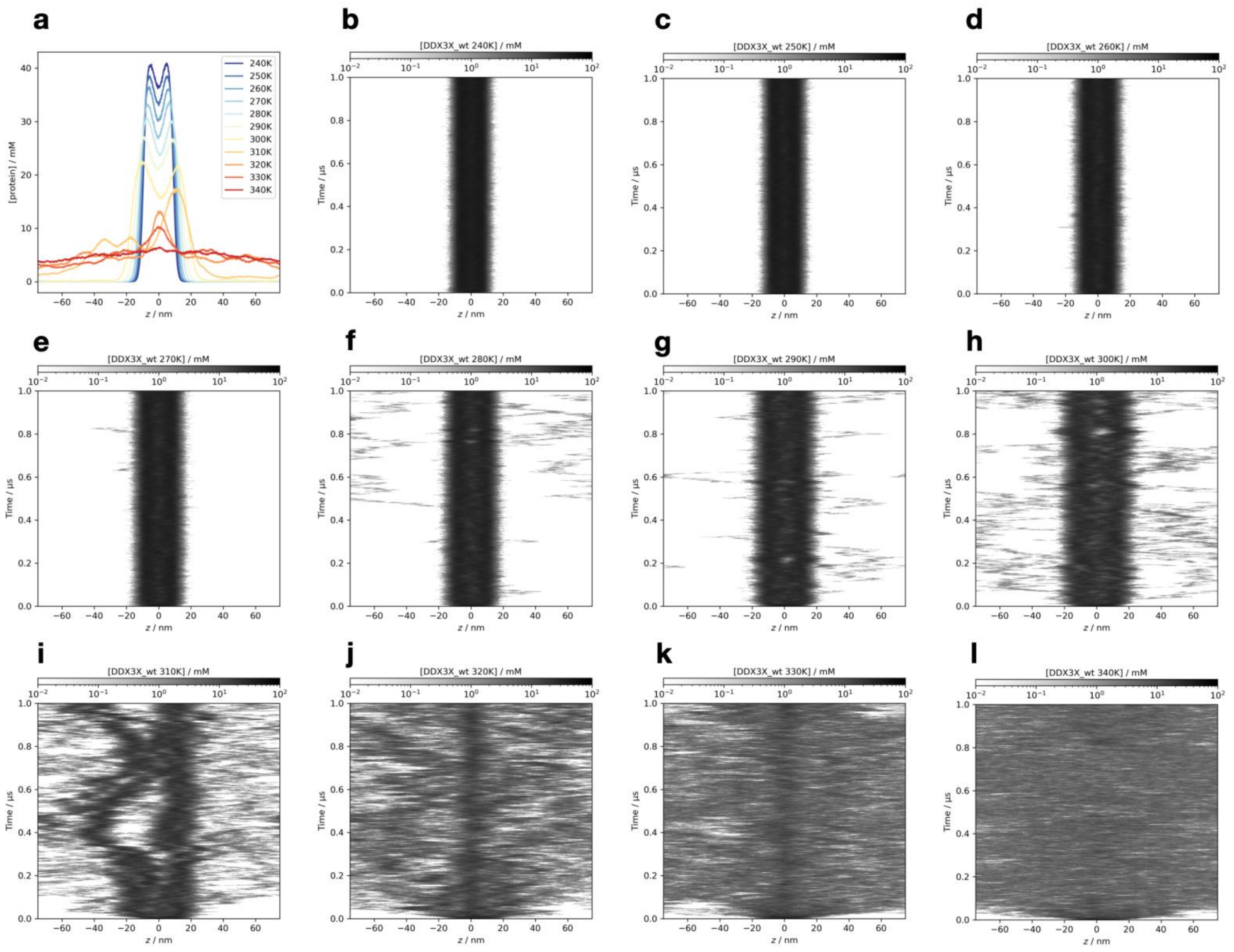


**Figure S40.** The detailed simulation results of the non-PTM DDX3X IDR1 domain at all temperatures, simulated using the Slab-PTM patched Mpipi model. (a) The average protein density along the z-axis. (b-l) The time-resolved average protein density along the z-axis from simulation trajectories at temperatures ranging from 240 K to 340 K.

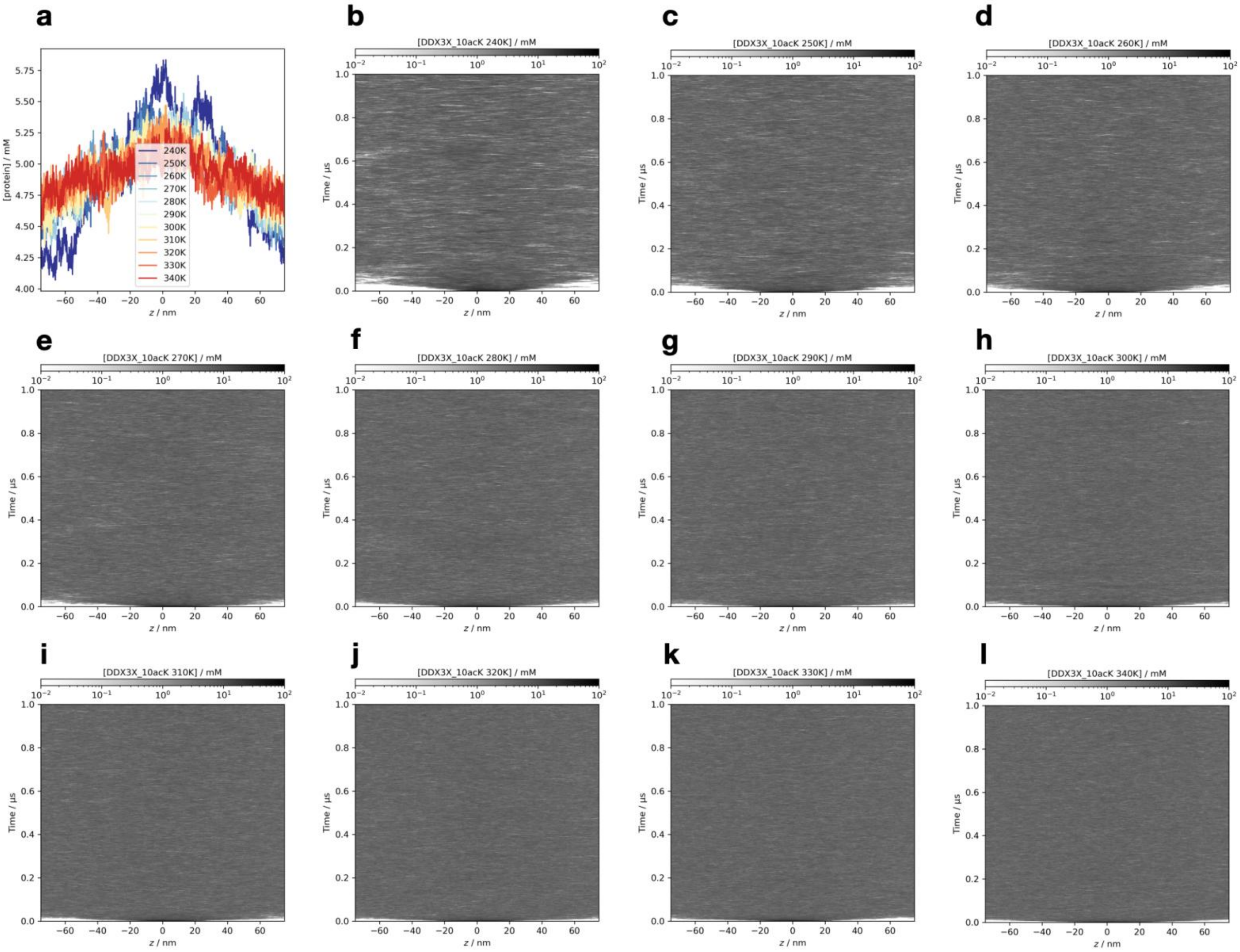


**Figure S41.** The detailed simulation results of the acetylated DDX3X IDR1 domain at all temperatures, simulated using the Slab-PTM patched CALVADOS2 model. (a) The average protein density along the z-axis. (b-l) The time-resolved average protein density along the z-axis from simulation trajectories at temperatures ranging from 240 K to 340 K.

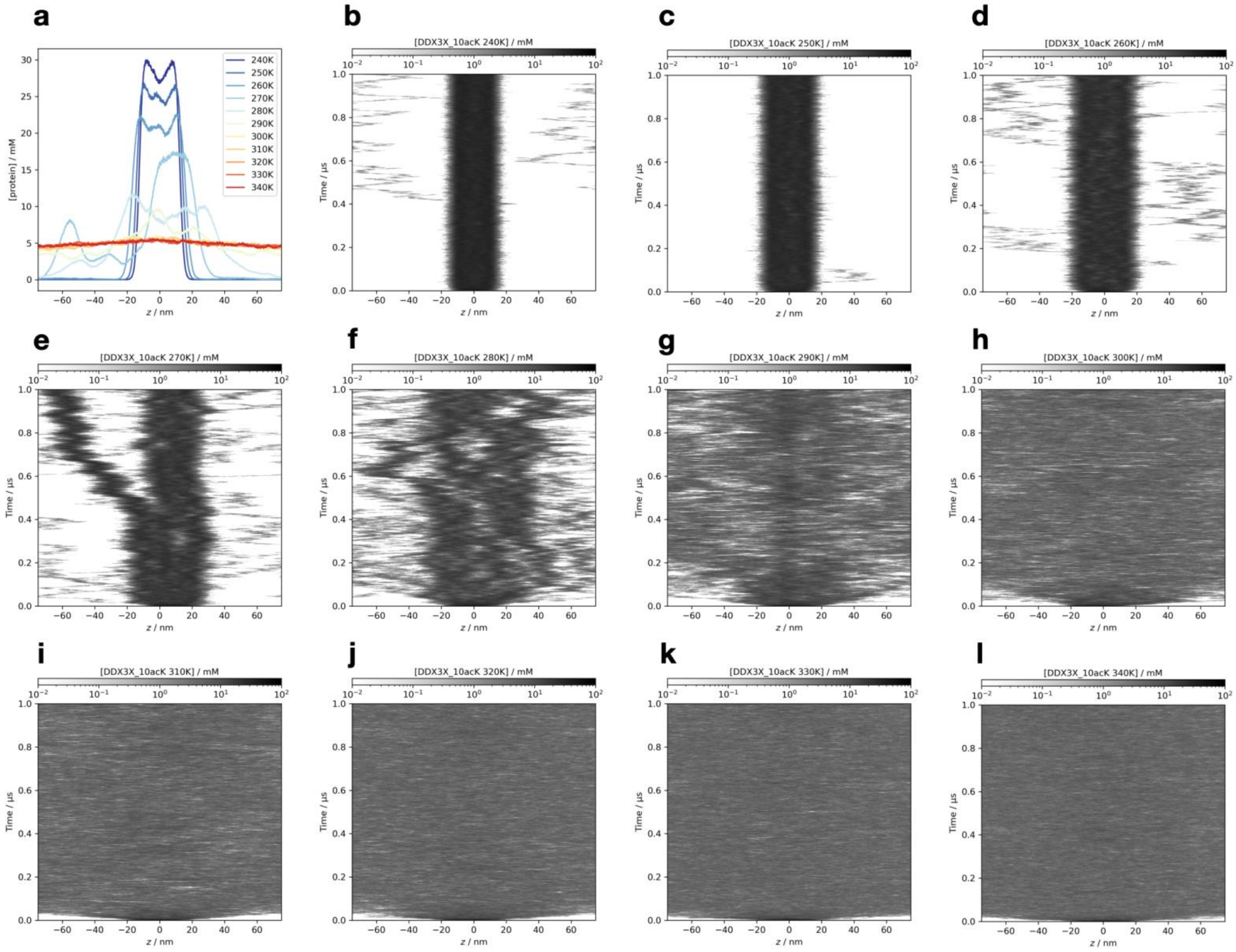


**Figure S42.** The detailed simulation results of the acetylated DDX3X IDR1 domain at all temperatures, simulated using the Slab-PTM patched Mpipi model. (a) The average protein density along the z-axis. (b-l) The time-resolved average protein density along the z-axis from simulation trajectories at temperatures ranging from 240 K to 340 K.

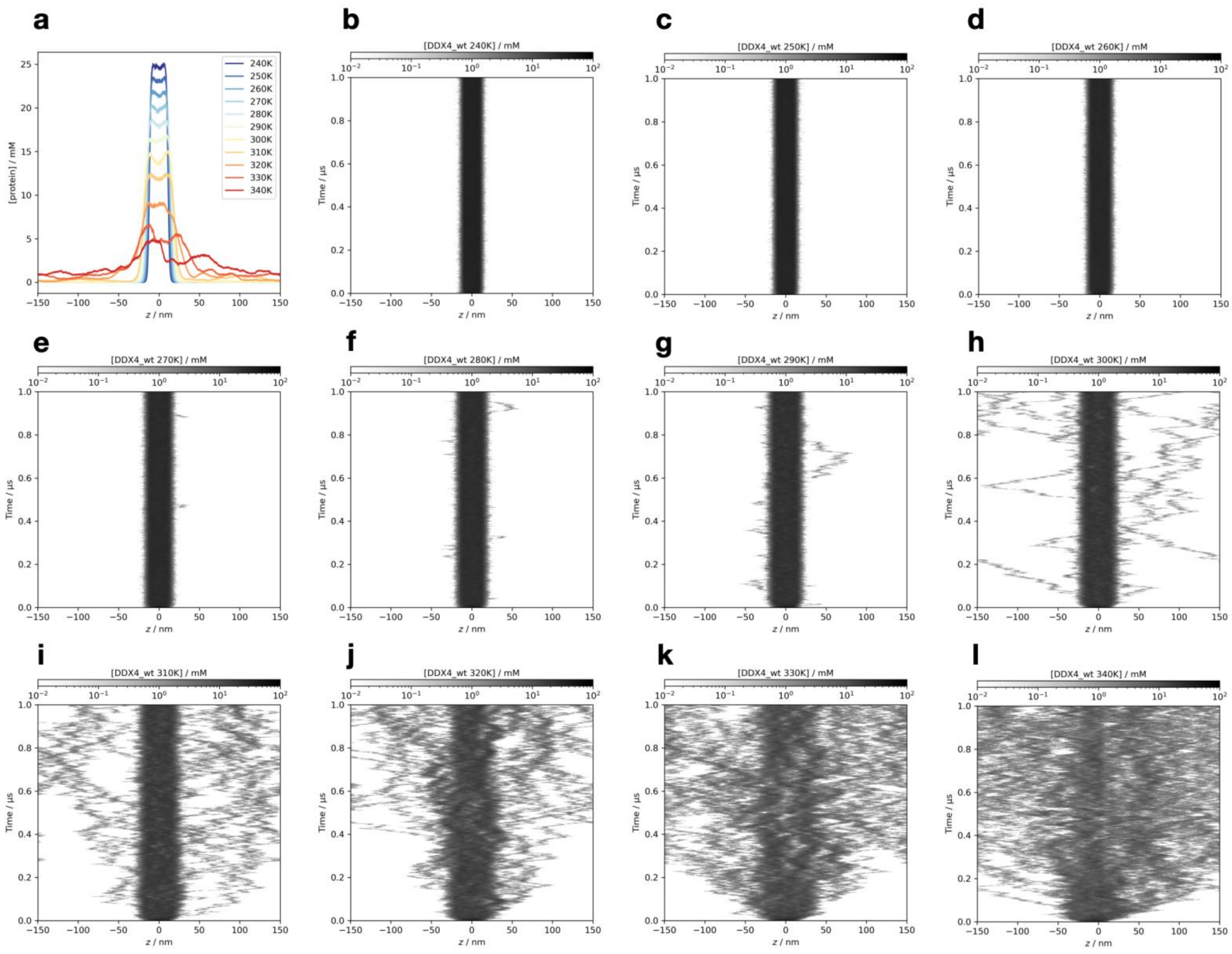


**Figure S43.** The detailed simulation results of the non-PTM DDX4 N-terminal domain at all temperatures, simulated using the Slab-PTM patched CALVADOS2 model. (a) The average protein density along the z-axis. (b-l) The time-resolved average protein density along the z-axis from simulation trajectories at temperatures ranging from 240 K to 340 K.

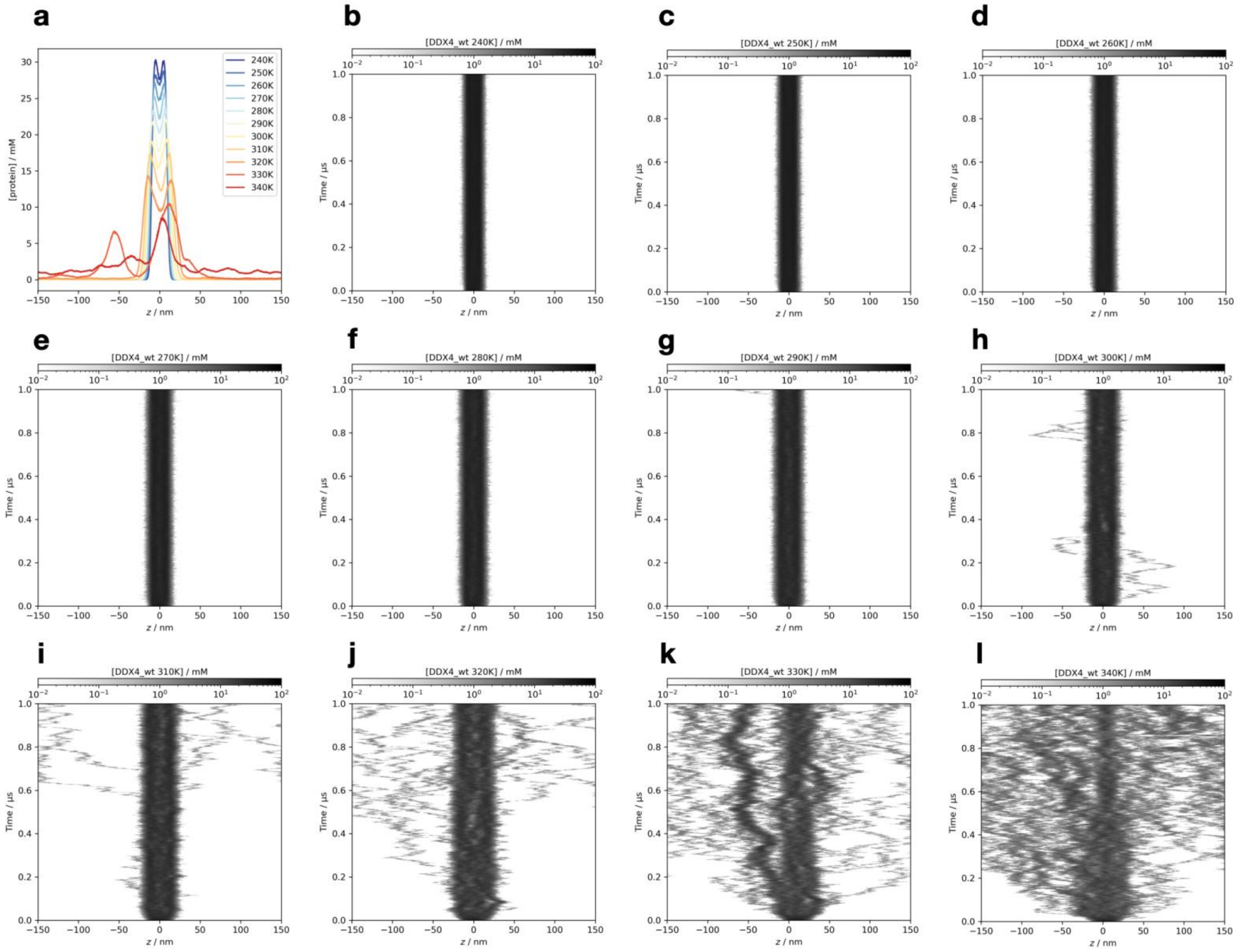

**Figure S44.** The detailed simulation results of the non-PTM DDX4 N-terminal domain at all temperatures, simulated using the Slab-PTM patched Mpipi model. (a) The average protein density along the z-axis. (b-l) The time-resolved average protein density along the z-axis from simulation trajectories at temperatures ranging from 240 K to 340 K.

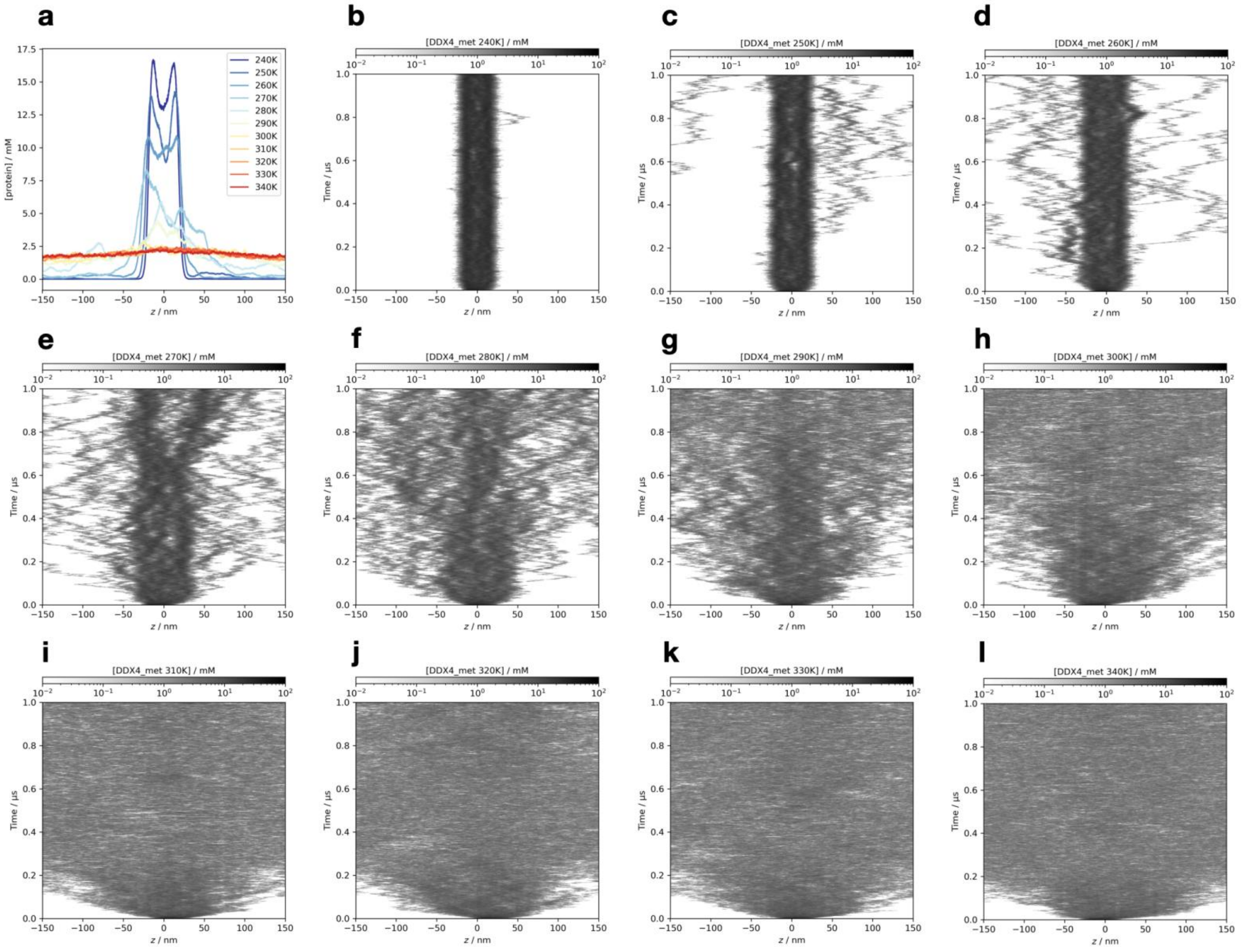


**Figure S45.** The detailed simulation results of the asymmetrically dimethylated DDX4 N-terminal domain at all temperatures, simulated using the Slab-PTM patched CALVADOS2 model. (a) The average protein density along the z-axis. (b-l) The time-resolved average protein density along the z-axis from simulation trajectories at temperatures ranging from 240 K to 340 K.

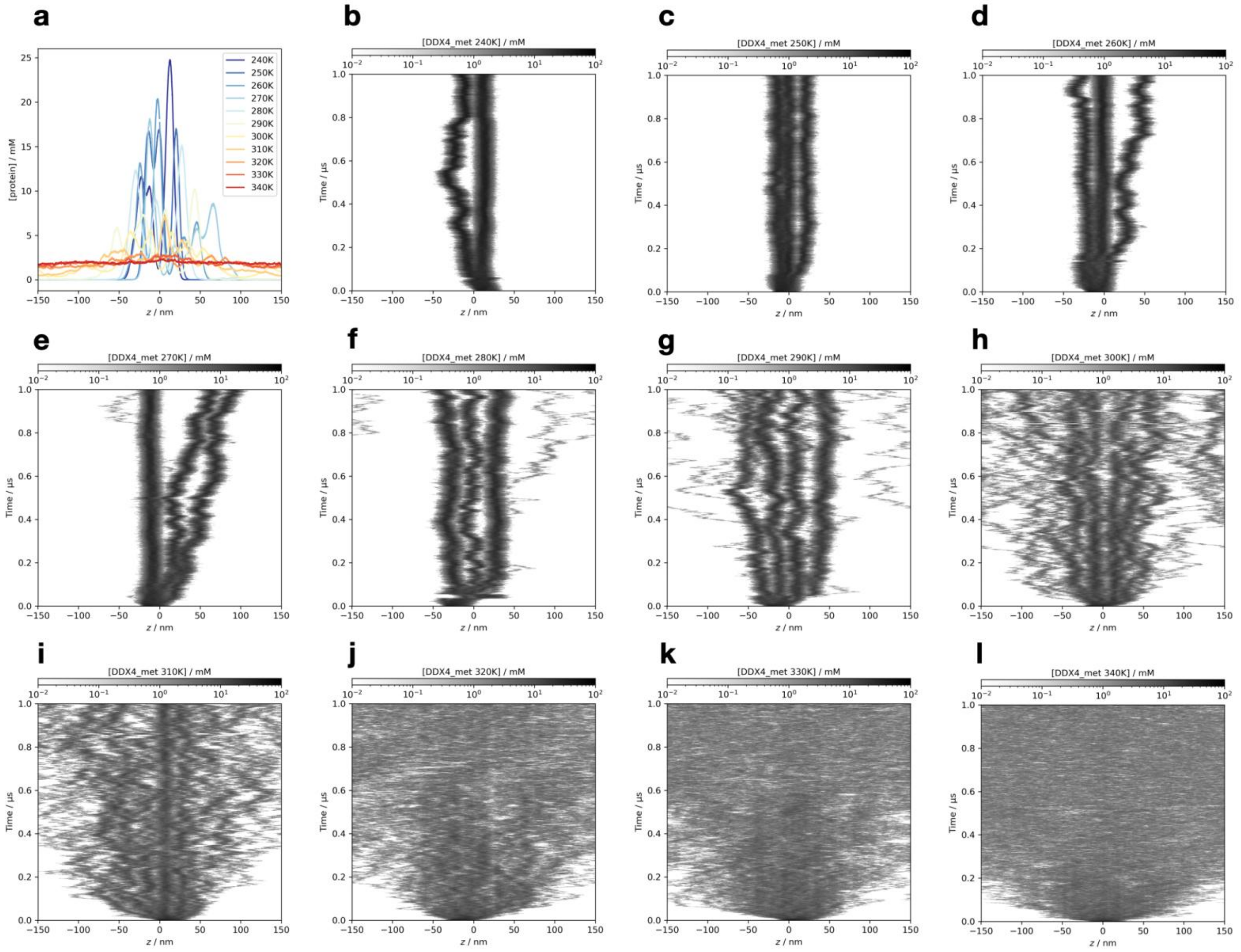


**Figure S46.** The detailed simulation results of the asymmetrically dimethylated DDX4 N-terminal domain at all temperatures, simulated using the Slab-PTM patched Mpipi model. (a) The average protein density along the z-axis. (b-l) The time-resolved average protein density along the z-axis from simulation trajectories at temperatures ranging from 240 K to 340 K.

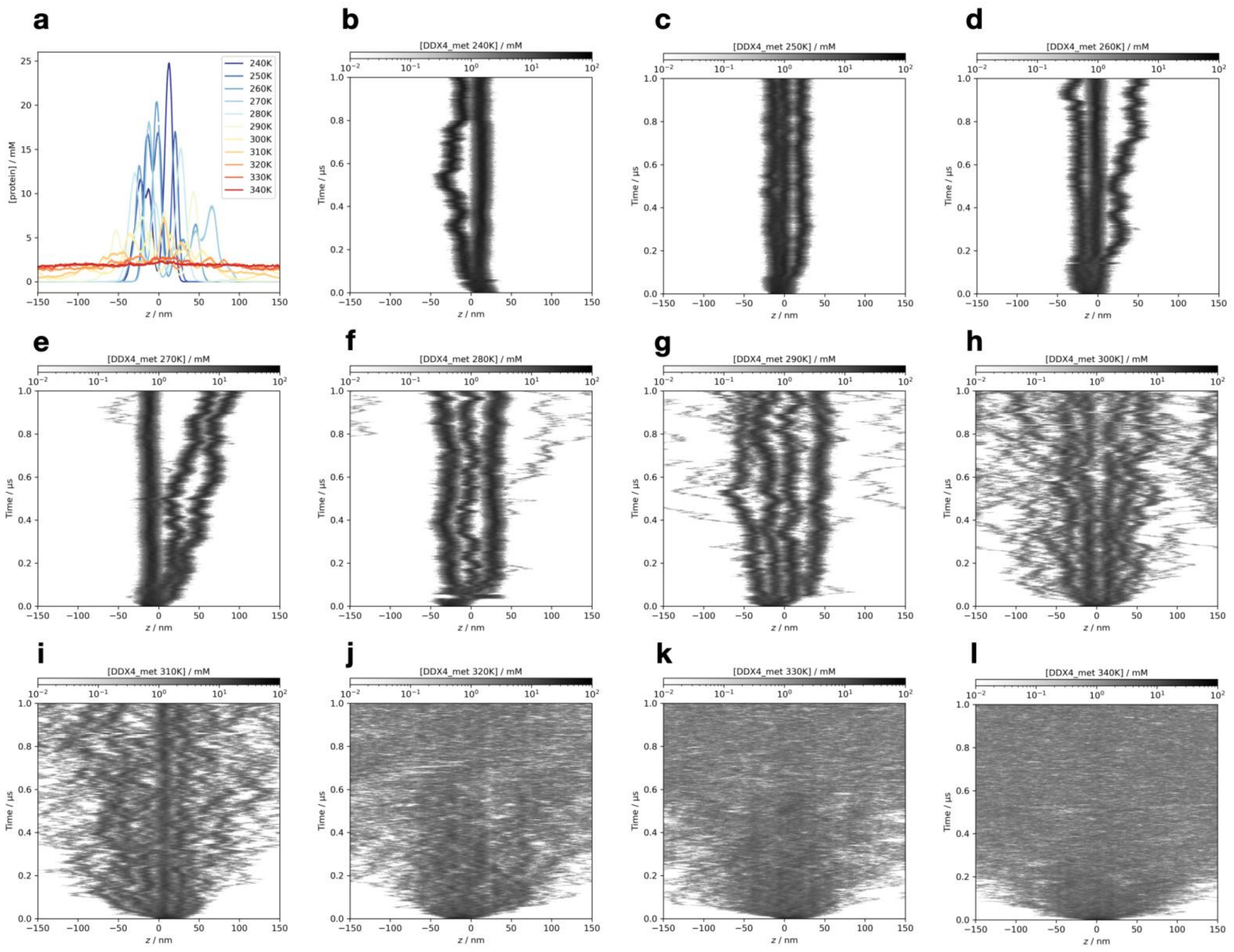


**Figure S46.** The detailed simulation results of the asymmetrically dimethylated DDX4 N-terminal domain at all temperatures, simulated using the Slab-PTM patched Mpipi model. (a) The average protein density along the z-axis. (b-l) The time-resolved average protein density along the z-axis from simulation trajectories at temperatures ranging from 240 K to 340 K.

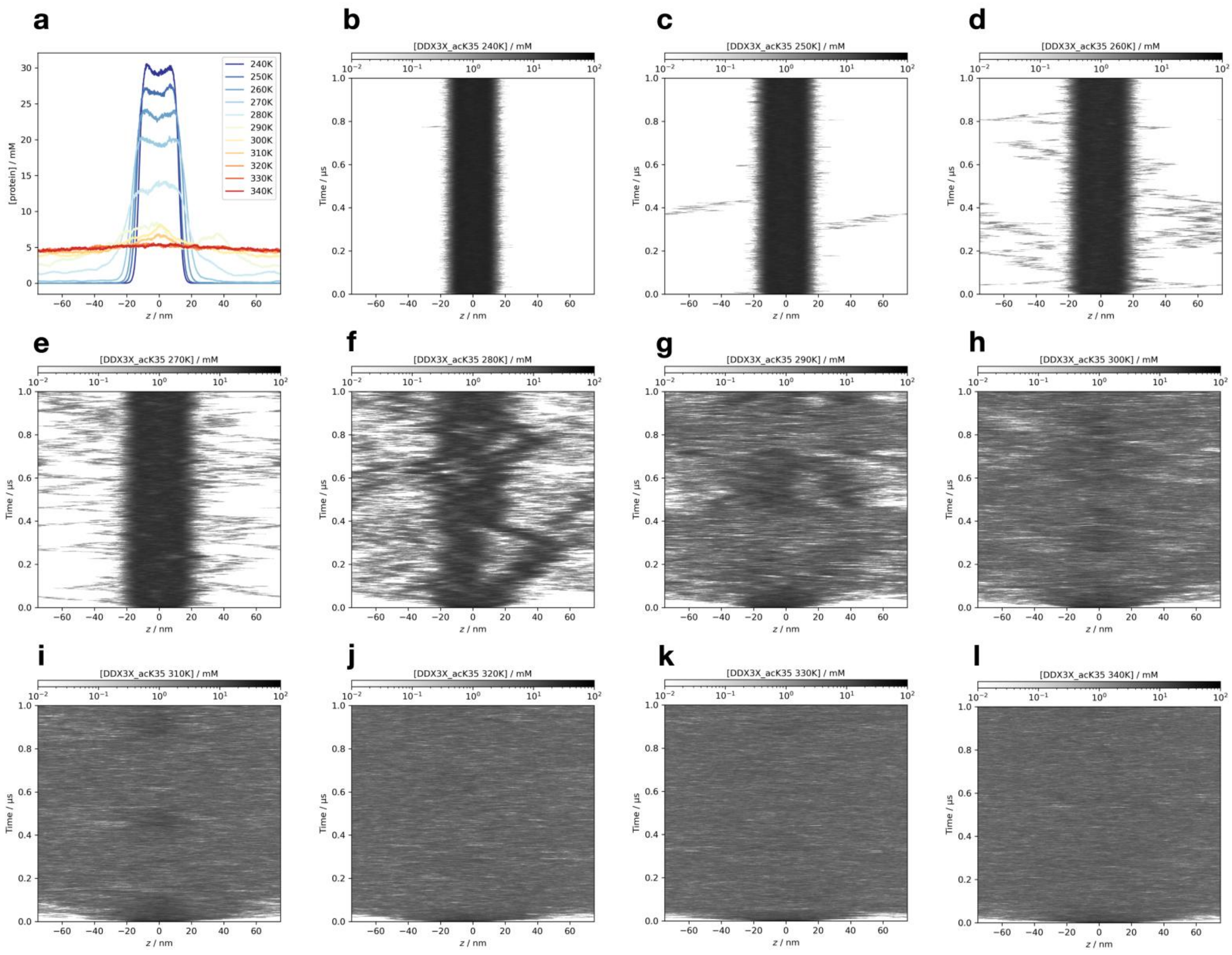

**Figure S47.** The detailed simulation results of the K35 acetylated DDX3X IDR1 domain at all temperatures, simulated using the Slab-PTM patched Mpipi model. (a) The average protein density along the z-axis. (b-l) The time-resolved average protein density along the z-axis from simulation trajectories at temperatures ranging from 240 K to 340 K.

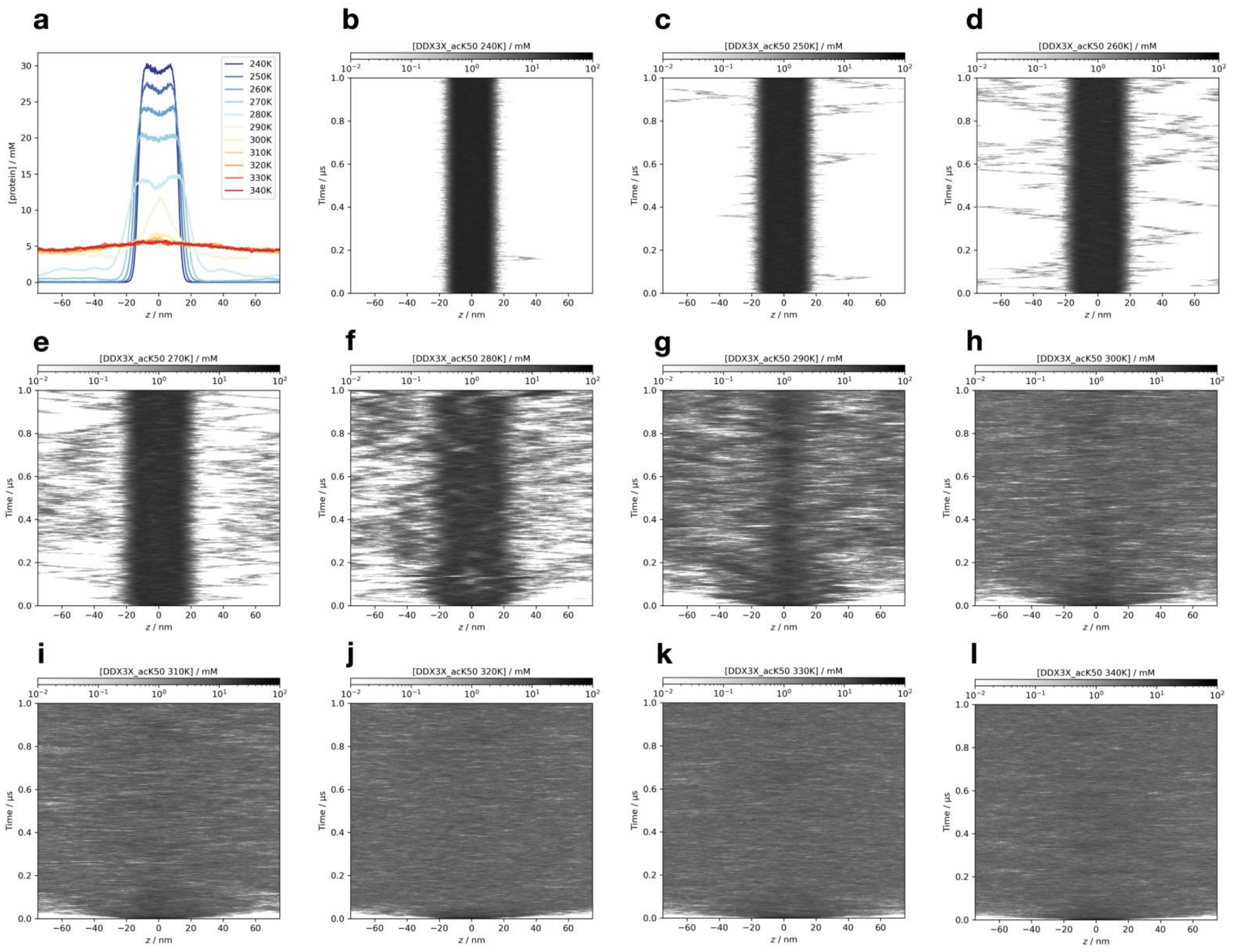


**Figure S48.** The detailed simulation results of the K50 acetylated DDX3X IDR1 domain at all temperatures, simulated using the Slab-PTM patched Mpipi model. (a) The average protein density along the z-axis. (b-l) The time-resolved average protein density along the z-axis from simulation trajectories at temperatures ranging from 240 K to 340 K.

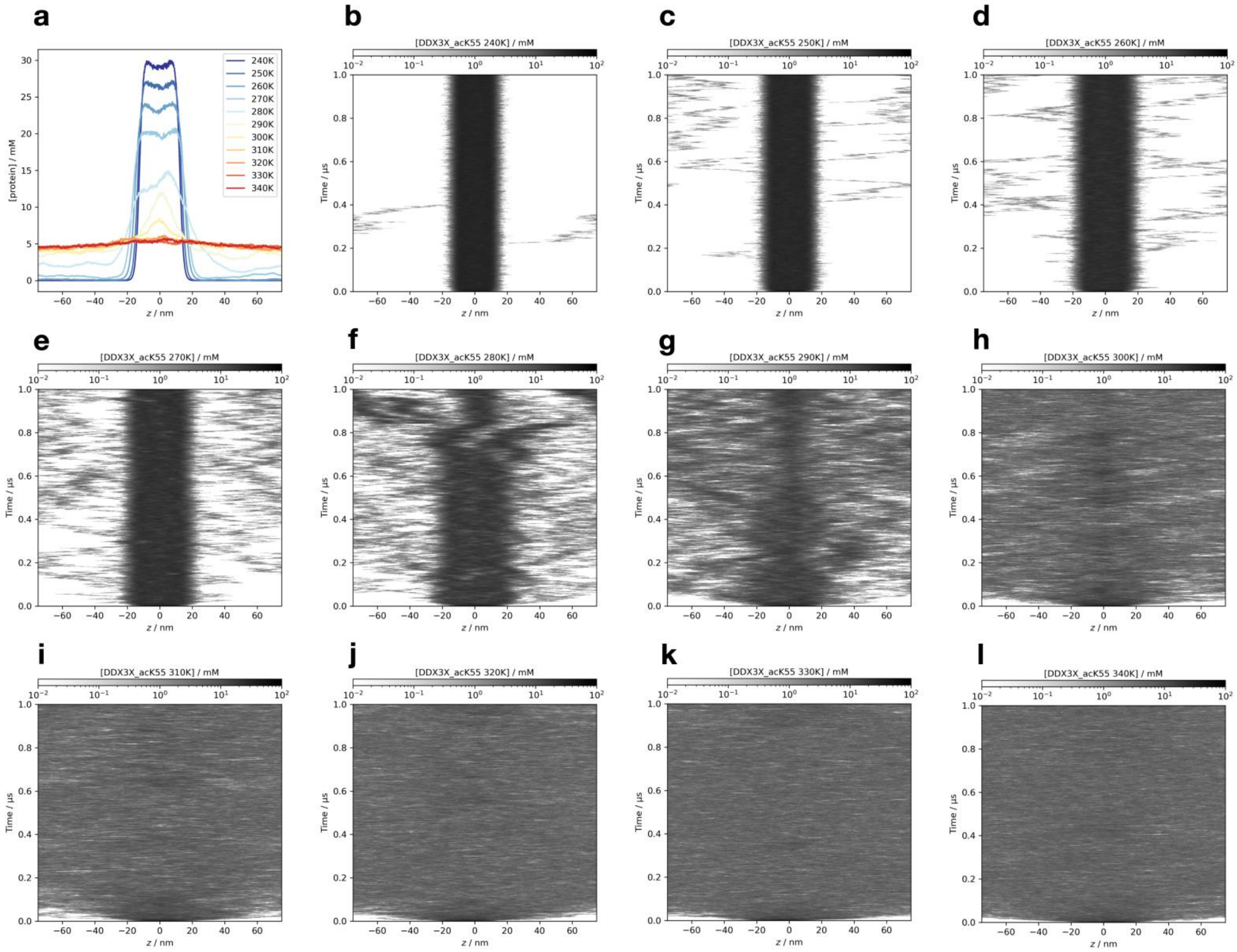


**Figure S49.** The detailed simulation results of the K55 acetylated DDX3X IDR1 domain at all temperatures, simulated using the Slab-PTM patched Mpipi model. (a) The average protein density along the z-axis. (b-l) The time-resolved average protein density along the z-axis from simulation trajectories at temperatures ranging from 240 K to 340 K.

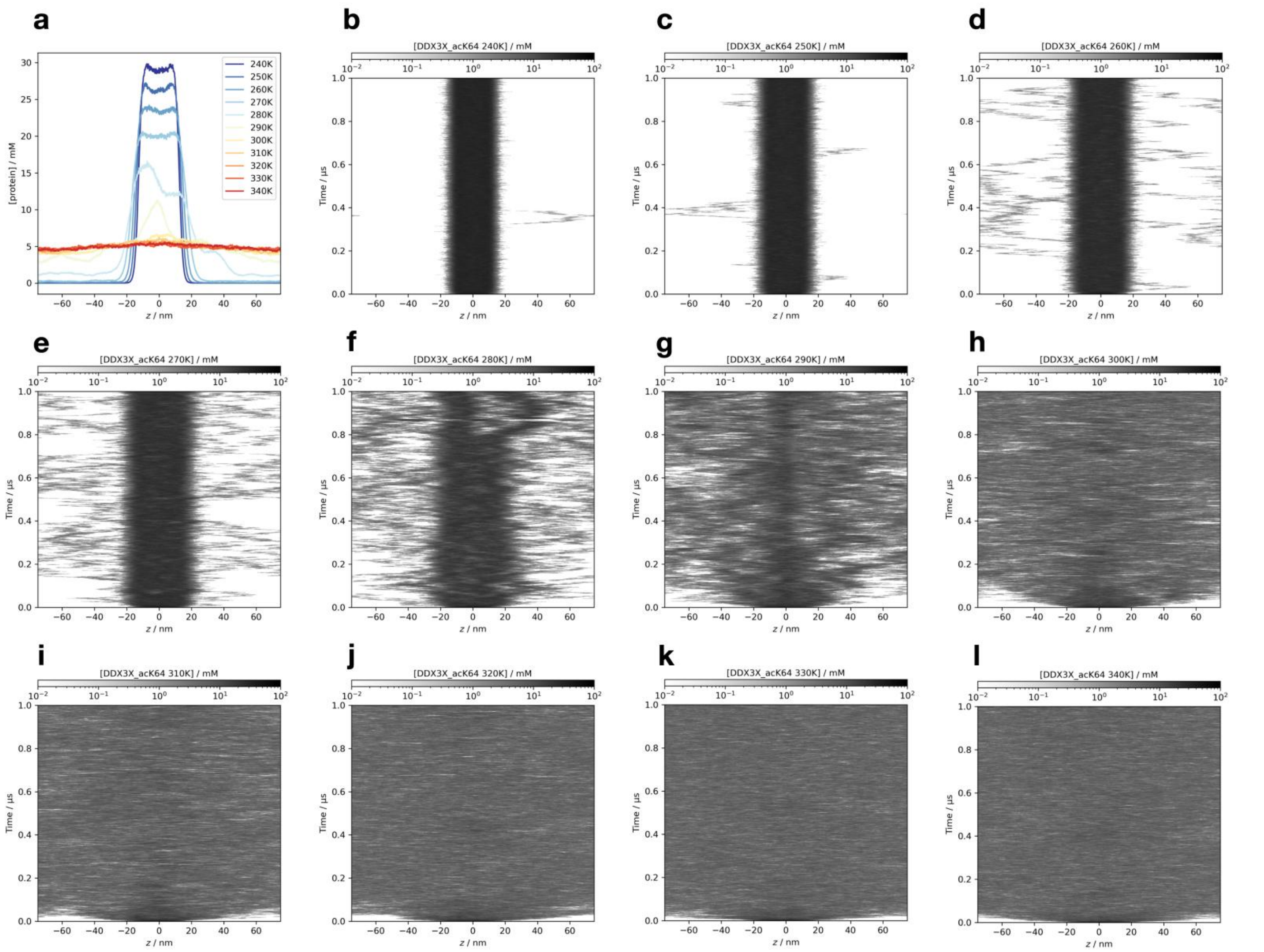


**Figure S50.** The detailed simulation results of the K64 acetylated DDX3X IDR1 domain at all temperatures, simulated using the Slab-PTM patched Mpipi model. (a) The average protein density along the z-axis. (b-l) The time-resolved average protein density along the z-axis from simulation trajectories at temperatures ranging from 240 K to 340 K.

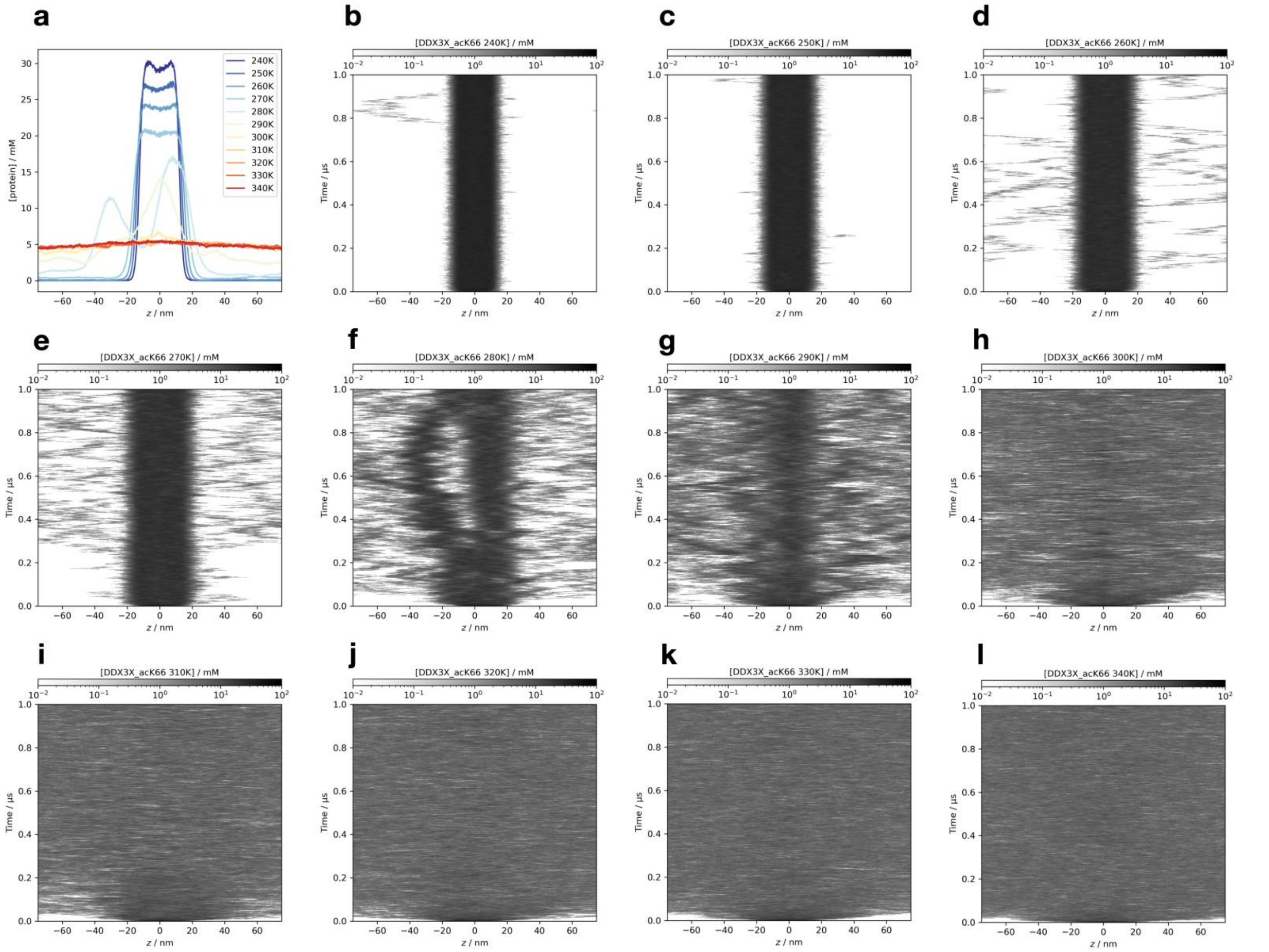


**Figure S51.** The detailed simulation results of the K66 acetylated DDX3X IDR1 domain at all temperatures, simulated using the Slab-PTM patched Mpipi model. (a) The average protein density along the z-axis. (b-l) The time-resolved average protein density along the z-axis from simulation trajectories at temperatures ranging from 240 K to 340 K.

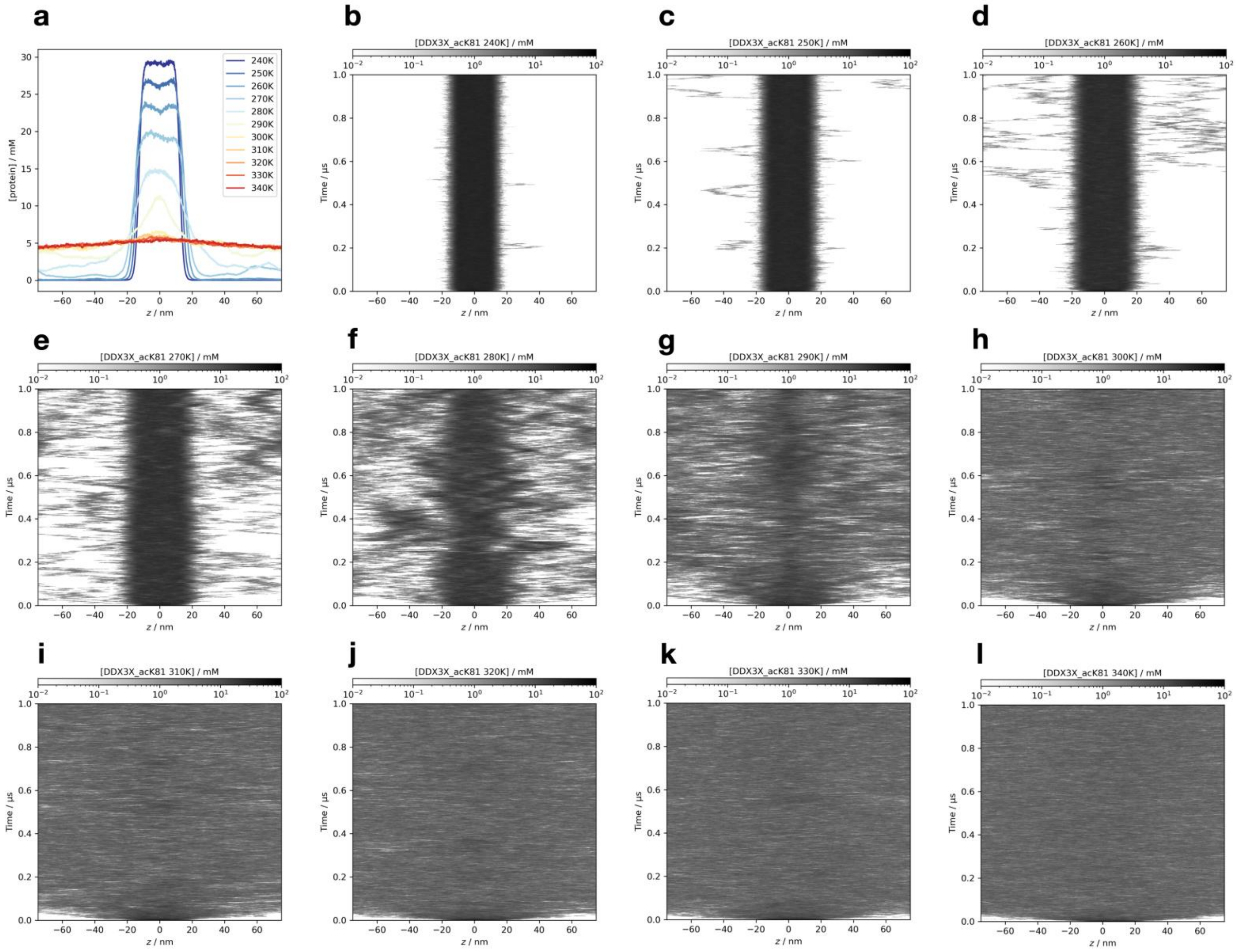


**Figure S52.** The detailed simulation results of the K81 acetylated DDX3X IDR1 domain at all temperatures, simulated using the Slab-PTM patched Mpipi model. (a) The average protein density along the z-axis. (b-l) The time-resolved average protein density along the z-axis from simulation trajectories at temperatures ranging from 240 K to 340 K.

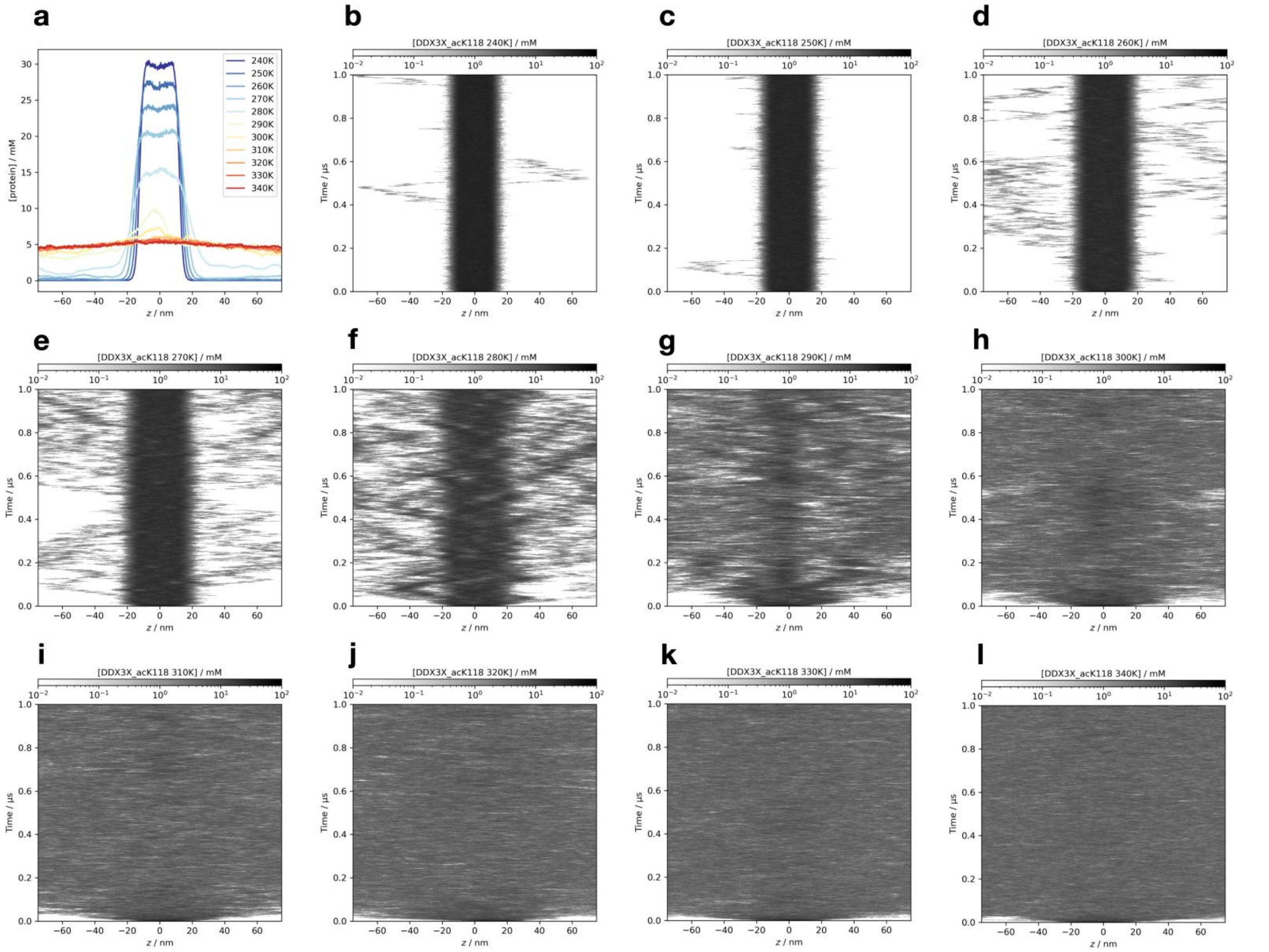


**Figure S53.** The detailed simulation results of the K118 acetylated DDX3X IDR1 domain at all temperatures, simulated using the Slab-PTM patched Mpipi model. (a) The average protein density along the z-axis. (b-l) The time-resolved average protein density along the z-axis from simulation trajectories at temperatures ranging from 240 K to 340 K.

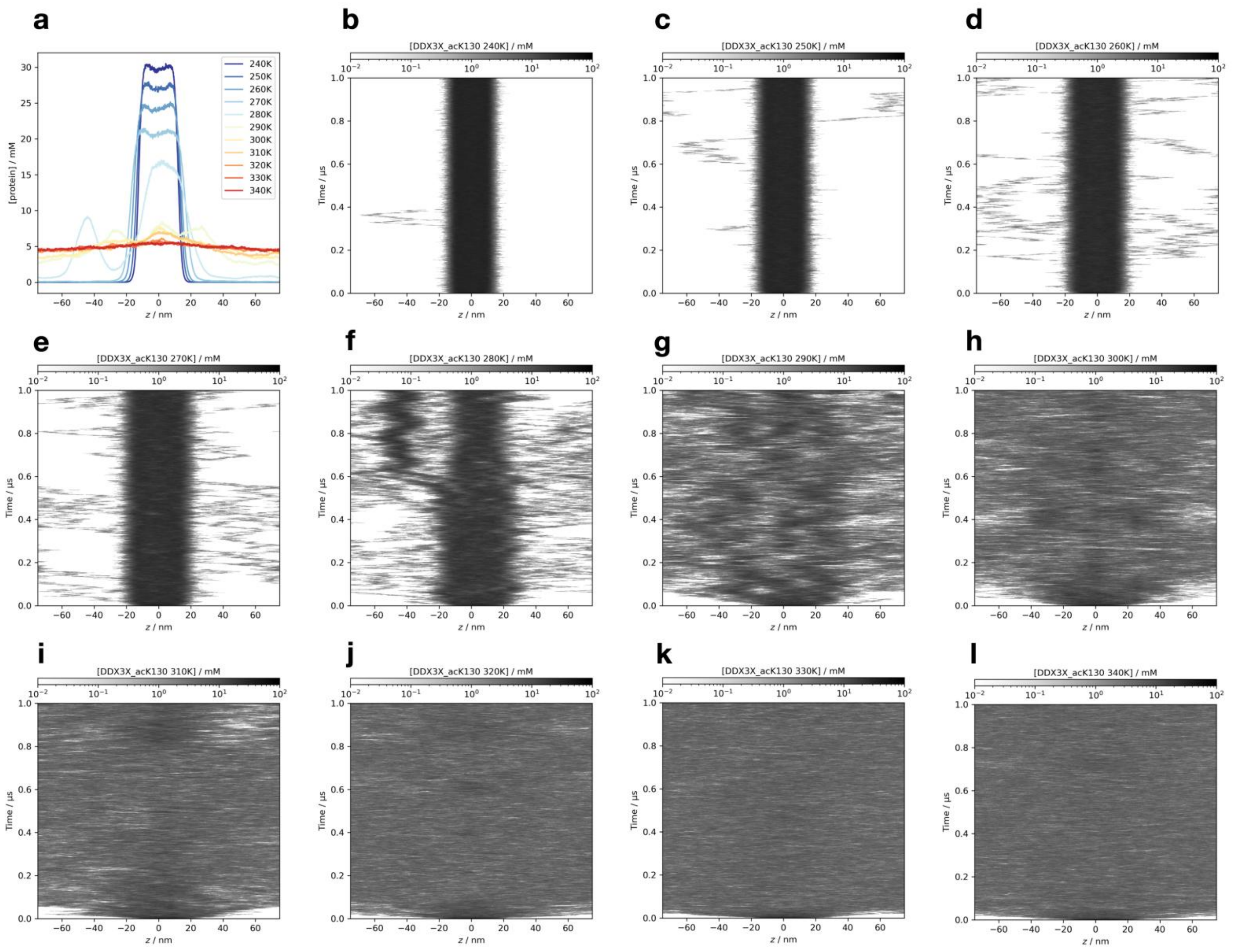


**Figure S54.** The detailed simulation results of the K130 acetylated DDX3X IDR1 domain at all temperatures, simulated using the Slab-PTM patched Mpipi model. (a) The average protein density along the z-axis. (b-l) The time-resolved average protein density along the z-axis from simulation trajectories at temperatures ranging from 240 K to 340 K.

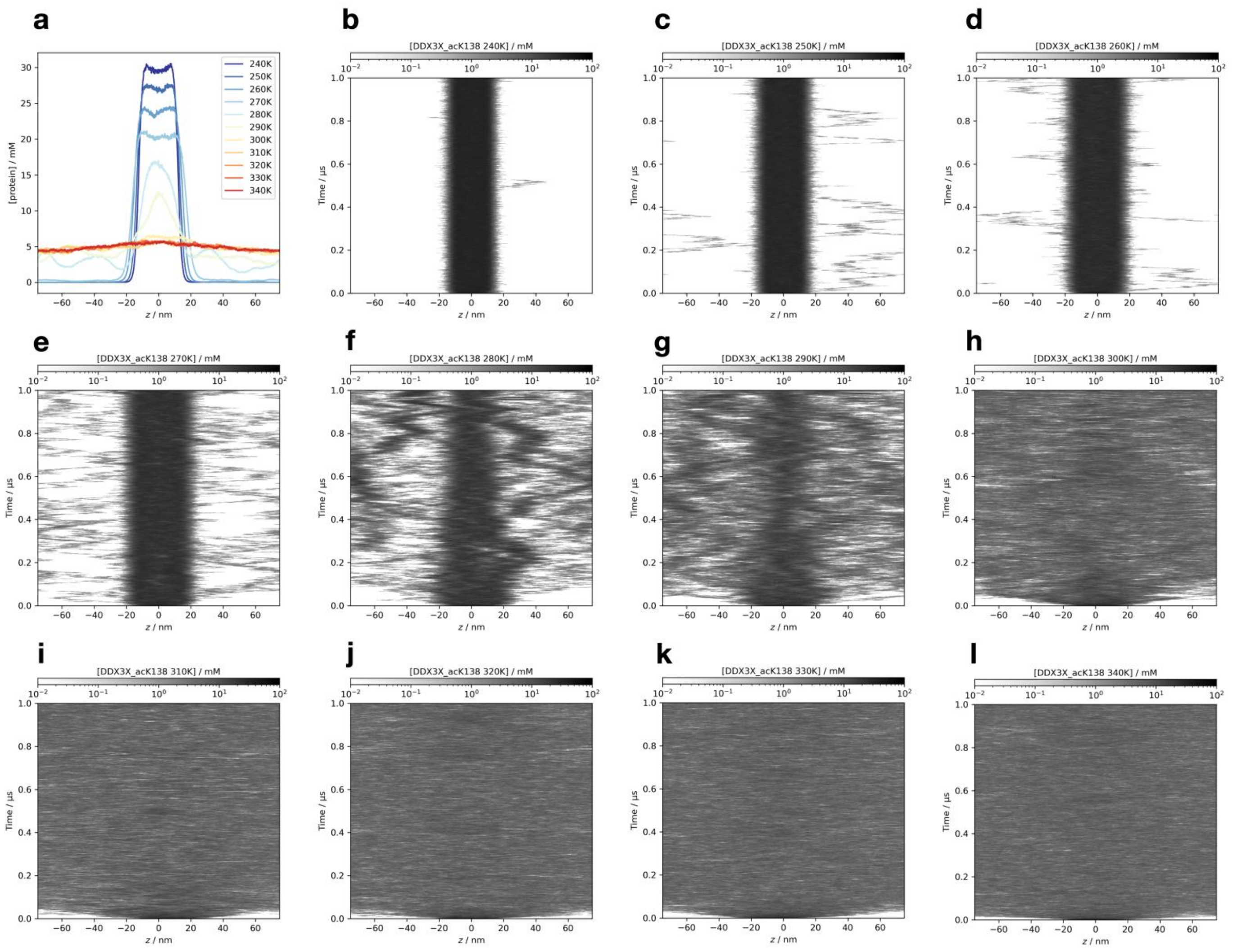

**Figure S55.** The detailed simulation results of the K138 acetylated DDX3X IDR1 domain at all temperatures, simulated using the Slab-PTM patched Mpipi model. (a) The average protein density along the z-axis. (b-l) The time-resolved average protein density along the z-axis from simulation trajectories at temperatures ranging from 240 K to 340 K.

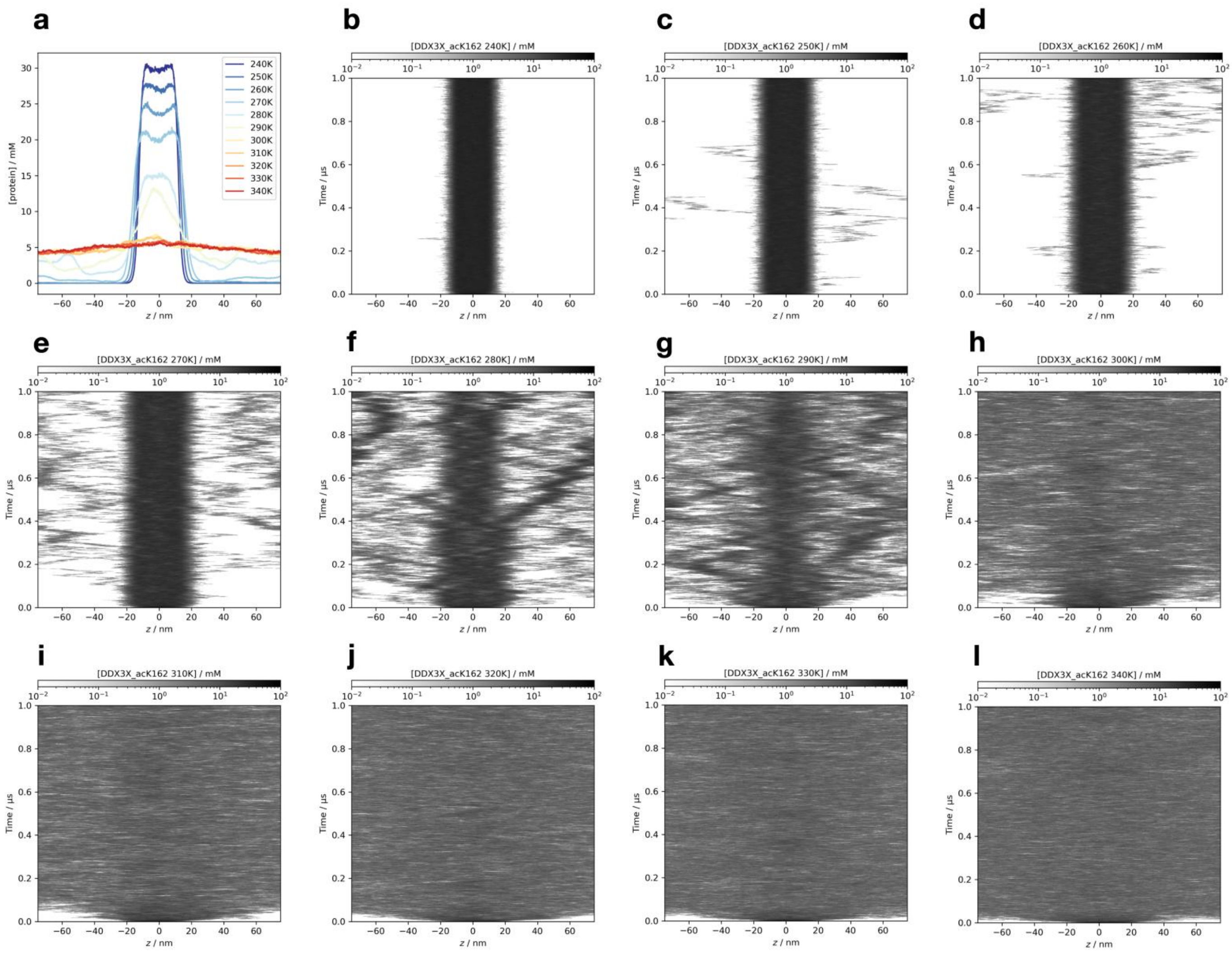

**Figure S56.** The detailed simulation results of the K162 acetylated DDX3X IDR1 domain at all temperatures, simulated using the Slab-PTM patched Mpipi model. (a) The average protein density along the z-axis. (b-l) The time-resolved average protein density along the z-axis from simulation trajectories at temperatures ranging from 240 K to 340 K.

| Ref Pair | PTM Pair | Ref Pair | PTM Pair | Ref Pair | PTM Pair | Ref Pair | PTM Pair | Ref Pair | PTM Pair |
|---|---|---|---|---|---|---|---|---|---|
| E-S | E-Sp | E-T | E-Tp | E-Y | E-Yp | E-Q | E-acK | E-R | E-Rm |
| D-S | D-Sp | D-T | D-Tp | D-Y | D-Yp | D-Q | D-acK | D-R | D-Rm |
| R-S | R-Sp | R-T | R-Tp | R-Y | R-Yp | R-Q | R-acK | R-R | R-Rm |
| K-S | K-Sp | K-T | K-Tp | K-Y | K-Yp | K-Q | K-acK | K-R | K-Rm |
| I-S | I-Sp | I-T | I-Tp | I-Y | I-Yp | I-Q | I-acK | I-R | I-Rm |
| V-S | V-Sp | V-T | V-Tp | V-Y | V-Yp | V-Q | V-acK | V-R | V-Rm |
| L-S | L-Sp | L-T | L-Tp | L-Y | L-Yp | L-Q | L-acK | L-R | L-Rm |
| T-S | T-Sp | T-T | T-Tp | T-Y | T-Yp | T-Q | T-acK | T-R | T-Rm |
| H-S | H-Sp | H-T | H-Tp | H-Y | H-Yp | H-Q | H-acK | H-R | H-Rm |
| M-S | M-Sp | M-T | M-Tp | M-Y | M-Yp | M-Q | M-acK | M-R | M-Rm |
| G-S | G-Sp | G-T | G-Tp | G-Y | G-Yp | G-Q | G-acK | G-R | G-Rm |
| A-S | A-Sp | A-T | A-Tp | A-Y | A-Yp | A-Q | A-acK | A-R | A-Rm |
| C-S | C-Sp | C-T | C-Tp | C-Y | C-Yp | C-Q | C-acK | C-R | C-Rm |
| S-S | S-Sp | S-T | S-Tp | S-Y | S-Yp | S-Q | S-acK | S-R | S-Rm |
| P-S | P-Sp | P-T | P-Tp | P-Y | P-Yp | P-Q | P-acK | P-R | P-Rm |
| N-S | N-Sp | N-T | N-Tp | N-Y | N-Yp | N-Q | N-acK | N-R | N-Rm |
| Q-S | Q-Sp | Q-T | Q-Tp | Q-Y | Q-Yp | Q-Q | Q-acK | Q-R | Q-Rm |
| F-S | F-Sp | F-T | F-Tp | F-Y | F-Yp | F-Q | F-acK | F-R | F-Rm |
| Y-S | Y-Sp | Y-T | Y-Tp | Y-Y | Y-Yp | Y-Q | Y-acK | Y-R | Y-Rm |
| W-S | W-Sp | W-T | W-Tp | W-Y | W-Yp | W-Q | W-acK | W-R | W-Rm |
| S-S | Sp-Sp | S-T | Sp-Tp | S-Y | Sp-Yp | S-Q | Sp-acK | S-R | Sp-Rm |
| T-S | Tp-Sp | T-T | Tp-Tp | T-Y | Tp-Yp | T-Q | Tp-acK | T-R | Tp-Rm |
| Y-S | Yp-Sp | Y-T | Yp-Tp | Y-Y | Yp-Yp | Y-Q | Yp-acK | Y-R | Yp-Rm |
| Q-S | acK-Sp | Q-T | acK-Tp | Q-Y | acK-Yp | Q-Q | acK-acK | Q-R | acK-Rm |
| R-S | Rm-Sp | R-T | Rm-Tp | R-Y | Rm-Yp | R-Q | Rm-acK | R-R | Rm-Rm |

**Table S1.** The comparison table of PTM amino acid dipeptide pairs and their reference dipeptide pairs.

| PTM Residue | pKa1 | pKa2 | Average charge at pH=7.0 |
|---|---|---|---|
| Phosphorylated serine (pSer) | 2.0 | 6.5 | ≈ -1.8 |
| Phosphorylated threonine (pThr) | 2.0 | 6.5 | ≈ -1.8 |
| Phosphorylated tyrosine (pTyr) | 2.0 | 6.0 | ≈ -2.0 |
| Asymmetric dimethylarginine (aDMA) | 12.5 | - | ≈ +1 |

**Table S2.** The pKa and charge values used for PTM residues in Slab-PTM.

| Protein Systems | pH | Ionic (mM) | Temperatures (K) | Ref |
|---|---|---|---|---|
| hnRNP A2 LCD | 5.5 | 0.025 | 240, 250, 260, 270, 280, 290, 300, 310, 320, 330, 340 | [1,2] |
| Phosphorylated hnRNP A2 LCD | 5.5 | 0.025 | | |
| Methylated hnRNP A2 LCD | 5.5 | 0.025 | | |
| TDP43 LCD | 7.0 | 0.15 | | [3] |
| Phosphorylated TDP43 LCD | 7.0 | 0.15 | | |
| FUS Nter | 7.0 | 0.15 | | [4] |
| Phosphorylated TDP43 LCD | 7.0 | 0.15 | | |
| NS2 | 7.4 | 0.1 | | [5] |
| Phosphorylated NS2 | 7.4 | 0.1 | | |
| DDX3X IDR1 | 7.5 | 0.125 | | [6] |
| Acetylated DDX3X IDR1 | 7.5 | 0.125 | | |
| DDX4 Nter | 7.0 | 0.1 | | [7] |
| Methylated DDX4 Nter | 7.0 | 0.1 | | |

**Table S3.** The simulation conditions used for benchmark, along with the corresponding experimental references.
